# Slow release of a synthetic auxin induces formation of adventitious roots in recalcitrant woody plants

**DOI:** 10.1101/2023.03.13.532257

**Authors:** Ohad Roth, Sela Yechezkel, Ori Serero, Avi Eliyahu, Inna Vints, Pan Tzeela, Alberto Carignano, Dorina P. Janacek, Verena Peters, Amit Kessel, Vikas Dwivedi, Mira Carmeli-Weissberg, Felix Shaya, Adi Faigenboim-Doron, Kien Lam Ung, Bjørn Panyella Pedersen, Joseph Riov, Eric Klavins, Corinna Dawid, Ulrich Z. Hammes, Nir Ben-Tal, Richard Napier, Einat Sadot, Roy Weinstain

**Affiliations:** School of Plant Sciences and Food Security, Faculty of Life Sciences, Tel Aviv University, Tel Aviv, Israel; The Institute of Plant Sciences, The Volcani Center, Ministry of Agriculture and Rural Development, Israel; The Robert H. Smith Institute of Plant Sciences and Genetics in Agriculture, The Robert H. Smith Faculty of Agriculture, Food and Environment, The Hebrew University of Jerusalem, Rehovot, Israel; Department of Electrical and Computer Engineering, University of Washington, Seattle, United States; Chair of Plant Systems Biology, Technical University of Munich, Freising, Germany; Chair of Food Chemistry and Molecular and Sensory Science, Technical University of Munich, Freising, Germany; Department of Biochemistry and Molecular Biology, School of Neurobiology, Biochemistry & Biophysics, Faculty of Life Sciences, Tel Aviv University, Tel Aviv, Israel; Department of Molecular Biology and Genetics, Aarhus University, Aarhus, Denmark; School of Life Sciences, University of Warwick, Coventry, UK

## Abstract

Clonal propagation of plants by induction of adventitious roots (ARs) from stem cuttings is a requisite step in breeding programs. A major barrier exists for propagating valuable plants that naturally have low capacity to form ARs. Due to the central role of auxin in organogenesis, indole-3-butyric acid (IBA) is often used as part of commercial rooting mixtures, yet many recalcitrant plants do not form ARs in response to this treatment. Here, we describe the synthesis and screening of a focused library of synthetic auxin conjugates in *Eucalyptus grandis* cuttings and identify 4-chlorophenoxyacetic acid-L-tryptophan-OMe as a competent enhancer of adventitious rooting in a number of recalcitrant woody plants, including apple and argan. Comprehensive metabolic and functional analyses reveal that this activity is engendered by prolonged auxin signaling due to initial fast uptake and slow release and clearance of the free auxin 4-chlorophenoxyacetic acid. This work highlights the utility of a slow-release strategy for bioactive compounds for more effective plant growth regulation.

## Main text

Adventitious roots (ARs) are defined as roots that regenerate from non-root tissues, in contrast to lateral roots (LR) that are post-embryonic roots formed from root tissue^1^. Clonal (vegetative) propagation of plants by induction of ARs from stem cuttings is a requisite step in selection and breeding programs as well as in routine agricultural practices and has tremendous economic importance^2^. Clonal propagation is also a cornerstone in forestry, the ornamental plant industry, and the development of elite rootstocks to provide resistance to pests, diseases and changing environmental conditions^2^. Despite its significant economic and agricultural importance, a major barrier still exists for propagating clones of many valuable plants that naturally have low or no capacity form ARs or that lose this ability during maturation^3–5^.

AR development is a heritable, quantitative genetic trait^6,7^ that shows high plasticity and is controlled by multiple intrinsic and environmental factors^8–10^. In particular, it was shown to be controlled by a complex network of plant hormones crosstalk, in which auxin signaling plays a central role in each step of the process^11–15^. In some plant species, lower endogenous indole 3-acetic acid (IAA) levels in difficult-to-root mature cuttings compared to easy-to-root juvenile ones, e.g., *Eucalyptus grandis* (*E. grandis*) and *Pisum sativum (P. sativum*), have been reported^16,17^, as well as absence of IAA maxima in the cambium zone of difficult-to-root pine cuttings^18^; the cambium being the tissue from which ARs typically form^19^ and where IAA maxima are often observed^20,21^. However, other plant species show comparable endogenous auxin levels in juvenile and mature shoots or even higher in the mature difficult-to-root ones^22,23^ yet the ability to form AR is significantly impaired in mature shoots, with or without exogenous auxin application. Thus, the accepted presumption to date is that auxin responsiveness (as derived from auxin metabolism, transport and perception) has changed in mature cuttings, not any more able to convey the correct signaling pathways to support AR formation. Indeed, stronger auxin response (*DR5:GUS*) was reported in young vs. mature cuttings of *P. sativum* upon similar exogenous auxin treatments^16^ and differential expression profiles of auxin-regulated genes were observed in easy-vs. difficult-to-root poplar^24,25^, pine^18,26,27^ and *Eucalyptus* species^17,28–31^ along AR induction.

Although IAA is the most prevalent endogenous auxin in plants, and the first to be used for induction of AR formation^32^, indole 3-butyric acid (IBA) and 1-naphthalaneacetic acid (NAA) have been found to be more efficient and for the past 60 years are the major components in most commercial rooting formulas^2,33^. Initially, the increased efficacy of IBA and NAA was attributed to their higher light-resistance, but more recent studies point to their differential metabolism and transport (compared to IAA) as the potential source for their efficacy^34–36^. Over the years, efforts have been made to increase the effectiveness of IBA by different approaches, including its conjugation to various molecules^37–40^. Nevertheless, many recalcitrant plants respond poorly to exogenous application of these compounds^41,42^, and their vegetative propagation remains a significant challenge.

The above observations have prompted us to hypothesize that synthetic auxins might represent an underexplored chemical space of bioactive compounds that could assist in overcoming the loss of rooting capability in difficult-to-root plants. Synthetic auxins constitute a large set of small organic molecules with structural resemblance to IAA and that mimic the effects of the endogenous IAA by promoting the interaction between the auxin receptors TRANSPORT INHIBITOR RESPONSE1 (TIR1)/AUXIN-SIGNALING F-BOX (AFB) and Aux/IAA^43^. Despite this central similarity, differences in metabolism^44^, transport^45,46^ and perception specificity^47–49^ have been observed between IAA and several synthetic auxins (and among themselves), that presumably lead to different expression profiles of auxin responsive genes and/or sets of auxin-related phenotypes^49,50^. A number of synthetic auxins have been previously shown to promote rooting^51^ (e.g. 2,4-dichlorophenoxyacetic acid (2,4-D) and 2,4,5-trichlorophenoxyacetic acid (2,4,5-T), however, with the exception of NAA, their high auxin activity limits their practical use due to high phytotoxicity, or promotion of callus instead of roots^52^. We envisioned that the inherent phytotoxicity and growth-inhibitory effect of synthetic auxins could be mitigated by their slow release *in planta*, maintaining a low yet functional level of the bioactive molecule over a prolonged time, thus opening the door to uses beyond their traditional role as herbicides^53^. Moreover, lengthy auxin treatments were reported to improve AR induction^54–57^, which could further enhance the effectiveness of a slow-release approach.

To test this hypothesis, we synthesized a rationally-designed, focused library of four synthetic auxins conjugated to different residues, under the presumption that the conjugates will be hydrolyzed *in planta* (either enzymatically or chemically) to release the parent synthetic auxin. The conjugates were evaluated on difficult-to-root cuttings obtained from mature parts of *E. grandis* trees (Fig. 1a). A leading compound was found to enhance basal regeneration rates by 2-3-fold when applied to cuttings from diverse woody species. The dynamics underlying the compound activity is described herein.

**Fig. 1:**
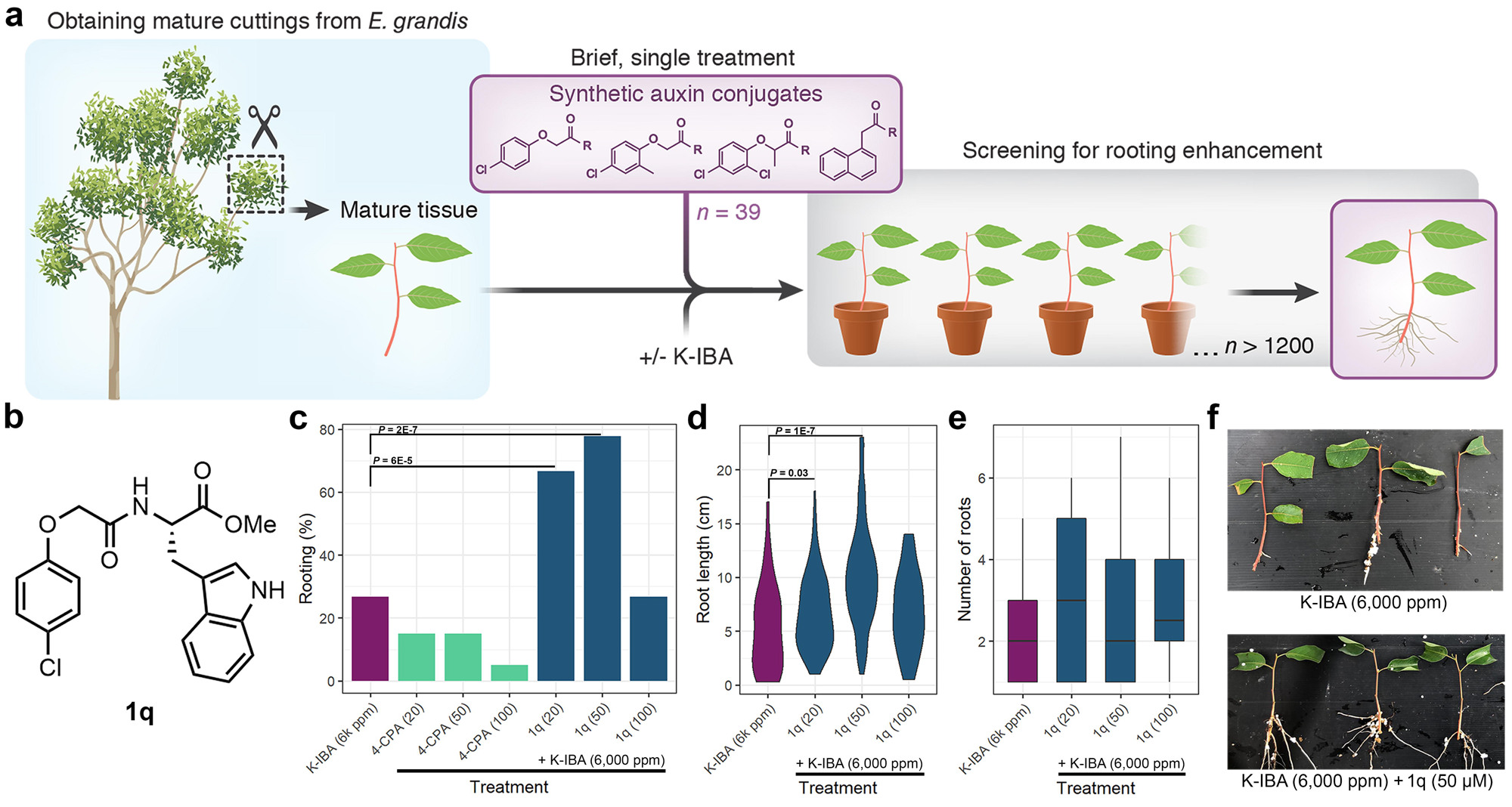
A chemical screen for rooting enhancers of difficult-to-root cuttings highlighted 4-CPA-Trp-OMe (1q). **a**, Illustration of the chemical screen. **b**, structure of **1q**; the most efficient compound. **c**, Rooting percentages 1 month after application. Fisher’s exact test p-values are presented for significantly better applications compared to K-IBA (6,000 ppm) as a single treatment (a one-sided test). n = 60, 20, 20, 20, 45, 45, and 45 cuttings per sample, respectively. Compound concentration (in µM) is shown in brackets. **d**, Distributions of root-length in regenerated cuttings. n = 37, 94, 98, and 35, respectively. Two-sided Mann-Whitney U test p-values are presented. **e**, Box-plot (center line, median; box limits, upper and lower quartiles; whiskers, 1.5x interquartile range) presenting number of roots per regenerated cutting. n = 17, 31, 36, and 12, respectively. None of the applications outperformed K-IBA significantly (Two-sided Mann-Whitney U test). **f**, Representative pictures of cuttings 35 days from the indicated treatment.

## Results

### Design and screening of synthetic auxins conjugates

To develop a suitable chemical library, the synthetic auxins 4-chlorophenoxyacetic acid (4-CPA) (**1**), 2-methyl-4-chlorophenoxyacetic acid (MCPA) (**2**), 2-(2,4-dichlorophenoxy) propionic acid (2-DP) (**3**), and NAA (**4**) were chosen for conjugation. The first three belong to the phenoxy acid family^58^ and feature a relatively strong, medium, and weak auxin activity, respectively, as determined by root elongation inhibition of *Prosopis juliflora*^59^. NAA belongs to the aromatic acetate family^60^ and is often used in commercial rooting enhancement mixtures^2^. Each of the synthetic auxins (**1**–**4**) was conjugated through its carboxylic acid, a required moiety for the hormone biological activity^61–63^, with a series of amine residues or methanol, forming a set of 39 conjugates (**1**–**4a**–**q,** Supplementary Fig. 1). The rooting enhancement capability of the conjugates and the free auxins (43 compounds in total) was evaluated using cuttings from mature *E. grandis* trees, which regenerate roots at low efficiency following 1 min submergence treatment with K-IBA, the potassium salt of IBA and the agricultural “gold standard” rooting enhancer^6^. The conjugates (100 μM) were applied by submerging the cutting base for 1 min or by spraying the cutting apical part, either as a standalone treatment or in combination with a 6,000 ppm K-IBA (24.9 mM) submergence treatment. The cuttings were then incubated in a rooting table for approximately 1 month before examination. In total, 20–90 cuttings were tested per conjugate-based treatment and ∼500 cuttings per K-IBA control treatment. At the chosen screening concentration (100 μM), none of the compounds outperformed K-IBA as a standalone treatment, however applications based on the combination of compounds **1a**, **1j**, **1o**, **1p** or **1q** with K-IBA showed significantly higher rooting rates (Supplementary Fig. 2). Of these compounds, **1q**, a conjugate of 4-CPA to L-tryptophan methyl ester (L-Trp-OMe, Fig. 1b), had the strongest effect, with nearly 40% root regeneration for either spray or submergence treatments when combined with K-IBA, compared to 17% for K-IBA alone (Supplementary Fig. 2). Of note, the corresponding free synthetic auxins at a similar concentration had no positive effect when combined with K-IBA. Likewise, increasing the amount of K-IBA applied as a single treatment from 6,000 up to 12,000 ppm did not improve rooting rates (Supplementary Fig. 3), and HPLC-MS/MS analysis shows similar IBA levels in cuttings 15 min after application of K-IBA or K-IBA+**1q** (Supplementary Fig. 4), ruling out mere increase in auxin levels or IBA uptake as underlying the effect observed when conjugates were added. Due to its hydrophobicity, applying higher concentrations of **1q** in a water-based solution was found to be challenging. As an alternative, we combined spray and submergence treatments, each at three different concentrations (20, 50 and 100 μM), in addition to 6,000 ppm K-IBA, in order to increase the applied concentration of **1q**. Strikingly, this dual application method resulted in AR induction efficiencies of 66% and 77% in response to **1q** at 20 and 50 μM, respectively (Fig. 1c), ∼3-folds higher than K-IBA alone. This effect was accompanied by the formation of comparable number, however significantly longer, roots per rooted cutting compared to K-IBA (Fig. 1d,e). To conclude, we find that a simple and short application of a synthetic auxin-based conjugate significantly augmented the saturated effect of K-IBA on de-novo root regeneration, which is a critical practice for the agricultural industry.

### Distinct bioavailability of 4-CPA underlies 1q activity

We speculated that **1q** exerts its bioactivity via a two-step process, in which **1q** is first hydrolyzed to its carboxylic acid form (**1r**), followed by removal of the amino acid that leads to release of bioactive 4-CPA (Fig. 2a). To rule out the possibility that **1q** itself can interact with the auxin perception machinery, and thus directly modulate AR formation, its ability to affect the TIR1-Aux/IAA7 auxin-perception complex formation was evaluated *in vitro* via surface plasmon resonance (SPR) measurements. The results show that neither **1q** nor **1r** have any measurable auxin or anti-auxin activity (Fig. 2b and Supplementary Fig. 5, respectively). Thus, the activity of **1q** seems to depend on its ability to release a bioactive 4-CPA. To understand the fate of **1q** *in planta*, cuttings of *E. grandis* were submerged and sprayed with either 4-CPA or **1q** (in addition to K-IBA submergence), and the small-molecules content of the cutting bases were analyzed periodically via HPLC-MS/MS for up to 8 days following treatment. Figure 2c shows the metabolic derivatives of **1q** following its application, and Figure 2d shows the levels of 4-CPA measured following **1q** or free 4-CPA application. The first time point, 1 h post-application, illuminates one of the features of **1q**; the esterification of the carboxylic acid leads to a more hydrophobic molecule (logD at pH = 7.0: 0.06 vs. 3.17), resulting in a 10-fold higher uptake of **1q** (Fig 2c) compared to free 4-CPA (Fig. 2d) (515.5 ± 24.4 vs. 53.2 ± 5.6 pg/mg fresh weight (FW)). This time point also demonstrates the rapid de-esterification of **1q** *in planta*, with ∼13% **1r** out of the measured **1q**-derived forms, and a negligible amount of 4-CPA, pointing to the amide bond cleavage as the rate-limiting step in 4-CPA release. Indeed, 6 h after application, **1q** levels decreased by ∼82% (to 90.9 ± 4.1 pg/mg FW) while comparable **1r** and 4-CPA levels were detected (48.5 ± 0.8 and 38.2 ± 1.1 pg/mg FW, respectively). This observation suggests that initially, a significant portion of **1q** is not available for immediate de-esterification. In the subsequent ∼48 h, **1q** level remained relatively constant whilst a clear conversion of **1r** to 4-CPA was detected. Interestingly, despite the higher uptake of **1q** compared to free 4-CPA, the maximal level of 4-CPA was comparable in both treatments (53.2 ± 4.0 and 72.0 ± 2.0 pg/mg FW for 4-CPA or **1q**, respectively) (Fig. 2d). However, the timing of their formation was strikingly different; while 4-CPA level peaked 1 h post-application for the free 4-CPA (Fig. 2d), it only peaked after 24 h for **1q** (Fig. 2c). In addition, clearance rates were very different; 4-CPA retained an approximate physiologically relevant level of an auxin (>10 pg/mg FW, as measured for IAA in *E. grandis* cuttings, Supplementary Fig. 6) for only 2 days when applied directly but persisted for >6 days when applied in the form of **1q** (Fig. 2c,d). The above observations suggest that **1q** application could support prolonged auxin signaling *in planta*. To further evaluate this point, we turned to *Arabidopsis thaliana* (*Arabidopsis*), first seeking to establish the activity of **1q** in this model plant and then to correlate it with auxin signaling. In line with the results in *E. grandis*, a brief (1.5 h) shoot application of **1q**, but not of 4-CPA or IBA (10 μM), resulted in a substantial increase in AR formation of intact etiolated *Arabidopsis* seedlings (Fig. 2e, Supplementary Fig. 7). In accord, applying the same treatment to *Arabidopsis DR5:Luciferase* line, encoding for a high turnover auxin reporter suitable for long-term imaging^64^, led to stronger and more prolonged auxin signaling in response to **1q** compared to 4-CPA (Fig. 2f). Importantly, these observations also demonstrate that K-IBA treatment is not necessarily a prerequisite for the activity of **1q**. Collectively, the results of the above experiments suggest that **1q** serves as a reservoir for continuous auxin release that promotes AR induction and development.

**Fig. 2:**
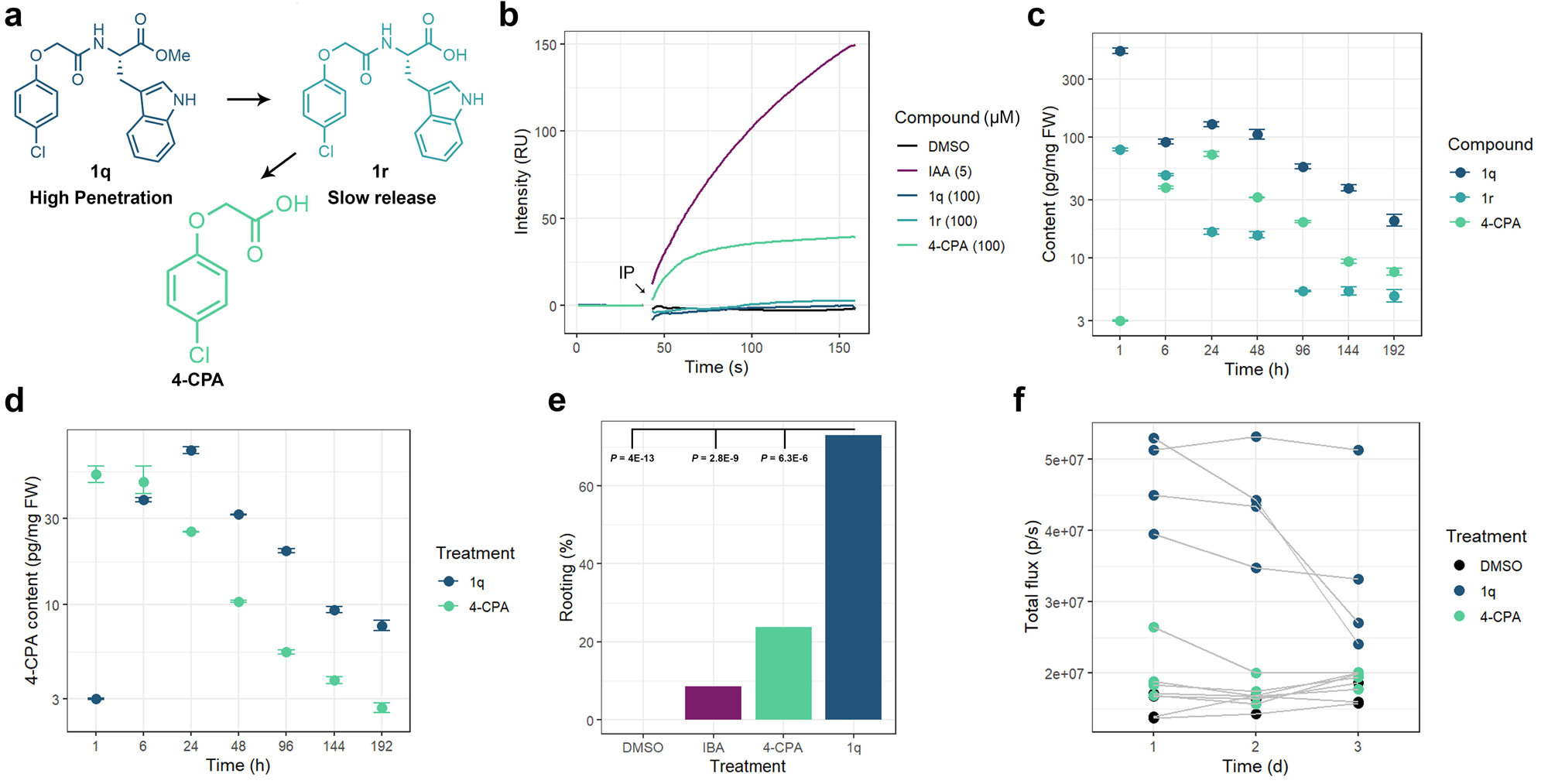
1q combines high penetration with slow 4-CPA release to facilitate prolonged auxin signaling. **a**, Schematic representation of 4-CPA release from **1q** *in planta.* **b**, SPR assay testing the intrinsic activity of the indicated compounds or DMSO (1%) *in-vitro* using TIR1 and IAA7 degron. IP stands for injection point of TIR1 mixed with the tested auxin in solution, RU stands for resonance unit. **c**, HPLC-MS/MS quantification of the compounds shown in **a** following application of **1q** (100 µM) + K-IBA (6,000 ppm). Each sample is composed of 3 replicates, extracted from a pool of 20 cutting-bases harvested together, and the means are presented in logarithmic scale. Error-bars represent standard errors. **d**, HPLC-MS/MS quantification of 4-CPA following **1q** (100 µM) + K-IBA (6,000 ppm) or 4-CPA (100 µM) + K-IBA (6,000 ppm). Each sample consists of 3 replicates except for 4-CPA after 1 h which consists of 2. Samples composition is as specified in **c**. Means are presented in logarithmic scale. Error-bars represent standard errors. **e,** Percentages of two-week-seedlings (*Arabidopsis*) that developed adventitious roots in response to the indicated treatments (10 µM for IBA, 4-CPA or **1q**, and 0.1% for DMSO applied specifically to shoots for 1.5 h via a split-dish). Shown are p-values of Fisher’s exact test testing the hypothesis that **1q** treatment results in higher rooting percentages (a one-sided test). n = 37,35, 38, and 37 respectively. The average AR number for rooted seedlings treated with **1q** was 4.48±0.65 (SE), n = 27. **f**, Time-lapse quantification of *DR5:luciferase* activity in *Arabidopsis* seedlings following the indicated treatments (10 µM for 4-CPA or **1q**, 0.1% for DMSO. Shoot specific 1.5 h application). Each dot represents ∼20 seedlings (pooled). *P =* 2.93E-05, repeated measures ANOVA.

### Evading IAA homeostasis regulators amplifies 4-CPA signaling

In addition to the characteristics of the conjugate, which engender higher uptake and slow auxin release, intrinsic properties of the released synthetic auxin might shape the cellular responses to **1q** and were therefore examined. SPR measurements showed that 4-CPA is a ∼2-orders of magnitude weaker binder of TIR1 than IAA (Fig. 2b). A comparable weaker auxin activity was found *in vivo*, using qualitative (lacZ-based, TIR1+Aux/IAA7, Supplementary Fig. 8a) and quantitative (degron-YFP based, TIR1+Aux/IAA9 and AFB2+Aux/IAA9) yeast-2-hybrid (Y2H) assays (Supplementary Fig. 8b). Initial weak auxin activity was also found in root growth inhibition and *DR5:Venus* response assays in *Arabidopsis* (Fig. 3a,b). Several synthetic auxins were shown to evoke unique expression profiles of auxin responsive genes compared to IAA^50^, which could underlie the AR promotion activity observed for **1q**. An extended analysis of 4-CPA binding performances by a systematic evaluation of 11 Aux/IAA and both TIR1 and AFB2 receptors (with the appropriate EC_50_ for each, calculated from the curves shown in Supplementary Fig. 8b), did not reveal a specific degradation pattern in response to 4-CPA (Fig. 3c and Supplementary Fig. 8c). Thus, based on the *Arabidopsis* auxin perception mechanism, a differential signaling response to 4-CPA as a result of unique binding is unlikely. Nevertheless, while *Arabidopsis* root growth recovers quickly from IAA inhibition, it is entirely arrested in response to 4-CPA (Fig. 3a), suggesting differences in transport and/or catabolism between the two molecules. A shoot-to-root movement assay in *Arabidopsis* implied that 4-CPA is a mobile auxin (Supplementary Fig. 9). However, although 4-CPA was found to utilize the native IAA importer AUXIN-RESISTANT1 (AUX1) (Fig. 3d,e), a solid-supported membrane (SSP)-based electrophysiology assay testing the transport activity of PIN-FORMED8 (PIN8), an adopted model for PINs activity^65^, demonstrated that unlike IAA (and the analogue 2,4-D^65^), 4-CPA did not induce a significant current response at the concentration tested (20 μM) (Fig. 3f). These observations suggest that 4-CPA is only partially subjected to the canonical polar auxin transport mechanism. Unlike 4-CPA, AUX1-expressing oocytes did not accumulate **1r** upon 30 min incubation (Supplementary Fig. 10a), and comparable levels of **1q** (and its derivative **1r**) were found in both AUX1-expressing and non-expressing oocytes (Supplementary Fig. 10b). To evaluate the contribution of 4-CPA movement to AR formation following **1q** treatment, we examined the *aux/lax* quadruple mutant^66^, and found it insensitive to AR induction by **1q** (brief shoot application, Supplementary Fig. 11). Together, these experiments suggest that cell-to-cell movement of 4-CPA, but not of its precursors, is crucial for effective AR induction in response to a brief treatment of **1q**. To address the hypothesis that 4-CPA differs from IAA not only in transport but also in catabolism, we adopted the *gh3* octuple mutant, in which IAA inactivation via conjugation to amino acids is deficient^67^. The activity of enzymes from this family was recently shown to be the first step in auxin catabolism^68^. By measuring root growth after 6 days of treatment with IAA or 4-CPA at 10 nM (conditions showing similar effect on growth of Col-0 roots, Fig. 3a,g) we found the *gh3* plants to be hyper-sensitive to IAA, but not to 4-CPA (Fig. 3g). These results are in line with previous conjugation rates measured for 2,4-D vs. IAA^44^, and favors the assumption that 4-CPA is not an efficient substrate for the main IAA-inactivation pathway in *Arabidopsis*. Collectively, this body of evidence suggests that 4-CPA weak binding to the auxin receptors is compensated by enhanced cellular persistence. Thus, the prolonged auxin signaling following **1q** application is achieved not only due to the slow release of 4-CPA, but also as a consequence of 4-CPA bypassing key auxin homeostasis regulators.

**Fig. 3:**
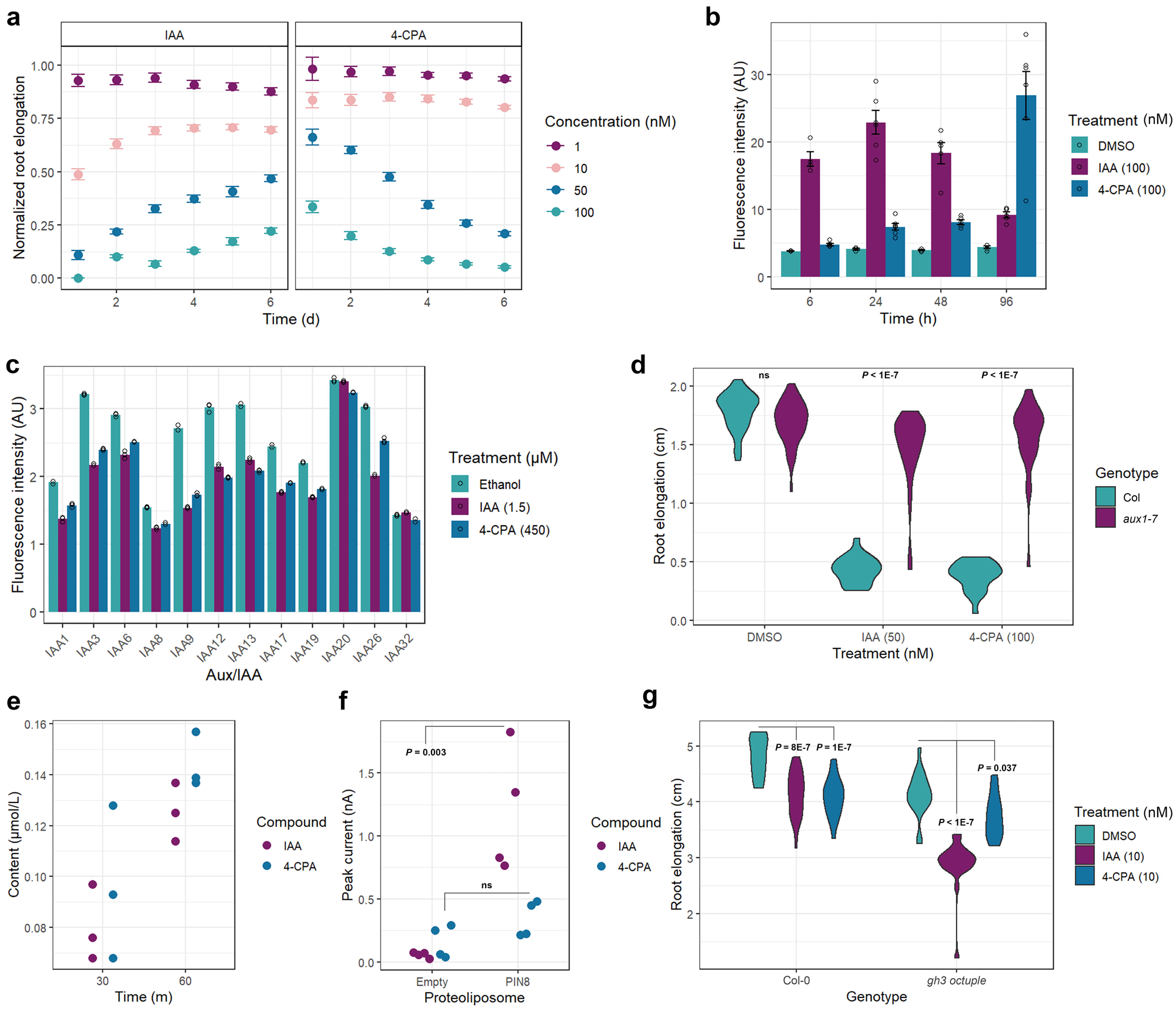
Bypassing key auxin homeostasis regulators supports 4-CPA long-term signaling. **a**, Means of normalized root elongation of seedlings incubated with IAA or 4-CPA at the indicated concentrations (0.1% DMSO as control), n > 18 plants per sample. Error-bars represent standard errors. **b**, Activity of the auxin reporter *DR5:Venus* in roots. n > 4 root tips per sample. Shown are means of fluorescence intensity. Error-bars represent standard errors **c**, Quantitative Y2H assay using TIR1 and YFP-tagged Aux/IAAs and the indicated auxin or ethanol (0.1%) as mock n = 3 biological replicates. Shown are means of fluorescence intensity. Error-bars represent standard errors. **d**, Root elongation of *aux1-7* seedlings in response to 3 days treatment. n > 40 for auxins, n > 20 for DMSO (0.1%). Two-sided Tukey’s HSD p-values are presented, ns stands for not significant. **e**, HPLC-MS/MS quantification of the indicated auxin in oocyte cells expressing AUX1 transporter. n = 3 biological replicates. **f**, Solid-supported membrane (SSM)-based electrophysiology assay with empty or PIN8-containing proteoliposomes, n = 4 biological replicates. Two-sided Student’s t test p-values are presented. **g**, Root elongation of *gh3* octuple mutant plants in response to a 3-day treatment with the indicated compounds. n > 18 plants per sample. Two-sided Tukey’s HSD p-values are presented.

### 4-CPA release is enzymatically regulated in plants

The rapid de-esterification of **1q** in *E. grandis* implies that 4-CPA release rate is largely determined by its amide bond hydrolysis (Fig. 2c). To investigate the mechanism of this step, we synthesized 4-CPA conjugated to D-Trp-OMe, forming the enantiomer of **1q** (**1s**, Fig. 4a). Since enantiomers possess similar chemical and physical properties, differences in their hydrolysis rate (or activity) *in planta* could be attributed to enzymatic regulation. HPLC-MS/MS analysis of apical and basal parts of *E. grandis* cuttings 24 h after application of **1q** or **1s** showed that 4-CPA accumulates only in response to **1q** treatment (Fig. 4a). Furthermore, in a root elongation assay using *Arabidopsis* seedlings, **1q** was found to engender ∼100-fold stronger inhibitory response than **1s** (Supplementary Fig. 12). From these results, a major role for enzymatic cleavage in 4-CPA release can be inferred. Members of the metallopeptidases family; IAA-Leu-RESISTANT1 (ILR1)/ILR1-like (ILLs) are known to hydrolase amides of indole-based compounds^69–72^, raising the possibility of a similar amido-hydrolase activity towards **1q** and/or **1r**. To test this hypothesis, we adopted the *Arabidopsis* triple mutant *ilr1 ill2 iar3*, which shows a compromised response to a range of IAA-amino acid conjugates^68,73^. The *ilr1 ill2 iar3* triple mutant was insensitive to **1q** in root elongation (continuous incubation, measured after 3 days, Fig. 4b) and in AR induction in etiolated seedlings (brief shoot application, Fig. 4c, Supplementary Fig. 13) assays. To validate these results, the appropriate GST-recombinant *Arabidopsis* enzymes were tested *in vitro* for their activity against **1q** and **1r**, or against IAA-alanine (IAA-Ala), an established substrate^71^ serving to verify the enzymes activity in the assay. While all three enzymes hydrolyzed IAA-Ala (Supplementary Fig. 14), only ILR1 and ILL2 efficiently hydrolyzed **1r**, and none hydrolyzed the parent **1q** (Fig. 4d). Of note, a marginal but detectable activity of ILR1 and ILL2 was also detected against the D-enantiomer of **1r** (**1t**, Supplementary Fig. 14d), which might explain the minor bioactivity observed for its parent compound **1s** in *Arabidopsis* (Supplementary Fig. 12). In an attempt to better understand their specificities, we turned to the three-dimensional structures of the three enzymes, using the available X-ray crystal structure of ILL2^74^, and AlphaFold^75^ predictions for ILR1 and IAR3. We found the ligand-binding pockets of the two active enzymes, ILR1 and ILL2, to contain a deep hydrophobic niche, in contrast to the pocket of IAR3, which is elongated, shallow and contains a smaller hydrophobic patch (Supplementary Fig. 15). In agreement, molecular docking calculations (Glide, Schrödinger, 2021-4) positioned the non-polar indole of **1r** inside the deep hydrophobic niche of the active enzymes, while in the non-active IAR3, neither the indole nor the phenoxy group formed sufficient non-polar interactions with the catalytic pocket (Fig. 4d). Further correlating the ligand-pocket non-polar interactions to substantial enzymatic activity, docking analysis positioned the indole group of IAA-Ala inside the IAR3 pocket, in close interaction with the hydrophobic patch (Supplementary Fig. 16). Having established that ILR1 and ILL2 are responsible for the hydrolysis of **1r**, we nevertheless observed a residual root growth inhibition for *ilr1 ill2 iar3* in response to longer incubation durations with **1q** (Supplementary Fig. 17), implying participation of additional amidohydrolase (Ah). We speculated that other ILL enzymes might underlie this effect, and generated two quintuple mutant lines; *ilr1 ill2 iar3 ill3 ill5* and *ilr1 ill2 iar3 ill1 ill6* (termed quintuple *3,5* or *1,6* respectively) using CRISPR-Cas9 (Supplementary Fig. 18–21). The *quintuple 3,5* was only slightly less sensitive to a 7-day incubation with **1q** (0.5 μM) compared to the triple mutant, while the *quintuple 1,6* was entirely resistant (Supplementary Fig. 17a). Structural modelling of the four enzymes (ILL1,3,5, and 6) revealed differences in the hydrophobicity and geometry of their ligand binding sites, with ILL1 and ILL6 binding sites being more hydrophobic than those of ILL3 and ILL5 (Supplementary Fig. 17b). Collectively, we established that the second, rate-limiting, step in 4-CPA release is enzymatically regulated, and that members of the ILR1/ILLs family are the major enzymes cleaving **1r** to release 4-CPA *in planta*.

**Fig. 4:**
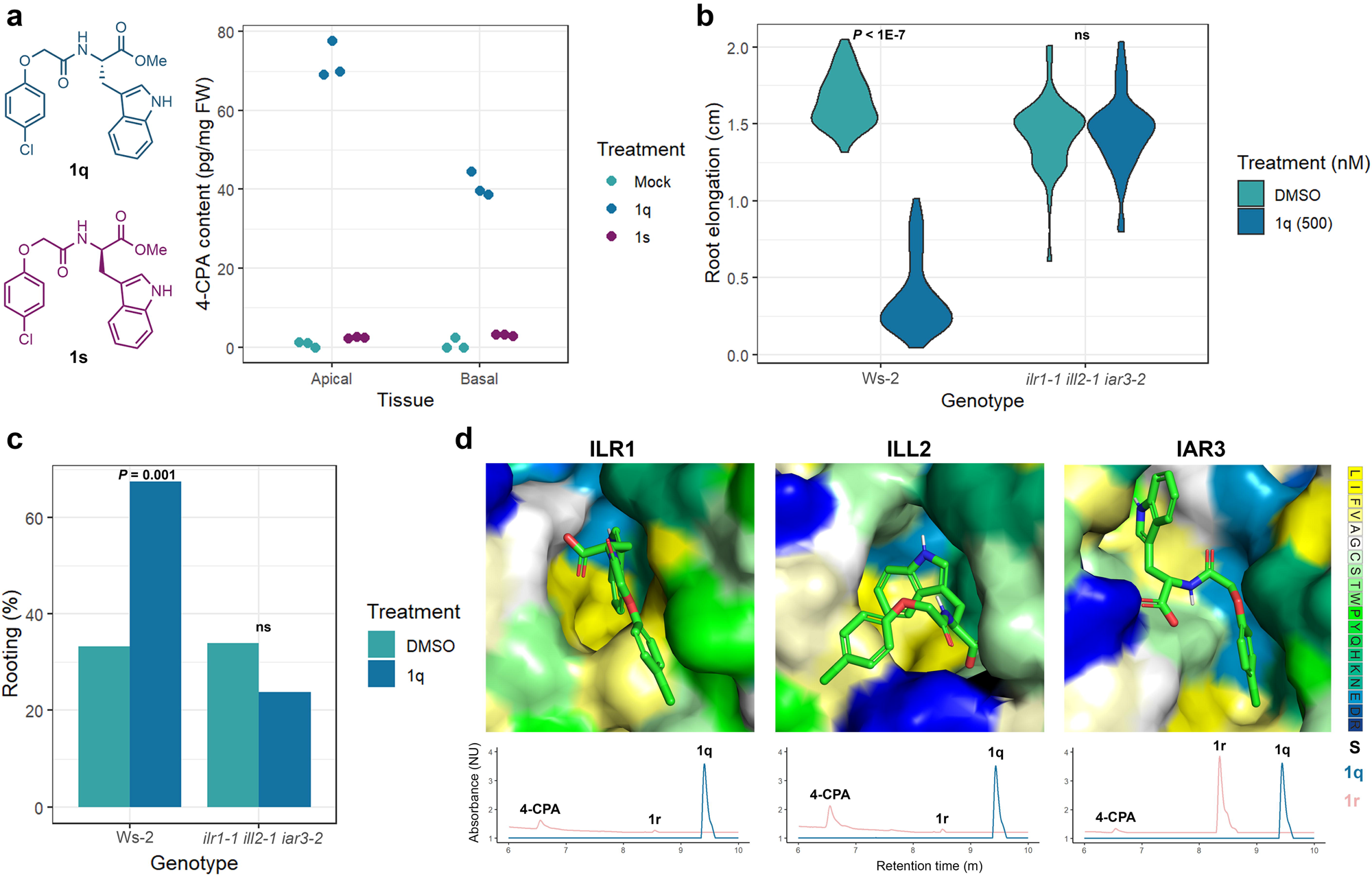
1r hydrolysis to 4-CPA is enzymatically regulated in *E. grandis* and *Arabidopsis.* **a**, Structures of the enantiomers **1q** and **1s** (*left*) and HPLC-MS/MS quantification of 4-CPA 24 h after apical or basal application with the indicated enantiomer (100 µM) + K-IBA basal treatment (6,000 ppm), or K-IBA (6,000 ppm) as mock (*right*). **b**, Root elongation of *ilr1-1 ill2-1 iar3-2* triple mutant in response to a 3-day treatment with **1q** or DMSO (0.1%). n > 47 plants per sample. Two-sided Tukey’s HSD p-values are presented, ns stands for not significant. **c**, Percentages of seedlings developed adventitious roots in response to **1q** treatment (10 µM for **1q**, 0.1% for DMSO. Shoot specific 1.5 h application in a split-dish), n > 43 plants per sample. P-value of Fisher’s exact test testing the hypothesis that **1q** treatment results with higher rooting percentages is shown (a one-sided test). ns stands for not significant. **d**, *Top:* Docking calculations of **1r** with ILR1, ILL2 and IAR3. Amino acids are color-coded according to the Kessel/Ben-Tal hydrophobicity scale^90^ (ranging from most hydrophobic (yellow) to most hydrophilic (blue)). *Bottom: In-vitro* hydrolytic activity of the enzymes towards **1q** (blue) or **1r** (pink) as monitored by HPLC-MS/MS. Absorbance was normalized by dividing each point with the minimal value in the according dataset. For clearer presentation, normalized units of the reactions with **1r** were multiplied by 1.1. **1q** is not cleaved by any of the enzymes while **1r** is cleaved by ILR1 and ILL2 (but not IAR3) to release free 4-CPA.

### Structural conservation of ILR1s supports 4-CPA release

Identifying specific members of the ILR1/ILLs family as the main activators of **1q** *in planta* opened the door to rationalizing and predicting its activation in other difficult-to-root cultivars. To this end, we performed a phylogenetic analysis based on 301 ILR1/ILLs proteins from 43 seed-plants that suggested two sub-trees (Fig. 5a). The two super-families are composed of two (AhA1-A2) and three (AhB1-3) distinct groups, with members of *Arabidopsis* occupying the AhA1 (ILL3), AhA2 (ILR1), AhB1 (ILL6), and AhB2 (ILL1, ILL2, IAR3 and ILL5) groups (Fig. 5a). We first sought to determine if activation of **1q** is functionally conserved between *Arabidopsis* and *E. grandis*. The *E. grandis* genome contains 11 *ILR1/ILLs* genes, of which we suggest only 9 to be active; based on proteins sequence-length and transcriptome of manually enriched vascular-cambium tissue (Fig. 5b, Supplementary Fig. 22). We focused on family AhA2 due to its high-confidence topology compared to AhB2 (Fig. 5a and alternative tree in Supplementary Fig. 23), and since the single *E. grandis* protein in the AhB1 group is apparently a pseudogene (*Eucgr.F03795*; expression not detected, and short putative protein sequence of 290 amino acids, Fig. 5b). Of the 3 active AhA2 genes, *EgK02589* (the suggested direct ortholog of ILR1; Fig. 5a) and *EgK02598* (which is clustered at the other orthologous group of Ah2A) were found to be highly expressed in vascular-cambium obtained 24 h after K-IBA treatment (Fig. 5b). The two genes were separately over-expressed in the *Arabidopsis ilr1 ill2 iar3* triple mutant background and their enzymatic activity was inferred from a root-growth complementation assay in the presence of **1q**. Interestingly, while lines overexpressing *EgK02589* restored the sensitivity to **1q** in a root-growth inhibition assay, lines expressing *EgK02598* did not (Fig. 5c). In agreement, structural modeling and docking calculations found the ligand binding pocket of EgK02598 flatter than those of EgK02589 and ILR1, and less favorable for the indole or phenoxy groups of **1r** to form significant non-polar and van der Waals interactions (Fig. 5d). These observations promote the hypothesis that EgK02589 contributes to the hydrolysis of **1q** to release active 4-CPA in the cambium. To broaden this observation, we similarly tested the activity of orthologs of ILR1/EgK02589 from *Populus trichocarpa* (Potri.006G207400, Pt400) and *Prunus persica* (Prupe.7G100000, Pp000), and one ortholog of EgK02598 from *Populus trichocarpa* (Potri.016G074100, Pt100). Again, only Pt400 and Pp000 but not Pt100 restored the response to **1q** (Fig. 5c), a trend that was further supported by structural modeling and docking calculations (Fig. 5d).

**Fig. 5:**
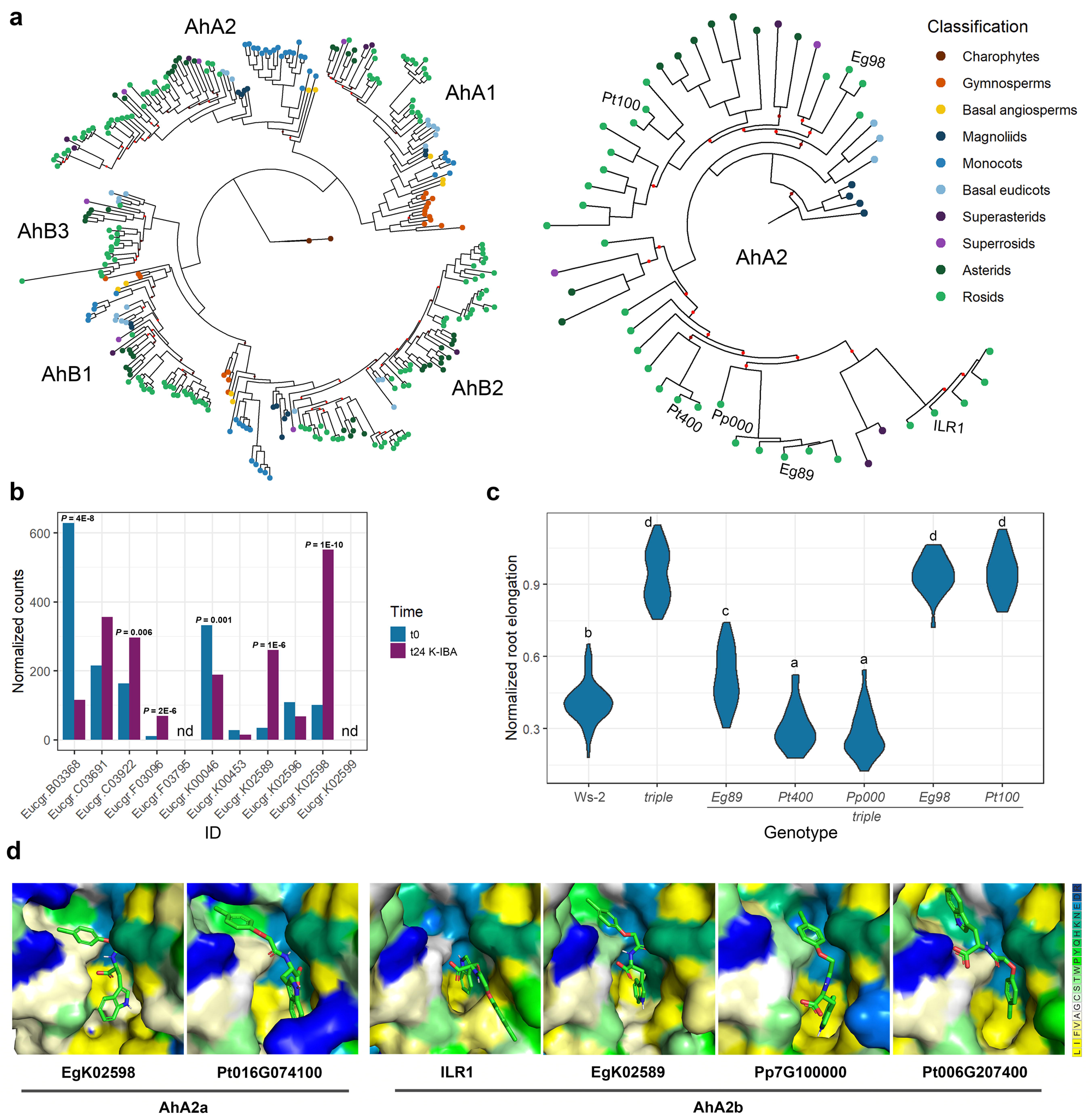
Structural conservation of ILR1 ligand binding pocket contributes to 4-CPA release. **a**, *Left:* Phylogenetic analysis of ILR1/ILLs family in seed-plants. Sequences from the charophyte algae *Klebsormidium nitens* were used as an outgroup. *Right:* Sub-tree presenting the phylogeny of core angiosperm-sequences in family AhA2. Annotated are the characterized enzymes; EgK02598 (Eg98), Potri.016G074100 (Pt100), Potri.006G207400 (Pt400), Prupe.7G100000 (Pp000), EgK02589 (Eg89), and ILR1. Branches are annotated in brown or red for bootstrap values lower than 85 or 70, respectively. **b**, Expression profile (shown as normalized counts according to DESeq2) of *E. grandis* ILR1/ILLs in samples enriched for cambium tissue of mature cuttings, either immediately after collecting the tissue (t0; blue) or 24 h after K-IBA (6,000 ppm) submergence (t24 K-IBA; pink). *Padj* (calculated by DESeq2) is presented. nd stands for not detected. **c**, Normalized root elongation in response to 4-day treatment with **1q** (300 nM) or DMSO (0.1%) as mock. n = 78 or 72 for Ws-2 or *ilr1-1 ill2-1 iar3-2* (*triple*), respectively, and >25 for over-expression lines. At least 10 T_2_ lines for each transgene were characterized, and data was collected from single, homozygous lines. Lower-case letters indicate significant groups based on two-sided Tukey’s HSD test (all significant comparisons are based on p-values smaller than 3E-7). **d**, Docking modeling of **1r** with the indicated enzymes. Amino acids are color-coded according to residues hydrophobicity, ranging from most hydrophobic (yellow) to most hydrophilic (blue)).

Collectively, the above experiments provide evidences that structural conservation of the ligand binding pocket among members of the ILR orthologous group supports **1r** cleavage, demonstrating the potential of structural modeling and docking calculations to predict the activation of **1q** in various plant species.

### 1q enhances ARs formation of distantly related woody species

The experimental evidences for enhanced de-novo root regeneration following **1q** application, together with the conservation of its key activating enzymes in diverse plant species, inspired us to examine the utility of **1q** in alleviating the barrier to rooting of agriculturally and environmentally-important difficult-to-root cultivars. We first examined *Eucalyptus x trabutii*; a very difficult to propogate hybrid of *E. camaldulensis* and *E. botryoides,* that has a relatively high resistance to cold^76^ and is important in supporting honeybee nutrition during the Israeli autumn^77^. For this hybrid, a combined application of **1q** and K-IBA dramatically outpreformed K-IBA alone in rooting efficency (45% vs. none, Fig. 6a). Likewise, for the apple (*Malus domestica*) rootstock clone CG41, which supports high yields, dwarfism and resistance to soil diseases but is considered difficult-to-root^78–80^, supplementation of **1q** increased rooting rate by ∼2-fold compared to K-IBA alone (Fig. 6b). As part of our efforts to support local cultivation of the argan tree (*Argania spinosa*); a species known for its tolerance to extreme environmental conditions and for its valuable oil, we evaluated several clones: three difficult-to-root clones (C124, C127, and ARS7^81^), of which the first two were directly obtained from the first trees that were planted in Israel as part of a botanical garden in 1931, and an easy-to-root clone, ARS1^81^. Application of **1q** doubled the rooting rates of cuttings from the >90-year-old C127 plant material but did not increase the low rooting efficiencies of C124 (Fig. 6c). For ARS7, again, **1q** doubled the basal root formation response to K-IBA, while for the more permissive ARS1, maximal response was found in both treatments (Fig. 6c). The success of the combined K-IBA + **1q** treatment enabled us to generate several plantations of selected elite clones of argan for further analyzing yield and profitability under different soil and climate conditions around the country (Supplementary Fig. 24). Altogether, these results suggest that woody, mature cuttings, for which poor regeneration rates are attributed to low auxin sensitivity, the saturated effect of IBA can be increased by low levels of **1q** (μM range). The results further suggest that ectopic addition of IBA is not necessarily a prerequisite for the rooting enhancement response to **1q** in mature woody tissues. Indeed, the rooting rates of *Populus alba* cuttings were doubled following application of **1q** as a standalone treatment (Fig. 6d). In line with the hypothesis of a conserved enzymatic hydrolysis being responsible for 4-CPA release, the basal parts of cuttings from *Eucalyptus x trabutii*, the ARS7 argan line, and *Populus alba* accumulated 4-CPA dominantly following **1q** application but not of its enantiomer **1s** (Supplementary Fig. 25).

**Fig. 6:**
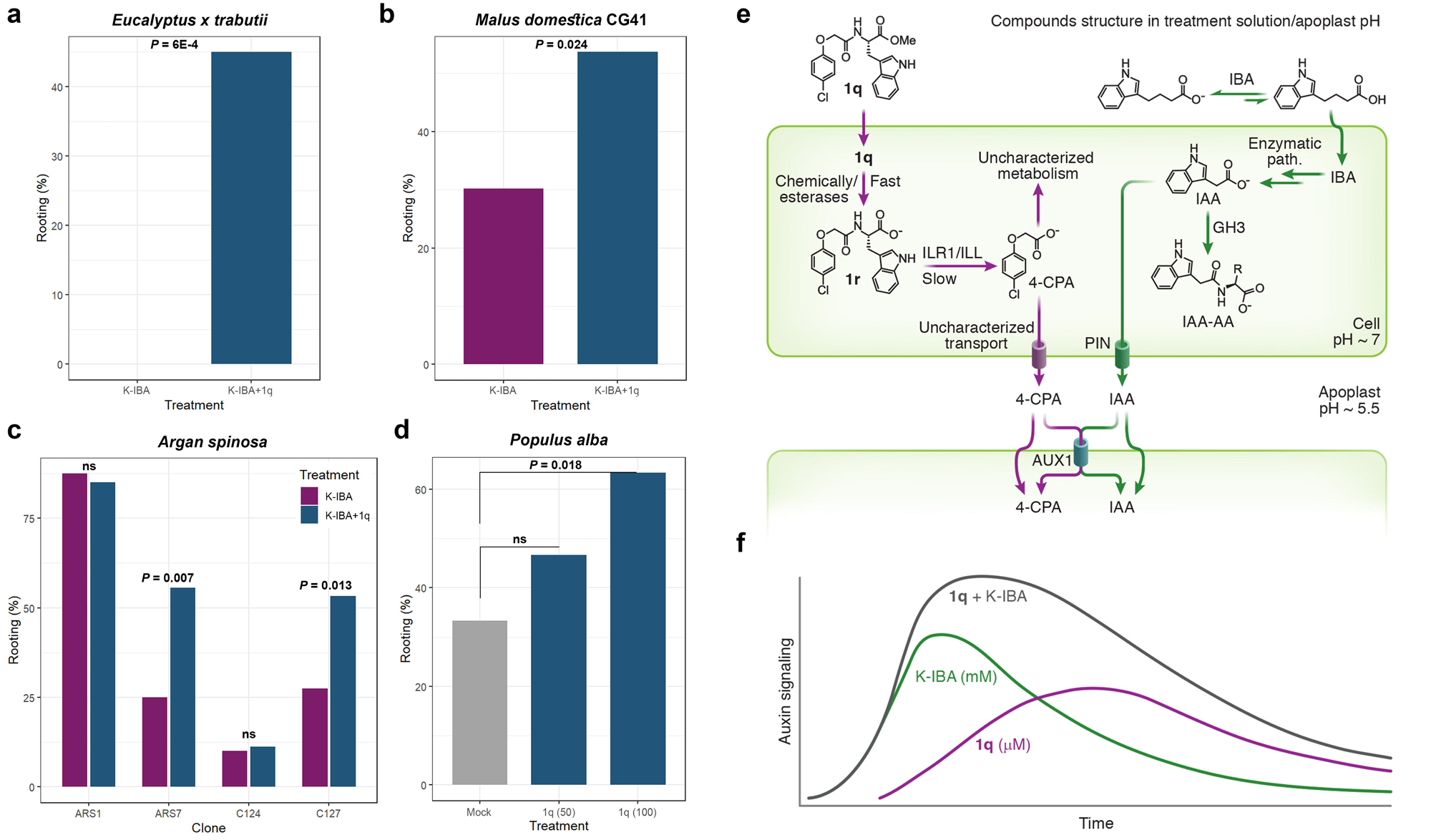
1q is a robust rooting enhancer for woody cuttings. **a**–**c**, Rooting percentages of mature cuttings obtained from the indicated species and clones applied with the dual treatment of K-IBA (6,000 ppm) + **1q** (50 µM). n = 20 (**a**), 41 and 43 (**b**), >36 (**c**) cuttings per sample. P-values of Fisher’s exact test testing the hypothesis that the dual treatment results with higher rooting percentages than K-IBA alone are shown (a one-sided test). ns stands for not significant. **d**, Rooting percentages of mature cuttings obtained from *Populus alba* treated with **1q** as a single agent at the indicated concentrations (shown in brackets as µM). n = 30 cuttings per sample. P-value of Fisher’s exact test testing the hypothesis that the **1q** treatment results with higher rooting percentages compared to mock (0.1% DMSO) is shown (a one-sided test). ns stands for not significant. The duration of all experiments was 1–2 months. **e-f**, A model comparing the fate of 1q and K-IBA following their application to woody cuttings. Schematic illustration of **1q** (purple) and K-IBA (green) fate when applied to woody cuttings (**e**) and their ensuing auxin signaling (**f**). Although K-IBA is applied at a very high concentration, its mostly negative charge under physiological conditions limits its accessibility to the plant, and later to cells. Inside the cell, IBA is converted into IAA, a strong yet highly regulated auxin. We suggest that these are the main factors underlying the auxin signaling pattern following a short K-IBA treatment (**f**). In contrast, **1q** is hydrophobic, limiting its concentration in a water-based solution, yet enabling enhanced tissue and cellular uptake. We suggest that efficient auxin delivery by **1q** is also a result of two distinct hydrolytic steps, responsible for a graduate, slow 4-CPA release. Moreover, 4-CPA is largely resistant to auxin feedback regulation, such as conjugation and transport, thus facilitating prolonged auxin signaling compared to IAA (**f**).

## Discussion

A model of the dynamics and metabolic fate of **1q** *in planta* is presented in Figure 6e. We suggest that following application, **1q** efficiently penetrates into the plant tissues and then into cells due to its neutral charge at a physiological pH and overall hydrophobicity. In the cells, the ester bond is quickly hydrolyzed (either chemically or by abundant cellular esterases) forming **1r**, which is mostly ionized in the cellular pH and therefore trapped inside the cell in the absence of efficient active transport^82^. Alternatively, **1q** could be hydrolyzed extracellularly. This scenario, however, is less likely considering that the highly acidic **1r** (predicted p*K*_a_ 3.3) is mostly ionized in the apoplast pH, which will result in low cellular accessibility^82^. In agreement, in long-exposure root elongation assays that mitigate differences in the uptake of small molecules, *Arabidopsis* roots were found to be more sensitive to **1q** than to **1r** (Supplementary Fig. 12). Subsequently, **1r** is cleaved by members of the ILR1/ILLs family, which presumably reside in the endoplasmic reticulum and in the cytosol^70,83^, to release 4-CPA intracellularly. Thus, although the measurements in *E. grandis* cutting bases detected comparable maxima levels of 4-CPA following **1q** or free 4-CPA treatments (Fig. 2c,d), we suggest higher intracellular accumulation of 4-CPA in response to **1q**. Practically, the dynamics and metabolic fate of **1q** *in planta* translate into a slow-release mechanism of a bioactive auxin inside the cells.

While the immediate auxin signaling elicited by 4-CPA is weaker than the one evoked in response to a native auxin, the higher cellular stability of 4-CPA supports an amplified and more sustained signaling over time. We provide biochemical evidence that 4-CPA is not a favorable substrate to the PIN8 transporter, and genetic evidence for its low affinity to the IAA conjugating enzymes GH3s^84^. Since transport and conjugation are two of the main feedback responses to auxin^85,86^, we suggest that evading these homeostasis regulators further contributes to prolonged auxin signaling. We also provide data suggesting that 4-CPA is able to move basipetally, and that AUX1 is required for its rooting enhancement effect. This in turn suggests that 4-CPA is a subject of an uncharacterized efflux transporter, which may support the compound basipetal movement, presumably through the phloem bulk flow, following apical application. Whether apical response to 4-CPA contributes indirectly to AR formation remains an open question. Identification of 4-CPA homeostasis regulators (e.g. metabolism, transport) will provide a better understanding of the contribution of these processes to the activity of **1q**. The delivery model we describe offers an advantage over the traditional application of free auxins (e.g., IBA, NAA, etc.) that are typically mostly ionized in the apoplast and may require active transport for efficient uptake. Together, our observations suggest that the short dual application of IBA and **1q** enables a fast and strong auxin response (K-IBA applied at mM concentrations) followed by a prolonged and sustaining signaling (**1q** applied at μM) (Fig. 6f).

Using a two-phases of auxin treatment, Ludwig-Muller et al. were able to distinguish between induction of callus proliferation and AR establishment^54^. In analogy to our system, a higher-resolution understanding of how the two compounds interact (temporally and spatially) during AR induction and development is of significant mechanistic and practical interest, and may assist in optimizing future applications. Moreover, the flexible molecular design of a synthetic auxin conjugate can be further fine-tuned by modulating either the synthetic auxin or its conjugated amino acid to provide a palette of auxin responses varying in strength and duration that could be tailored to different plant species and even specific clones.

One apparent limitation of **1q** is that easy-to-root trees do not necessarily benefit from its application, which might even be inhibitory, as can be seen in the example of the argan clone ARS1 (Fig. 6c). This could be a result of an already high level of auxin in the cuttings. Nevertheless, easy-to-root trees may benefit from treatment with **1q** when rooting is performed outside of the optimal season^87^. The enzyme-dependency of auxin release means that the efficacy of **1q** in different trees might be attenuated by the expression level of these enzyme and could require an ad hoc optimization of the applied concentration. Finally, the application method explored herein relies on dipping of cuttings in a solution containing an organic solvent to impart solubility. Although this is a commonly practiced method^2,55^, further formulation of **1q** to industry standards^88^ (e.g. talc powder, water dispersible granules or an emulsifiable concentrate) can be leveraged to eliminate the need for an organic solvent or to allow for alternative application methods. In any case, it would important to retain the hydrophobic nature of **1q** to maintain its efficient uptake into tissues.

In conclusion, the ability of **1q** to significantly improve rooting rates in cuttings from multiple species of commercially-relevant, difficult-to-root trees together with an industry-compatible application method, and given the widespread use of rooting enhancers in many agriculture, horticulture and forestry sectors, suggest a high level of commercial readiness. In addition, the slow-release approach as applied herein can be incorporated into other agricultural practices in which auxin is applied beyond as an herbicide, such as modulating root system complexity, controlling fruit growth or the timing of fruits set^89^, to allow for more optimized responses.

## Supporting information

Source data main text

## Acknowledgments

We thank the undergraduate students Ariel Verblun and Geffen Yehezkely in R.W. lab for supporting experiments. We also thank Eilon Shani for providing *DR5:Luciferase, DR5:Venus,* and *aux1-7 Arabidopsis* lines, Malcolm Bennett and Ranjan Swarup for providing the *aux/lax* quadruple mutant, Mark Estelle for providing Y2H vectors, Karin Ljung for providing the *gh3* octuple mutant, and Bonnie Bartel for providing the *ilr1-1 ill2-1 iar3-2* triple mutant and vectors expressing GST-recombinant version of ILR1, ILL2, and IAR3. We also thank Itay Mayrose and Keran Halabi for their assistance in constructing ILR1/ILLs phylogenetics. This work was supported by funding from the Israel Science Foundation (grant number 1057/21 to R.W.), the Chief Scientist of the Ministry of Agriculture and Rural Development, Israel (grant numbers 20-10-0067, 13-37-0005 and 20-01-0270 to R.W. and E.S.), the US-Israel Binational Agricultural Research and Development Fund (BARD, grant number IS-5195-19R to R.W, E.S, and Chris J. Staiger (Purdue University, IN)), the Yuri Milner 70@70 Fellowship (to O.R.), the Deutsche Forschungsgemeinschaft (SFB024 TPB12 to C.D and SFB924 TPA08 and HA3468/6-3 to U.Z.H) and the Tel Aviv University Center for AI and Data Science (to N.B.T).

## Author Contributions Statement

J.R., E.S. and R.W. conceived of the project and E.S. and R.W. supervised the project. O.R. designed and ran experiments and analyzed data. S.Y., O.S., A.E. and P.T. ran rooting experiments of cuttings from the different trees. I.V. synthesized and characterized compounds. A.E. and V.D. performed RNA-seq and qPCR experiments. F.S. ran mass-spectrometry analyses of auxins and conjugates and analyzed data under the supervision of M.C.W. A.K. designed and ran molecular simulation experiments with input from O.R. under the supervision of N.B.T. A.F.D. performed bioinformatics analysis of the RNA sequencing data. R.N. designed and performed surface plasmon resonance measurements. D.P.J. performed oocyte uptake assay under the supervision of U.Z.H. U.Z.H. performed SSM assays. V.P. performed mass spectrometry analyses on oocyte extracts and membranes under supervision of C.D. K.L.U. performed PIN purification under supervision of B.P.P. A.C. performed and analyzed binding assays of auxin receptors under the supervision of E.K. E.S., R.W. and O.R. wrote the manuscript with inputs from all co-authors.

## Methods

### Trees material and experimental procedure

*Eucalyptus grandis* trees were grown in 50-liter pots containing peat and TUFF (70:30) and 2 g/L Osmokot. For rooting assays, ten-year-old trees grown in a net-house were pruned at 1.5–2 meter above the ground. The cuttings used for rooting assays (10–15 cm long) included the top 2–4 leaves, and ∼60% of each leaf was removed. After excision, the base and/or the foliage were treated as indicated. The submergence treatment was 1 min dipping, the foliar treatment was spraying with 0.05% Triton X-100 as a surfactant. The cuttings were planted in rooting medium; peat, vermiculite and polystyrene flakes in a ratio of 1:2:3, placed in a heated rooting table (25 °C) under 90% humidity. Fungicides were applied on a weekly basis.

### Metabolites extraction from cuttings

Samples were washed in soap for 1 min and two consecutive 1 min washes in double distilled water before collecting the tissue into liquid nitrogen. Each biological sample included tissue from at least 20 cuttings, except of the samples that were taken for K-IBA uptake analysis that included 10 cuttings. Next, samples were grinded in liquid nitrogen, using a lab mill (IKA). From each biological sample, 3 samples of 180–240 mg (fresh weight) frozen tissue were extracted in 1 mL of cold 79% isopropanol, 20% methanol and 1% acetic acid. When endogenous auxins were measured, 20 ng of ^12^C-labeled internal standards were added to the extraction solution. Samples were vortexed for 1 h at 4 °C, then centrifuged at 12,000 x g for 15 min. The supernatants were transferred to fresh 2 mL Eppendorf tubes, and the pellet was extracted two more times using 0.5 mL of extraction solvent. Supernatants from the same sample were combined, and evaporated to dryness in a SpeedVac. The pellets were dissolved in pre-chilled 200 µL of 50% methanol, centrifuged, and the supernatants were filtered through 13 mm 0.22 µm PDFV syringe filters into fresh tubes. The extracts were kept at −20 °C until UPLC-MS/MS analysis.

### Quantification of metabolites extracted from cuttings

UPLC-MS/MS analyses of **1q**, **1r**, 4-CPA and endogenous auxins were conducted using an UPLC-Triple Quadrupole-MS (Waters Xevo TQ MS). Separation was performed on a Waters Acquity UPLC BEH C18 1.7 µm 2.1 x 100 mm column with a VanGuard precolumn (BEH C18 1.7 µm 2.1 x 5 mm). UPLC separation used a water-acetonitrile gradient. For **1q** and **1r**, the gradient was of 5% to 70% solvent B in 8 min, following by 70% to 95% in 1 min, 3 min at 95%, 95% to 5% in 1 min and finally 3 min at 5% solvent B. For 4-CPA and endogenous auxins the gradient was of 5% to 60% solvent B in 7 min, following by 60% to 95% in 1 min, 3 min at 95%, 95% to 5% in 1 min and finally 3 min at 5% solvent B (solvent A = water, solvent B = acetonitrile, both contain 0.1% formic acid as an additive). The flow rate was 0.3 mL/min, the injection volume was 10 µL, and the column temperature was kept at 35 °C. The analyses were performed using the ESI source in negative ion mode with the following settings: capillary voltage 3.1 KV, cone voltage 30 V, desolvation temperature 300 °C (**1q** and **1r**) or 400 °C (4-CPA), desolvation gas flow 565 L/h, source temperature 140 °C. Quantitation was performed using multiple-reaction monitoring (MRM) acquisition by monitoring the 387/327, 387/355 for **1q**, RT – 7.75, 373/327, 373/130 for **1r**, RT – 6.90, and 185/127, 185/141 for 4-CPA, RT - 5.77. Dwell time of 78 msec (**1q** and **1r**) and 161 msec (4-CPA) for each transition. A calibration curve was used to calculate concentrations. Acquisition of LC-MS data was performed under MassLynx V4.1 software (Waters). For simplicity purposes, when a certain metabolite was not detected in all replicates of a biological group, it was assigned as not detected. When at least one replicate could be measured, not detected replicates were transformed to 0, with the exception of Figure 25 in the Supplementary information where the not detected replicate is not shown.

### Enriching for *Eucalyptus* cambium, RNA extraction and Bioinformatic analysis

Bases of *E. grandis* cuttings were either harvested immediately (T_0_ sample), or submerged for 1 min in 6,000 ppm K-IBA (Sigma) and planted for 24 hours prior to harvest. Cambium cells were isolated as previously described^2,3^. Briefly, bark of each cutting was rapidly peeled off using a sharp scalpel, and the inner tissue from the bark as well as residual soft tissue from the stem was scraped gently and immediately frozen in liquid nitrogen. The latter was referred as cambium enrich fraction. The rest of the inner stem part was used as xylem enriched fraction. An average yield of 50 mg was harvested per biological sample. RNA was extracted using Norgen-Bioteck RNA Extraction Kit according to the basic manufacturer protocol, including an on-column DNAase treatment. Samples of cambium enriched fractions were sent for sequencing to Macrogen laboratories in South Korea. The raw reads were subjected to a filtering and cleaning procedure. The FASTX Toolkit (http://hannonlab.cshl.edu/fastx_toolkit/index.html, version 0.0.13.2) was used to trim read-end nucleotides with quality scores <30, using the FASTQ Quality Trimmer, and to remove reads with less than 70% base pairs with a quality score ≤ 30 using the FASTQ Quality Filter. Clean-reads were aligned to the *E. grandis* genome extracted from the Phytozome database (Eucalyptus_grandisv2; https://phytozome.jgi.doe.gov/pz/portal.html) using Tophat2 software (v2.1)^4^; gene abundance estimation was performed using Cufflinks (v2.2)^5^, combined with gene annotations from the Phytozome. Differential expression analysis was completed using the DESeq2 R package^6^.

### qRT-PCR

For quantitative real time PCR, isolated RNA (from cambium enriched or xylem enriched fractions) was treated with DNAse 1 (Thermo scientific, CA, USA). cDNA was synthesized from 1 μg of total RNA using qPCRBIO cDNA synthesis kit (PCR Biosystems Ltd, UK). Each qRT-PCR reaction was performed in a 10 µl reaction volume containing cDNA sample, 225 nM of each forward and reverse primer (Supplementary Table 1), and Fast SYBR Green qPCR Master Mix (Applied Biosystems, USA) in StepOne Real-Time PCR System (Applied Biosystems, USA). The thermal cycling conditions were: 95 °C for 20 sec, 40 cycles at 95 °C for 3 sec, 60 °C for 30 sec. After the final cycle, a melting curve analysis was performed at 95 °C for 15 sec and 60 °C for 60 sec, followed by 95 °C for 15 sec to verify reaction specificity. While *WOX4* [(*WUSCHEL related homeobox 4*) - (*Eucgr.F02320*) and *HB8* [(*Homeobox gene 8*) - (*Eucgr.C00605*)] were used as cambium markers, *Isocitrate dehydrogenase* [(*IDH*) - (*Eucgr.F02901*)], and *α-tubulin* [(*A-Tub*) - (*Eucgr.F00470*)] were used as the internal control for normalization of expression^7^. Relative expression of genes was calculated according to the delta-delta Ct method^8^.

### *Arabidopsis* materials and experimental procedures

*Arabidopsis* plants were grown under long-day fluorescent light conditions (16 h light per day, 21 °C, 100–150 μE m^−2^ s^−1^ light intensity). The mutant lines *aux1-7*^9^, and *gh3* octuple^10^, and the auxin reporter lines *DR5:Luciferase*^11^, and *DR5:Venus*^12^, are in Col-0 background. The triple mutant line *ilr1-1 ill2-1 iar3-2*^13^ and the over-expression complementation lines are in Ws-2 background. To test root elongation, 5-day-old seedlings (germination was determined by radicle emergence) were transferred to the mentioned treatments and incubation times. Next, plates were scanned and root length was determined manually in Fiji ImageJ^14^ platform. To assay adventitious roots formation, germinated seeds were transferred to dark conditions for 3 days to induce etiolation. Next, etiolated seedlings were transferred to a split-dish containing the indicated treatment only in the upper part, so only the shoots (defined as hypocotyl above the root-shoot junction cotyledons and) were directly exposed. Following 1.5 h, the seedlings were transferred to fresh media for 10 min, then transferred again to a fresh media and allow to grow for 10 days. To test luciferase activity during AR induction, 4 biological repeats (each of ∼20 plants per plate) per treatment were used. Each plate was imaged three times; 24, 48 and 72 h after the mentioned treatment. 3 h before each imaging, seedlings were transferred to media that were pre-sprayed with 0.5 mL of 1 mM D-Luciferin (GOLDBIO) with 0.01% Tween-20 as a surfactant. Images were taken by IVIS Lumina III (PerkinElmer) with a constant exposure time of 2 min and the total flux (p/s) values were used to determine enzymatic activity. In all assays, the media was half-strength MS (Duchefa) at pH 5.7, supplemented with 1% sucrose and 0.8% plant-agar (Duchefa).

### Confocal microscopy and image analysis

*Arabidopsis* root tips were imaged with a Zeiss LSM 780 laser spectral scanning confocal microscope, with a 10X air (Plan-Neofluar 10X/0.3 M27) objective. YFP was exited with a 514 nm argon laser. To determine Venus fluorescence, Z-stack images were acquired, and signal intensity was quantified using Fiji imageJ^14^ using the whole root tip as region of interest.

### Expression, purification and activity evaluation of GST-recombinant enzymes

To express GST-recombinant enzymes, vectors^15^ were transformed into the BL21 (DE3) strain of *E. coli* and positive colonies were selected on 100 µg/ml carbenicillin. 1 mL starter from fresh colony was used to inoculate 100 mL culture of LB media containing 100 µg/mL ampicillin. The cells were grown to an OD_600_ of 0.6 at 37 °C, then protein expression was induced with 100 µM IPTG. The cells were then grown for 12–16 hours at 16 °C, pelleted, kept over-night in −80 °C, resuspend with lysis buffer (TRIS-HCl 7.4 20 mM, DTT 2 mM, 0.05% Triton-100 and PMSF 1 mM) and kept over-night in −80 °C. Next, the lysate was incubated with 1 mg/mL Lysozyme for 30 min in room temperature, before further mild sonication lysis. Lysate was centrifuged and filtered through 22 µm filter, then passed over the Pierce Glutathione Spin Column according to the manufacturer protocol. Correct-size validation was performed by SDS-PAGE and the protein concentration was determined by Pierce BCA Protein Assay according to the manufacturer protocol. Enzymatic activity was examined in a reaction solution (50 mM TRIS-HCl pH 8, 1mM DTT, 1mM MnCl_2_, 0.1 mM substrate and approximately 20 ng/µL protein) for 24 hours in 25 °C. The reactions were terminated by the addition of 1% acetic acid in methanol to a final ratio of 1:3, and the products were analyzed by HPLC-MS, using Waters HPLC with XBridge C18 column (100 × 3 mm, 5 µm) coupled with LC/MS Acquity QDa. HPLC separation used water-acetonitrile gradient of 10% to 100% solvent B in 16 min then 5 min at 100% solvent B and finally 4 min at 10% solvent B, at flow rate of 1 mL/min (solvent A = water, solvent B = acetonitrile, both contain 0.1% TFA as an additive). The compounds were identified according to their absorbance at 215 nm.

### Phylogenetic analysis

Reciprocal Protein Basic Local Alignment Search Tool (BLAST) was used to identify homologs of the *Arabidopsis* ILR1/ILLs, and sequences were retrieved from the Phytozome and PhycoCosm databases. 355 proteins with at least 30% identity were found, of which the length of 277 sequences (78%) is 400-500 amino acids (Supplementary Fig. 26), and sequences of 350-550 amino acids were selected (301 sequences) for further analysis. Multiple sequence analysis was generated by MAFFT using the E-INS-i strategy for proteins with more than one domain; based on the ILL2 structure^16^ that has two domains (Fig. 5a) or using automate settings (Supplementary Fig. 23). Quality of the MSAs was evaluated by Guidance2^17^, and both MSAs had score higher than 0.93. Next, the trees were built using IQ-TREE^18^ with JTT+I+G4 substitution model (based on the program automatic fitting), 1000 ultrafast-bootstrap and SH-aLRT branch test replicates. The R packages ggtree^19^, and treeio^20^ were used for visualizations.

### Cloning and plant transformation

Based on sequences targeting ILL1,3,5,6 (calculated using the Chop-Chop tool^21^; Supplementary Table 2) two guide DNAs were generated for each gene by PCR using pICH86966 as template (Supplementary Table 2), and cloned downstream the AtU6 promoter (Level 0 vector; pICSL01009)^22^ to form Level 1 vector, using the Golden Gate cloning method^23^. Next, four Level 1 vectors were similarly assembled into binary vector that encodes for intronized CAS9 and Basta resistance cassette (pAGM65879)^24^, to generate one vector that targets ILL1 and ILL6, and another that targets ILL3 and ILL5. To overexpress enzymes in *Arabidopsis*, the coding sequences of *EgK02589*, *EgK02598*, *Potri.006G207400*, *Prupe.7G100000* and *Potri.016G074100* were retrieved from the Phytozome database. Codon optimization (to *Arabidopsis thaliana* codon usage), synthesis and cloning into pENTR vector were all conducted by Twist Bioscience. Overexpression binary vectors were generated by LR reaction into pH2GW7 destination vector that encodes for hygromycin resistance cassette^25^. *Arabidopsis* transgenic lines were generated by floral-dipping^26^, using the GV3101 strain of *Agrobacterium tumefaciens*, and resistant lines were selected on 10 µg/mL phosphinothricin or 20 µg/mL hygromycin.

### Yeast two-hybrid assays

For the qualitative assays, TIR1 bait vector pGILDA was co-transformed^27^ with IAA7 prey vector pB42AD (or empty pB42AD as negative control) into *Saccharomyces cerevisiae* strain EGY48, as described^28^. Yeast two-hybrid assays were performed as described previously^27^. For the quantitative assays, yeast strains were constructed as described previously^29^. Synthetic complete growth medium was used to grow the cells overnight from glycerol stock. All yeast cultures in all experiments of this study were grown in a 30 °C shaker incubator at 250 RPM. Steady state data were collected during log phase via the following preparation: 16 h of overnight growth in the synthetic complete medium in a shaker incubator followed by dilution to 30 events/µL into fresh, room-temperature medium. After 10 h of growth at 30 °C, a new dilution to 30 events/µL in 3 mL of medium was performed, the inducers were added to the indicated concentrations and 100 µL samples were collected at complete response time. Ten thousand events were collected for each condition. Experimental replicates are intended as biological replicates of the same overnight sample (replicates conducted on the same day) or the same glycerol stock (replicates conducted on different days).

### TIR1 Protein expression, purification and binding analysis by surface plasmon resonance

Protein for auxin binding assays and the analysis by SPR was done as described previously^30^. Briefly, *Arabidopsis TIR1*, was codon-optimized for expression in insect and cloned into pOET5 transfer vector (Oxford Expression Technologies, Oxford, U.K.) with Arabidopsis *ASK1.* Recombinant baculoviruses were used to infect *Spodoptera frugiperda9 (Sf9)* cells in tissue culture. Cells were harvested by centrifugation 2 days post infection and stored at −80 °C. Cell lysates were loaded onto a nickel immobilized metal affinity chromatography column (cOmplete His-Tag Purification Resin, Roche) and eluted with 250 mM imidazole. Auxin binding assays on a Biacore T200 (Cytiva Life Sciences) were done as described previously^31^. Biotinylated AtAux/IAA7 degron peptide was immobilized on streptavidin-coated SPR chips, and binding was measured in the presence of IAA, or auxin analogue, by recruitment of the TIR1/AFB protein from solution as the co-receptor complex formed on the chip. Auxins were maintained as stock solutions in DMSO and used at with DMSO at 1% final concentration.

### Oocyte uptake assay

Experiments were performed as described before^32^ with the following changes: *Xenopus laevis* oocytes were injected with 150 ng AUX1 cRNA. The stock solutions of IAA of 4-CPA, in methanol were diluted in 2 ml Barth’s solution (88 mM NaCl, 1 mM KCl, 0.8 mM MgSO_4_, 0.4 mM CaCl_2_, 0.3 mM Ca(NO_3_)_2_, 2.4 mM NaHCO_3_, 10 mM HEPES) to reach a final concentration of 20 µM. As a negative control, 2 µL of methanol were diluted in 2 mL Barth’s solution. The oocytes were incubated at room temperature for 30 min and 1 h in the respective solution (n = 35 oocytes per sample and time point). Oocytes were washed twice in Barth’s solution, transferred to a reaction tube and homogenized in homogenization buffer without PhosSTOP (400 µl per 35 oocytes). The soluble fraction (cytosol) was stored at −80 °C until analysis by LC-MS/MS. The assay was repeated three times with oocytes collected from different females.

### Quantification of metabolites extracted from *Xenopus* oocytes

Samples were incubated with 20 µL of IS solution (155 µmol/L forchlorfenuron, 25.5 µmol/L 4-chlorphenylacetic acid, 5.16 mmol/L indole-3-acetic-2,2-d2 acid, Sigma-Aldrich, Steinheim, Germany) and diluted with an acetonitrile/methanol + 1% formic acid solution (4:1; v/v) in extractions-tubes. Extraction was performed using a bead beater homogenizer (8,000 RPM for 3 × 30 sec and 25 sec breaks, Precellys evolution Homogenizer, Bertin Technologies, Montigny Le Bretonneux, France) at 0 °C and after equilibration (1 h) and centrifugation (10 min, 6,000 x g) the supernatant was membrane filtered (Minisart RC 15, 0.45 µm, Sartorius AG, Göttingen, Germany) and used for analysis. Calibration curve: A stock solution for the quantitation of 4-CPA (339.9 µmol/L) and IAA (3.22 µmol/L) were prepared in methanol (1 mL) and the exact concentrations of each reference compound was verified by means of qHNMR in methanol-d4^33^. This calibration stock solution was then sequentially diluted 1+1. To each dilution IS (20 µL), were added before analysis. For the recovery experiments the analytes were spiked into analytes free cytosols and membranes using the concentration ranges of the Calibration Curve each as triplicates. After addition of IS (20 µL) the samples were prepared following the instructions above. For the determination of the limits of detection (LoD) and the limits of quantitation (LoQ) the standard solutions were further diluted. For the LoD, the signal-to-noise was set to a ratio of 3, and for the LoQ to a ratio of 9. The intraday precision was determined by calculating the relative standard deviation (RSD) for the analysis of the spiked recovery samples and interday precision was determined by the analysis of the spiked recovery samples after 4 days. LC-MS/MS analysis: The samples were chromatographically separated by means of an ExionLC (Sciex, Darmstadt, Germany), consisting of two LC pump systems ExionLC AD Pump, an ExionLC degasser, an ExionLC AD autosampler, an ExionLC AC column oven, and an ExionLC controller, and connected with a QTRAP 6500+ mass spectrometer (Sciex, Darmstadt, Germany) controlled by the Analyst 1.6.3 software (Sciex, Darmstadt, Germany). Data interpretation was performed using MultiQuant software (version 3.0.2, Sciex, Darmstadt, Germany; Peak model: MQ4) and Analyst 1.6.3 (Sciex, Darmstadt, Germany). LC-MS/MS analysis of 4-CPA and IAA: the compounds were separated on a Kinetex C18 column (100 x 2.1 mm, 1.7 µm, 100 A, Phenomenex, Aschaffenburg, Germany) with a flow rate of 0.4 mL/min. The following gradient consisting of formic acid in water (0.1%, solvent A) and 0.1% formic acid in methanol (0.1%, solvent B) was used: 0 min, 15% B; 1 min, 15% B; 5 min, 100% B; 6 min, 100% B; 7 min, 15% B; 8 min, 15% B was used for the separation of the compounds. Mass spectrometer settings: MRM-(low mass), ion spray voltage (4,500 V), curtain gas (35 psi), temperature (450 °C), gas 1 (55 psi), gas 2 (65 psi), collision activated dissociation (−2 V), and entrance potential (10 V). MRM+ (low mass), ion spray voltage (4,500 V), curtain gas (35 psi), temperature (450 °C), gas 1 (55 psi), gas 2 (65 psi), collision activated dissociation (2 V), and entrance potential (10 V). Quantitative ^1^H Nuclear Magnetic Resonance Spectroscopy (qNMR): Quantitative proton NMR spectroscopy (qHNMR) was recorded on a 400 MHz Avance III spectrometer (Bruker, Rheinstetten, Germany) equipped with a Broadband Observe BBFO plus Probe (Bruker, Rheinstetten, Germany). Methanol-*d_4_* (600 µL) was used as solvent and chemical shifts are reported in ppm relative to the methanol d4 solvent signal. Data processing was performed using Topspin NMR software (vers. 3.2, Bruker, Rheinstetten, Germany). The quantitation by qHNMR was performed as reported earlier by calibration of the spectrometer using the ERETIC 2 tool based on the PULCON methodology.

### Solid-supported membrane electrophysiology assay

Solid supported membrane (SSM) electrophysiology was carried out as described^34^. Proteoliposomes with an lipid to protein ratio of 1:5 were kept in non-activating solution; Ringer solution without Ca2+ (115 mM NaCl, 2.5 mM KCl, 1 mM NaHCO3, 10 mM HEPES pH 7.4, 1 mM MgCl2). At the beginning of the experiment non-activating buffer was exchanged for fresh identical non-activating buffer and after 1 s activating buffer (same buffer containing 20 µM substrate) was added. After a further 1 sec, buffer was again exchanged to non-activating buffer. Current response was recorded throughout the entire 3 sec. Peak current responses were extracted from the current traces using the SURFE2R control v 1.6.0.1 software with the default settings.

### Docking calculations

Docking calculations were carried out using the Glide software^35–37^ (Glide, Schrödinger, LLC, New York, NY, 2021), as part of Schrödinger Release 2021-4. The docking used the available solved structure of ILL2^16^ and the AlphaFold^38^ models for all other enzymes. The structures of the above proteins are overall similar (rmsd of 2.3 Å at the most), particularly in the active site region. Thus, to place the two Mn^2+^ ions in each of the predicted structures, we used the ILL2 crystal structure as a template, and made only small manual adjustments to compensate for differences in the exact locations of the coordinating residues in other sites. Each of the structures was prepared for docking and energy-minimized using the Schrodinger’s Protein Preparation Wizard^39^ (Schrödinger LLC, New York, NY), with protonation states predicted by PropKa 3.1^40,41^. The structure of **1r** was prepared for docking using LigPrep (LigPrep, Schrödinger, LLC, New York, NY, 2021), with its ionization state determined at pH 7.0±2.0 using Epik^42^. In the ligand’s structure, Trp was kept in the (*2S*) configuration. The receptor grid was generated around the centroids of the catalytic glutamate residue (for example E172 in ILL2) and Mn^2+^ cations, with a 10 Å enclosing box. Docking calculations were carried out using the extra precision (XP) scoring function^35^, with flexible ligand and expanded sampling.

### General synthetic and analytical methods

All chemicals were purchased from Sigma-Aldrich or Combi-Blocks and used as received unless otherwise stated. Anhydrous solvents and reagents (DCM, THF) were obtained as SureSeal bottles from Sigma-Aldrich. Thin-layer chromatography and flash chromatography were performed using EMD pre-coated silica gel 60 F-254 plates and silica gel 60 (230–400 mesh), respectively. HPLC-MS analysis was performed on Waters HPLC coupled with Acquity QDa (low resolution ESI) with XBridge C18 column (100 × 3 mm, 5 *µ*m) using a water-acetonitrile gradient of 0% to 100% solvent B in 17 min then 3 min at 100% solvent B at flow rate of 1 mL/min (solvent A = water, solvent B = acetonitrile, both contain 0.1% TFA as an additive). High resolution ESI mass spectrometry was performed on a Waters SYNAPT system. ^1^H- and ^13^C-NMR spectra were collected in CDCl_3_, or DMSO-*d_6_* (Cambridge Isotope Laboratories, Cambridge, MA) at 25 °C using a Bruker Advance III spectrometer at 400 MHz and 101 MHz respectively at the Department of Chemistry NMR Facility at Tel-Aviv University. All chemical shifts are reported in the standard δ notation of parts per million using the either TMS or residual solvent peak as an internal reference. Abbreviations: THF: tetrahydrofuran, DMF: dimethylformamide, DCM: dichloromethane, DIPEA: diisopropylethylamine, DMSO: dimethylsulphoxide, Et_3_N: trimethylamine, EtOAc: ethyl acetate, Hex: n-hexanes, TFA: trifluoroacetic acid, RT: room temperature, TLC: thin later chromatography, CDI: carbomyldiimidazole, NAA: 1-naphthaleneacetic acid, 4-CPA: 4-chlorophenoxyacetic acid, MCPA: 2-methyl-4-chlorophenoxyacetic acid, 2-DP: 2-(2,4-dichlorophenoxy) propionic acid.

### LogD calculation

LogD calculations were conducted using the Chemaxon LogD Predictor plugin. https://disco.chemaxon.com/calculators/demo/plugins/logd/.

### p*K*_a_ prediction

p*K*_a_ predictions were conducted using MolGpKa^43^ server https://xundrug.cn/molgpka.

### Synthetic methods

All compounds were synthesized according to one of the following methods, as appears in Supplementary Table 3:

#### Method A

To a solution of 4-CPA, MCPA or 2-DP (300–500 mg) in DCM (30 mL) and few drops of THF, CDI (1.2 eq.) was added. After stirring the solution for 2 h at room temperature, the amine was added (1.05 eq.). For aromatic amines the reaction was stirred overnight. For aliphatic amines the reaction was monitored by TLC. Upon completion, the reaction mixture was washed with water, the organic phase was separated, and the aqueous residue was extracted with DCM (2 × 20 mL). The combined organic phase was washed with 1 M HCl (20 mL), dried over MgSO_4_ and concentrated under vacuum. Purity of product was analyzed by HPLC-MS. If needed, the crude residue was purified by silica gel chromatography (EtOAc:Hex). 45–95% yield.

#### Method B

SOCl_2_ (5 eq.) was slowly added to a solution of NAA in DCM (30 mL) at 0 °C. After stirring at room temperature for 2 h, the reaction mixture was concentrated in vacuum and the crude acid chloride was diluted with 30 mL of DCM. A solution of the amine (1 eq.) and NEt_3_ (1 eq.) in DCM (20 mL) was added dropwise to acid chloride solution at 0 °C. The resulting mixture was allowed warm to room temperature and stirred overnight. The reaction mixture was quenched with saturated NaHCO_3_ and extracted with DCM (2 × 20 mL). The combined organic phase was dried over MgSO_4_ and concentrated under vacuum. The crude amide product was purified by silica gel chromatography (EtOAc:Hex). Purity of product was analyzed by HPLC-MS. 30–90% yield.

#### Method C

SOCl_2_ (3 eq.) was slowly added to a solution of NAA, 4-CPA, MCPA or 2-DP (300–400 mg, 1eq) in methanol (30 mL) at room temperature. The mixture was heated to 60 °C and stirred for 30 min. The reaction mixture was concentrated in vacuum, diluted with ethyl acetate, washed with saturated NaHCO_3_, brine, dried over MgSO_4_ and concentrated under vacuum. Purity of product was analyzed by HPLC-MS. If needed, the crude residue was purified by silica gel chromatography (EtOAc:Hex). 83–97% yields.

### Synthesis of 1r and 1t

Into a 10 mL process vial equipped with a stirring bar, **1q** or **1s** (200 mg, 0.52 mmol, 1 eq.) in 3 mL of methanol added, followed by the addition of aqueous solution of sodium hydroxide (3 eq. in 1 mL water). The vial was fitted with a snap-on cap, inserted to CEM Discover SP microwave and stirred for 10 sec under the following conditions: method type: Dynamic, pressure limit: 250 PSI, vessel Type: 10 mL, temperature: 90 °C, power: 100 W, hold time (h:m:s): 00:10:00, permixing: no, stirring: high, cooling: on. The solution was transferred to 20 mL vail and evaporated to dryness. The crude was dissolved in H_2_O (5 mL) and the pH is adjusted to 3 with 2N HCl. When precipitation of product is complete, the solid was filtered and washed with water. The solid was lyophilized overnight. Finally, a stochiometric amount of NaOH in 5 mL water added and the solid was lyophilized, providing the final product as sodium salt. White powder. Yields: **1r** 185 mg (0.47 mmol, 90%), **1t** 188 mg (0.48 mmol, 92%).

### Data Availability

All data supporting the findings of this study are available within the paper and its Supplementary Information. RNA sequencing data associated with this work is available at BioProject accession PRJNA1029024 (manuscript ref. ^91^).

## References

1. Verstraeten, I., Schotte, S. & Geelen, D. hypocotyl adventitious root organogenesis differs from lateral root development. Front. Plant Sci. 5, (2014).

2. Hartmann, H. T., Kester, D. E., Davis, F. T., Geneve, R. L., Wilson, S. B., Hartmann & Kester’s Plant Propagation: Principles and Practices (Pearson, 2017)

3. Poethig, R. S. Phase change and the regulation of shoot morphogenesis in plants. Science (1979) 250, 923–930 (1990).

4. Pijut, P. M., Woeste, K. E. & Michler, C. H. Promotion of adventitious root formation of difficult-to-root hardwood tree species. Hortic. Rev. 38, 213 (2011).

5. Hackett, W. P. Juvenility, maturation, and rejuvenation in woody plants. Hortic. Rev. 7, 109–154 (2011).

6. Wilcox, J. R. & Farmer, R. E. Heritability and C effects in early root growth of eastern cottonwood cuttings. Heredity 23, 239–245 (1968).

7. Grattapaglia, D., Bertolucci, F. L. & Sederoff, R. R. Genetic mapping of QTLs controlling vegetative propagation in *Eucalyptus grandis* and *E. urophylla* using a pseudo-testcross strategy and RAPD markers. Theoretical and Applied Genetics 90, 933–947 (1995).

8. Visser, E. J. W., Cohen, J. D., Barendse, G. W. M., Blom, C. W. P. M. & Voesenek, L. A. C. J. An ethylene-mediated increase in sensitivity to auxin induces adventitious root formation in flooded Rumex palustris Sm. Plant Physiol. 112, 1687–1692 (1996).

9. Sorin, C. et al. Auxin and light control of adventitious rooting in Arabidopsis require ARGONAUTE1. Plant Cell 17, 1343–1359 (2005).

10. Bellini, C., Pacurar, D. I. & Perrone, I. Adventitious roots and lateral roots: similarities and differences. Annu. Rev. Plant Biol. 65, 639–666 (2014).

11. Pacurar, D. I., Perrone, I. & Bellini, C. Auxin is a central player in the hormone cross-talks that control adventitious rooting. Physiol. Plant. 151, 83–96 (2014).

12. Caboni, E. et al. Biochemical aspects of almond microcuttings related to in vitro rooting ability. Biol. Plant. 39, 91–97 (1997).

13. Lakehal, A. et al. A molecular framework for the control of adventitious rooting by TIR1/AFB2-Aux/IAA-dependent auxin signaling in *Arabidopsis*. Mol. Plant 12, 1499–1514 (2019).

14. Bellamine, J., Penel, C., Greppin, H. & Gaspar, T. Confirmation of the role of auxin and calcium in the late phases of adventitious root formation. Plant Growth Regul. 26, 191–194 (1998).

15. Blažková, A. et al. Auxin metabolism and rooting in young and mature clones of *Sequoia sempervirens*. Physiol. Plant. 99, 73–80 (1997).

16. Rasmussen, A., Hosseini, S. A., Hajirezaei, M.-R., Druege, U. & Geelen, D. Adventitious rooting declines with the vegetative to reproductive switch and involves a changed auxin homeostasis. J. Exp. Bot. 66, 1437–1452 (2015).

17. Abu-Abied, M. et al. Microarray analysis revealed upregulation of nitrate reductase in juvenile cuttings of *Eucalyptus grandis*, which correlated with increased nitric oxide production and adventitious root formation. Plant J. 71, 787–799 (2012).

18. Abarca, D. et al. The GRAS gene family in pine: transcript expression patterns associated with the maturation-related decline of competence to form adventitious roots. BMC Plant Biol. 14, 354 (2014).

19. Fahn, A. Plant Anatomy. (Pergamon, 1990)

20. Uggla, C., Moritz, T., Sandberg, G. & Sundberg, B. Auxin as a positional signal in pattern formation in plants. Proc. Natl. Acad. Sci. U.S.A 93, 9282–9286 (1996).

21. Tuominen, H., Puech, L., Fink, S. & Sundberg, B. A radial concentration gradient of indole-3-acetic acid is related to secondary xylem development in hybrid aspen. Plant Physiol. 115, 577–585 (1997).

22. Ballester, A., San-José, M. C., Vidal, N., Fernández-Lorenzo, J. L. & Vieitez, A. M. Anatomical and biochemical events during in vitro rooting of microcuttings from juvenile and mature phases of chestnut. Ann. Bot. 83, 619–629 (1999).

23. Vidal, N., Arellano, G., San-José, M. C., Vieitez, A. M. & Ballester, A. Developmental stages during the rooting of in-vitro-cultured Quercus robur shoots from material of juvenile and mature origin. Tree Physiol 23, 1247–1254 (2003).

24. Legué, V., Rigal, A. & Bhalerao, R. P. Adventitious root formation in tree species: involvement of transcription factors. Physiol. Plant. 151, 192–198 (2014).

25. Ranjan, A. et al. Molecular basis of differential adventitious rooting competence in poplar genotypes. J. Exp. Bot. 73, 4046–4064 (2022).

26. Díaz-Sala, C. Direct reprogramming of adult somatic cells toward adventitious root formation in forest tree species: the effect of the juvenile–adult transition. Front. Plant Sci. 5, (2014).

27. Solé, A. et al. Characterization and expression of a Pinus radiata putative ortholog to the Arabidopsis SHORT-ROOT gene. Tree Physiol. 28, 1629–1639 (2008).

28. Abu-Abied, M. et al. Gene expression profiling in juvenile and mature cuttings of *Eucalyptus grandis* reveals the importance of microtubule remodeling during adventitious root formation. BMC Genom. 15, 826 (2014).

29. de Almeida, M. R. et al. Reference gene selection for quantitative reverse transcription-polymerase chain reaction normalization during in vitro adventitious rooting in *Eucalyptus globulus* Labill. BMC Mol. Biol. 11, 73 (2010).

30. de Almeida, M. R. et al. Comparative transcriptional analysis provides new insights into the molecular basis of adventitious rooting recalcitrance in *Eucalyptus*. Plant Sci. 239, 155–165 (2015).

31. Ruedell, C. M., de Almeida, M. R. & Fett-Neto, A. G. Concerted transcription of auxin and carbohydrate homeostasis-related genes underlies improved adventitious rooting of microcuttings derived from far-red treated *Eucalyptus globulus* Labill mother plants. Plant Physiol. Biochem. 97, 11–19 (2015).

32. Cooper, W. C. Hormones in relation to root formation on stem cuttings. Plant Physiol. 10, 789 (1935).

33. Oinam, G., Yeung, E., Kurepin, L., Haslam, T. & Lopez-Villalobos, A. Adventitious root formation in ornamental plants: I. General overview and recent successes. Propag. Ornam. Plants 11, 78–90 (2011).

34. Epstein, E. & Ludwig-Müller, J. Indole-3-butyric acid in plants: occurrence, synthesis, metabolism and transport. Physiol. Plant. 88, 382–389 (1993).

35. Strader, L. C. & Bartel, B. Transport and metabolism of the endogenous auxin precursor indole-3-butyric acid. Mol. Plant 4, 477–486 (2011).

36. Wiesman, Z., Riov, J. & Epstein, E. Comparison of movement and metabolism of indole-3-acetic acid and indole-3-butyric acid in mung bean cuttings. Physiol. Plant. 74, 556–560 (1988).

37. Felker, P. & Clark, P. R. Rooting of mesquite (*Prosopis*) cuttings. J. Range Manag. 34, 466–468 (1981).

38. Van der Krieken, W. M. et al. Increased induction of adventitious rooting by slow release auxins and elicitors. In Biology of Root Formation and Development. 95–104 (Springer, 1997).

39. Mihaljevic, S. & Salopek-Sondi, B. Alanine conjugate of indole-3-butyric acid improves rooting of highbush blueberries. Plant Soil Environ. 58, 236–241 (2012).

40. Haissig, B. E. Influence of aryl esters of indole-3-acetic and indole-3-butyric acids on adventitious root primordium initiation and development. Physiol. Plant. 47, 29–33 (1979).

41. Pizarro, A. & Díaz-Sala, C. Cellular dynamics during maturation-related decline of adventitious root formation in forest tree species. Physiol. Plant. 165, 73–80 (2019).

42. Vilasboa, J., da Costa, C. T. & Fett-Neto, A. G. Rooting of eucalypt cuttings as a problem-solving oriented model in plant biology. Prog. Biophys. Mol. Biol. 146, 85–97 (2019).

43. Quareshy, M., Prusinska, J., Li, J. & Napier, R. A cheminformatics review of auxins as herbicides. J. Exp. Bot. 69, 265–275 (2018).

44. Eyer, L. et al. 2,4-D and IAA amino acid conjugates show distinct metabolism in *Arabidopsis*. PLoS One 11, e0159269 (2016).

45. Yang, Y., Hammes, U. Z., Taylor, C. G., Schachtman, D. P. & Nielsen, E. High-affinity auxin transport by the AUX1 influx carrier protein. Curr. Biol. 16, 1123–1127 (2006).

46. Hoyerova, K. et al. Auxin molecular field maps define AUX1 selectivity: many auxin herbicides are not substrates. New Phytol. 217, 1625–1639 (2018).

47. Calderón Villalobos, L. I. A., et al. A combinatorial TIR1/AFB–Aux/IAA co-receptor system for differential sensing of auxin. Nat. Chem. Biol. 8, 477–485 (2012).

48. Lee, S. et al. Defining binding efficiency and specificity of auxins for SCFTIR1/AFB-Aux/IAA co-receptor complex formation. ACS Chem. Biol. 9, 673–682 (2014).

49. Vain, T. et al. Selective auxin agonists induce specific AUX/IAA protein degradation to modulate plant development. Proc. Natl. Acad. Sci. U.S.A 116, 6463–6472 (2019).

50. Pufky, J., Qiu, Y., Rao, M. v, Hurban, P. & Jones, A. M. The auxin-induced transcriptome for etiolated Arabidopsis seedlings using a structure/function approach. Funct. Integr. Genom. 3, 135–143 (2003).

51. Delargy, J. A. & Wright, C. E. Root formation in cuttings of apple in relation to auxin application and to etiolation. New Phytol. 82, 341–347 (1979).

52. Verstraeten, I., Beeckman, T. & Geelen, D. Adventitious root induction in *Arabidopsis thaliana* as a model for in vitro root organogenesis. in Plant Organogenesis: Methods and Protocols 159–175 (Springer, 2013)

53. Grossmann, K. Auxin herbicides: current status of mechanism and mode of action. Pest Manag. Sci. 66, 113–120 (2010).

54. Ludwig-Müller, J., Vertocnik, A. & Town, C. D. Analysis of indole-3-butyric acid-induced adventitious root formation on Arabidopsis stem segments. J. Exp. Bot. 56, 2095–2105 (2005).

55. Blythe, E. K., Sibley, J. L., Tilt, K. M. & Ruter, J. M. Methods of auxin application in cutting propagation: a review of 70 years of scientific discovery and commercial practice. J. Environ. Hortic. 25, 166–185 (2007).

56. Riov, J. et al. Improved method for vegetative propagation of mature *Pinus halepensis* and its hybrids by cuttings. Isr. J. Plant Sci. 67, 5–15 (2020).

57. Eliasson, L. & Areblad, K. Auxin effects on rooting in pea cuttings. Physiol. Plant. 61, 293–297 (1984).

58. Wain, R. L., Wightman, F. & Russell, E. J. The growth-regulating activity of certain *ω*-substituted alkyl carboxylic acids in relation to their *β*-oxidation within the plant. Proc. R. Soc. Lond. B: Biol. Sci. 142, 525–536 (1954).

59. Behrens Richard & Howard Morton L. Some factors influencing activity of 12 phenoxy acids on mesquite root inhibition. Plant Physiol. 38, 165–170 (1963).

60. Zimmerman, P. W. Several chemical growth substances which cause initiation of roots and other responses in plants. Contrib. Boyce Thompson Inst. 7, 209–229 (1935).

61. Katekar, G. F. Auxins: on the nature of the receptor site and molecular requirements for auxin activity. Phytochemistry 18, 223–233 (1979).

62. Kepinski, S. & Leyser, O. The *Arabidopsis* F-box protein TIR1 is an auxin receptor. Nature 435, 446–451 (2005).

63. Dharmasiri, N., Dharmasiri, S. & Estelle, M. The F-box protein TIR1 is an auxin receptor. Nature 435, 441–445 (2005).

64. Xuan, W., Opdenacker, D., Vanneste, S. & Beeckman, T. Long-term in vivo imaging of luciferase-based reporter gene expression in Arabidopsis roots. in Root Development 177–190 (Springer, 2018).

65. Ung, K. L. et al. Structures and mechanism of the plant PIN-FORMED auxin transporter. Nature 609, 605–610 (2022).

66. Bainbridge, K. et al. Auxin influx carriers stabilize phyllotactic patterning. Genes. Dev. 22, 810–823 (2008).

67. Casanova-Sáez, R. et al. Inactivation of the entire Arabidopsis group II GH3s confers tolerance to salinity and water deficit. New Phytol. 235, 263–275 (2022).

68. Hayashi, K. et al. The main oxidative inactivation pathway of the plant hormone auxin. Nat. Commun. 12, 1–11 (2021).

69. Bartel, B. & Fink, G. R. ILR1, an amidohydrolase that releases active indole-3-acetic acid from conjugates. Science (1979) 268, 1745–1748 (1995).

70. Davies, R. T., Goetz, D. H., Lasswell, J., Anderson, M. N. & Bartel, B. *IAR3* encodes an auxin conjugate hydrolase from Arabidopsis. Plant Cell 11, 365–376 (1999).

71. LeClere, S., Tellez, R., Rampey, R. A., Matsuda, S. P. T. & Bartel, B. Characterization of a family of IAA-amino acid conjugate hydrolases from *Arabidopsis*. J. Biol. Chem. 277, 20446–20452 (2002).

72. Campanella, J. J., Olajide, A. F., Magnus, V. & Ludwig-Muller, J. A novel auxin conjugate hydrolase from wheat with substrate specificity for longer side-chain auxin amide conjugates. Plant Physiol. 135, 2230–2240 (2004).

73. Rampey, R. A. et al. A family of auxin-conjugate hydrolases that contributes to free indole-3-acetic acid levels during Arabidopsis germination. Plant Physiol. 135, 978–988 (2004).

74. Bitto, E. et al. X-ray structure of ILL2, an auxin-conjugate amidohydrolase from *Arabidopsis thaliana*. Proteins. 74, 61–71 (2009).

75. Jumper, J. et al. Highly accurate protein structure prediction with AlphaFold. Nature 596, 583–589 (2021).

76. Valentini, R., Mugnozza, G. S., Giordano, E. & Kuzminsky, E. Influence of cold hardening on water relations of three *Eucalyptus* species. Tree Physiol. 6, 1–10 (1990).

77. Eliyahu, A. et al. Vegetative propagation of elite Eucalyptus clones as food source for honeybees (*Apis mellifera*); adventitious roots versus callus formation. Isr. J. Plant Sci. 67, 83–97 (2020).

78. Robinson, T. L. et al. Performance of Cornell-Geneva rootstocks across North America in multi-location NC-140 rootstock trials. in Proceedings of the 1st International Symposium on Rootstocks for Deciduous Fruit Tree Species. 241–245 (2004, ISHS).

79. Marini, R. P. et al. Performance of ‘golden delicious’ apple on 23 rootstocks at 12 locations: a five-year summary of the 2003 nc-140 dwarf rootstock trial. J. Am. Pomol. Soc. 63, 115 (2009).

80. Marini, R. P. et al. Performance of ‘Golden Delicious’ apple on 23 rootstocks at eight locations: a ten-year summary of the 2003 NC-140 dwarf rootstock trial. J. Am. Pomol. Soc. 68, 54–68 (2014).

81. Tzeela, P. et al. Comparing adventitious root-formation and graft-unification abilities in clones of *Argania spinosa*. Front. Plant Sci. 13, (2022).

82. Ruberv, P. H. & Sheldrake, A. R. Effect of pH and surface charge on cell uptake of auxin. Nat. New Biol. 244, 285–288 (1973).

83. Sanchez Carranza, A. P., et al. Hydrolases of the ILR1-like family of Arabidopsis thaliana modulate auxin response by regulating auxin homeostasis in the endoplasmic reticulum. Sci. Rep. 6, 24212 (2016).

84. Staswick, P. E. et al. Characterization of an Arabidopsis enzyme family that conjugates amino acids to indole-3-acetic acid. Plant Cell 17, 616–627 (2005).

85. Hagen, G. & Guilfoyle, T. Auxin-responsive gene expression: genes, promoters and regulatory factors. Plant Mol. Biol. 49, 373–385 (2002).

86. Wabnik, K., Govaerts, W., Friml, J. & Kleine-Vehn, J. Feedback models for polarized auxin transport: an emerging trend. Mol BioSyst. 7, 2352–2359 (2011).

87. Hudson, J.P. Propagation of Plants by Root Cuttings: II. Seasonal Fluctuation of Capacity to Regenerate from Roots. J. Hortic. Sci. 30, 242–251 (1955).

88. Ohkouchi, T. & Tsuji, K. Basic Technology and Recent Trends in Agricultural Formulation and Application Technology. J. Pest Sci. 47, 155–171 (2022)

89. Skůpa, P., Opatrný, Z. & Petrášek, J. Auxin Biology: Applications and the Mechanisms Behind. In Applied Plant Cell Biology 69–102 (Springer, 2014)

90. Kessel, A. & Ben-Tal, N. Free energy determinants of peptide association with lipid bilayers. Curr. Top. Membr. 52, 205–253 (2002).

91. Roth, O. et al. Slow release of a synthetic auxin induces formation of adventitious roots in recalcitrant woody plants. Cambium-enriched samples of *Eucalyptus grandis* mature cuttings. BioProject. https://www.ncbi.nlm.nih.gov/bioproject/?term=PRJNA1029024 (2023).

## Methods-only references

1. Kessel, A. & Ben-Tal, N. Free energy determinants of peptide association with lipid bilayers. Curr. Top. Membr. 52, 205–253 (2002).

2. Ridoutt, B. G., Pharis, R. P. & Sands, R. Identification and quantification of cambial region hormones of *Eucalyptus globulus*. Plant Cell Physiol. 36, 1143–1147 (1995).

3. Foucart, C. et al. Transcript profiling of a xylem vs phloem cDNA subtractive library identifies new genes expressed during xylogenesis in Eucalyptus. New Phytol. 170, 739–752 (2006).

4. Kim, D. et al. TopHat2: accurate alignment of transcriptomes in the presence of insertions, deletions and gene fusions. Genome Biol. 14, 1–13 (2013).

5. Trapnell, C. et al. Transcript assembly and quantification by RNA-Seq reveals unannotated transcripts and isoform switching during cell differentiation. Nat. Biotechnol. 28, 511–515 (2010).

6. Love, M. I., Huber, W. & Anders, S. Moderated estimation of fold change and dispersion for RNA-seq data with DESeq2. Genome Biol. 15, 1–21 (2014).

7. Moura, J. C. M. S. et al. Validation of reference genes from Eucalyptus spp. under different stress conditions. BMC Res. Notes 5, 634 (2012).

8. Livak, K. J. & Schmittgen, T. D. Analysis of relative gene expression data using real-time quantitative PCR and the 2^−ΔΔCT^ method. methods 25, 402–408 (2001).

9. Bennett, M. J. et al. *Arabidopsis AUX1* gene: a permease-like regulator of root gravitropism. Science (1979) 273, 948–950 (1996).

10. Casanova-Sáez, R. et al. Inactivation of the entire Arabidopsis group II GH3s confers tolerance to salinity and water deficit. New Phytol. 235, 263–275 (2022).

11. Moreno-Risueno, M. A. et al. Oscillating gene expression determines competence for periodic *Arabidopsis* root branching. Science (1979) 329, 1306–1311 (2010).

12. Heisler, M. G. et al. Patterns of auxin transport and gene expression during primordium development revealed by live imaging of the *Arabidopsis* inflorescence meristem. Curr. Biol. 15, 1899–1911 (2005).

13. Rampey, R. A. et al. A family of auxin-conjugate hydrolases that contributes to free indole-3-acetic acid levels during Arabidopsis germination. Plant Physiol. 135, 978–988 (2004).

14. Schneider, C. A., Rasband, W. S. & Eliceiri, K. W. NIH Image to ImageJ: 25 years of image analysis. Nat. Methods 9, 671–675 (2012).

15. LeClere, S., Tellez, R., Rampey, R. A., Matsuda, S. P. T. & Bartel, B. Characterization of a family of IAA-amino acid conjugate hydrolases from *Arabidopsis*. J. Biol. Chem. 277, 20446–20452 (2002).

16. Bitto, E. et al. X-ray structure of ILL2, an auxin-conjugate amidohydrolase from *Arabidopsis thaliana*. Proteins 74, 61–71 (2009).

17. Sela, I., Ashkenazy, H., Katoh, K. & Pupko, T. GUIDANCE2: accurate detection of unreliable alignment regions accounting for the uncertainty of multiple parameters. Nucleic Acids Res. 43, W7–W14 (2015).

18. Trifinopoulos, J., Nguyen, L.-T., von Haeseler, A. & Minh, B. Q. W-IQ-TREE: a fast online phylogenetic tool for maximum likelihood analysis. Nucleic Acids Res. 44, W232–W235 (2016).

19. Yu, G. Smith, D.K. Zhu, H. Guan, Y. GGTREE: an R package for visualization and annotation of phylogenetic trees with their covariates and other associated data. Methods Ecol. Evo. 8, 28–36 (2017).

20. Wang, L.-G. et al. Treeio: an R package for phylogenetic tree input and output with richly annotated and associated data. Mol. Biol. Evol. 37, 599–603 (2020).

21. Labun, K. et al. CHOPCHOP v3: expanding the CRISPR web toolbox beyond genome editing. Nucleic Acids Res. 47, W171–W174 (2019).

22. Nekrasov, V., Staskawicz, B., Weigel, D., Jones, J. D. G. & Kamoun, S. Targeted mutagenesis in the model plant *Nicotiana benthamiana* using Cas9 RNA-guided endonuclease. Nat. Biotechnol. 31, 691–693 (2013).

23. Weber, E., Engler, C., Gruetzner, R., Werner, S. & Marillonnet, S. A modular cloning system for standardized assembly of multigene constructs. PLoS One 6, e16765 (2011).

24. Grützner, R. et al. High-efficiency genome editing in plants mediated by a Cas9 gene containing multiple introns. Plant Commun. 2, 100135 (2021).

25. Karimi, M., Inzé, D. & Depicker, A. GATEWAY^TM^ vectors for *Agrobacterium*-mediated plant transformation. Trends Plant Sci. 7, 193–195 (2002).

26. Zhang, X., Henriques, R., Lin, S.-S., Niu, Q.-W. & Chua, N.-H. *Agrobacterium*-mediated transformation of *Arabidopsis thaliana* using the floral dip method. Nat. Protoc. 1, 641–646 (2006).

27. Prigge, M. J., Lavy, M., Ashton, N. W. & Estelle, M. *Physcomitrella patens* auxin-resistant mutants affect conserved elements of an auxin-signaling pathway. Curr. Biol. 20, 1907–1912 (2010).

28. Gietz, R. D. & Schiestl, R. H. High-efficiency yeast transformation using the LiAc/SS carrier DNA/PEG method. Nat Protoc. 2, 31–34 (2007).

29. Pierre-Jerome, E., Jang, S. S., Havens, K. A., Nemhauser, J. L. & Klavins, E. Recapitulation of the forward nuclear auxin response pathway in yeast. Proc. Natl. Acad. Sci. U.S.A 111, 9407–9412 (2014).

30. Prusinska, J. et al. The differential binding and biological efficacy of auxin herbicides. Pest Manag. Sci. (2022).

31. Lee, S. et al. Defining binding efficiency and specificity of auxins for SCFTIR1/AFB-Aux/IAA co-receptor complex formation. ACS Chem. Biol. 9, 673–682 (2014).

32. Fastner, A., Absmanner, B. & Hammes, U. Z. Use of *Xenopus laevis* oocytes to study auxin transport. In Plant hormones 259–270 (Springer, 2017).

33. Frank, O., Kreissl, J. K., Daschner, A. & Hofmann, T. Accurate determination of reference materials and natural isolates by means of quantitative ^1^H NMR spectroscopy. J. Agric. Food Chem. 62, 2506–2515 (2014).

34. Ung, K. L. et al. Structures and mechanism of the plant PIN-FORMED auxin transporter. Nature 609, 605–610 (2022).

35. Friesner, R. A. et al. Extra precision glide: docking and scoring incorporating a model of hydrophobic enclosure for protein−ligand complexes. J. Med. Chem. 49, 6177–6196 (2006).

36. Halgren, T. A. et al. Glide: a new approach for rapid, accurate docking and scoring. 2. Enrichment factors in database screening. J. Med. Chem. 47, 1750–1759 (2004).

37. Friesner, R. A. et al. Glide: a new approach for rapid, accurate docking and scoring. 1. Method and assessment of docking accuracy. J. Med. Chem. 47, 1739–1749 (2004).

38. Jumper, J. et al. Highly accurate protein structure prediction with AlphaFold. Nature 596, 583–589 (2021).

39. Madhavi Sastry, G., Adzhigirey, M., Day, T., Annabhimoju, R. & Sherman, W. Protein and ligand preparation: parameters, protocols, and influence on virtual screening enrichments. J. Comput. Aided Mol. Des. 27, 221–234 (2013).

40. Søndergaard, C. R., Olsson, M. H. M., Rostkowski, M. & Jensen, J. H. Improved treatment of ligands and coupling effects in empirical calculation and rationalization of p*K*_a_ values. J. Chem. Theory Comput. 7, 2284–2295 (2011).

41. Olsson, M. H. M., Søndergaard, C. R., Rostkowski, M. & Jensen, J. H. PROPKA3: consistent treatment of internal and surface residues in empirical p*K*_a_ predictions. J. Chem. Theory Comput. 7, 525–537 (2011).

42. Greenwood, J. R., Calkins, D., Sullivan, A. P. & Shelley, J. C. Towards the comprehensive, rapid, and accurate prediction of the favorable tautomeric states of drug-like molecules in aqueous solution. J. Comput. Aided Mol. Des. 24, 591–604 (2010).

43. Pan, X., Wang, H., Li, C., Zhang, J. Z. H. & Ji, C. MolGpka: a web server for small molecule p*K*_a_ prediction using a graph-convolutional neural network. J. Chem. Inf. Model. 61, 3159–3165 (2021).

