## Supplementary Information for "Slow release of a synthetic auxin induces formation of adventitious roots in recalcitrant woody plants"

**Table of Contents**


1. Supplementary Figures and Tables…………..…………..…………………………S2
2. $^{1}$H- and $^{13}$C-NMR and mass-spectrometry data…………………...…………...……...S19
3. $^{1}$H- and $^{13}$C-NMR spectra……………………………………..……………………..S29

### Supplementary Figures and Tables

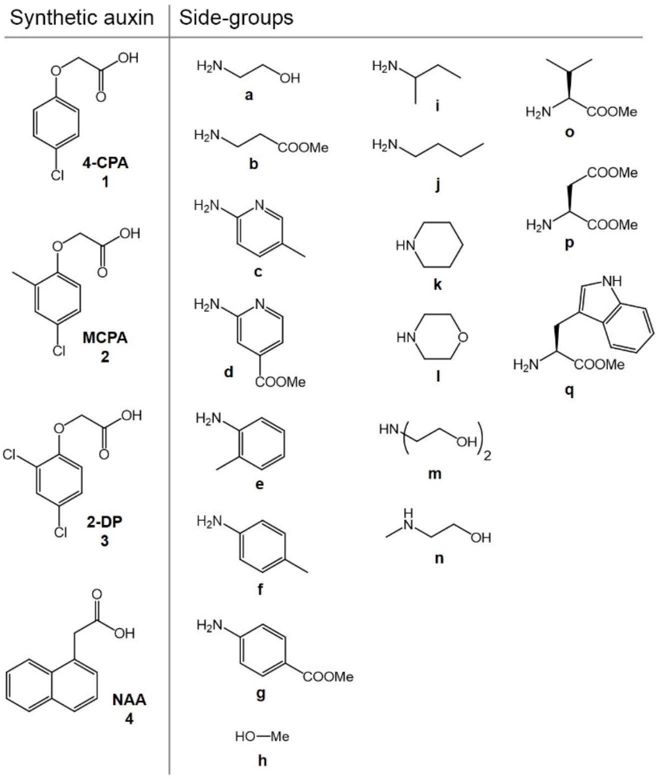


**Figure 1. Chemical Structures of the synthetic auxins and side-groups used for conjugation.**

The conjugates were synthesized in three sets: first, all four synthetic auxins (**1**–**4**) were conjugated to molecules **a**–**h** forming conjugates **1**–**4a**–**h**. Their evaluation on *E. grandis* cuttings showed that only 1a significantly enhanced rooting when combined with 6,000 ppm K-IBA and that from the four top performers, three where conjugates of 4-CPA (**1a**, **1b** and **1g**, up to 25% rooting, see Supplementary Fig. 2). Therefore, a second set of conjugates was synthesized based on 4-CPA and moieties structurally resembling a (conjugates **1i**–**m**). On average, this set of conjugates engendered higher rooting percentages in *E. grandis* cuttings when combined with K-IBA (up to 30% rooting, see Supplementary Fig. 2). A structure-activity relationship (SAR) analysis of all 4-CPA conjugates (**1a**–**n**) showed that conjugates of primary amines with p$K_a$ ~8.5–10.5 had the highest rooting promoting activity in this assay. We therefore considered amino acids, which bear a primary amine with a p$K_a$ of ~9.6 as potential conjugation partners to 4-CPA. Thus, conjugates of 4-CPA with three amino acids, L-valine (Val, **o**), L-aspartate (Asp, **p**) and L-tryptophan (Trp, **q**) in their methyl ester form were synthesized (**1o**–**1q**).

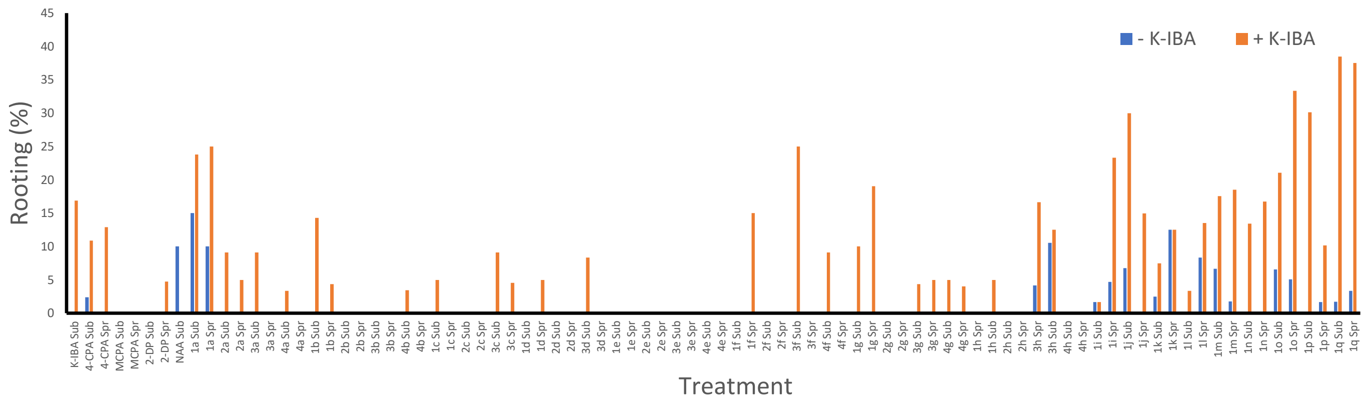


**Figure 2. Chemical screen for rooting enhancers in *E. grandis*.**

Rooting percentages in response to application of the indicated compounds (100 µM) by submerging (Sub) the cutting base for 1 minute, or spraying (Spr) the leaves, in the absence (blue) or in the presence (orange) of 6,000 ppm K-IBA in the submerging solution. Rooting was evaluated 1 month from application. n > 500 for K-IBA, n > 20 for conjugate-based application.

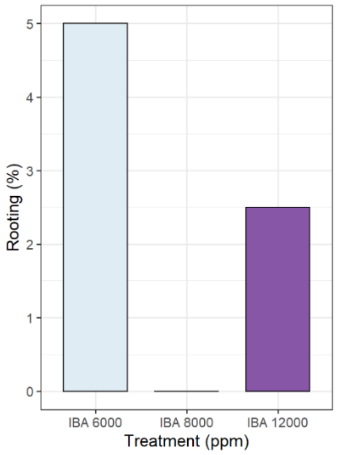


**Figure 3. Increasing K-IBA concentration does not improve rooting efficiency of *E. grandis* mature cuttings.**

Rooting percentages of mature *E. grandis* cuttings in response to K-IBA submergence treatment at the indicated concentrations. Rooting was evaluated 1 month from application. n = 40 cuttings per sample.

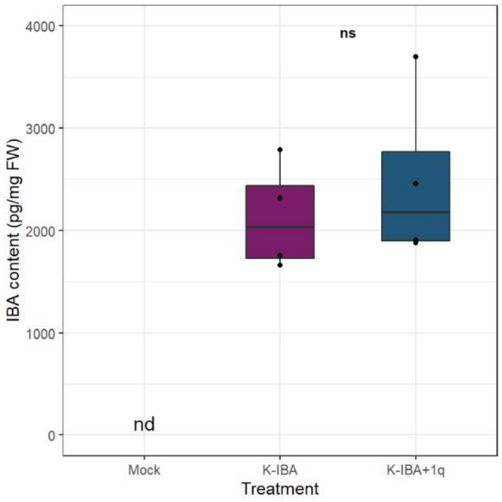


**Figure 4. Conjugate-K-IBA co-application does not increase IBA uptake in *E. grandis* mature cuttings.**

Cutting bases were submerged for 1 min with mock (0.1% DMSO), K-IBA (6,000 ppm) or K-IBA (6,000 ppm) + 1q (100 µM) and planted in a rooting table for 15 min. Each dot represents a sample of 10 cuttings. n = 4 biological replicates. nd stands for not detected. ns stands for not significant according to two-sided Student's t test. Box-plot elements definition - center line, median; box limits, upper and lower quartiles; whiskers, 1.5x interquartile range.

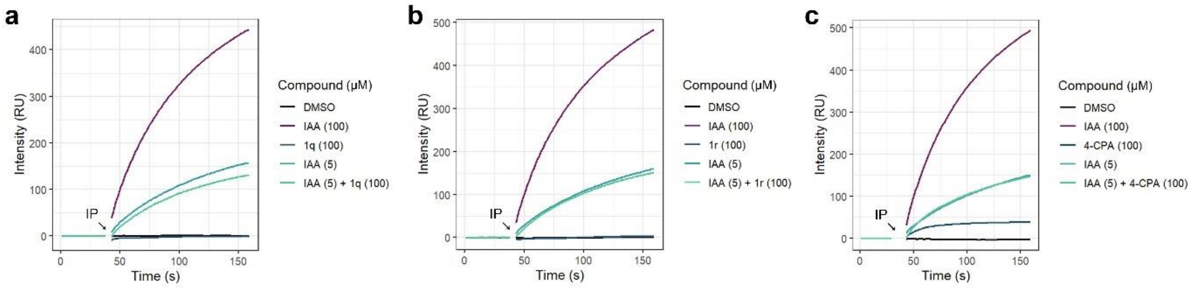


**Figure 5. Examinations for anti-IAA activity.**

**a-c,** SPR assay testing anti-IAA activity of **1q** (**a**), **1r** (**b**), or 4-CPA (**c**), using TIR1 and IAA7 degron domain. IP stands for injection point of TIR1 mixed with the indicated compounds in solution, RU stands for resonance unit.

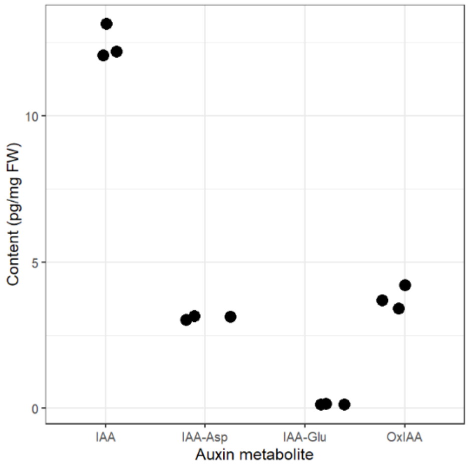


**Figure 6. Endogenous auxin and auxin metabolites levels in mature *E. grandis* cuttings.**

Replicates were extracted from a pool of 20 cutting-bases harvested and grinded together. Asp and Glu stand for aspartate and glutamate, respectively.

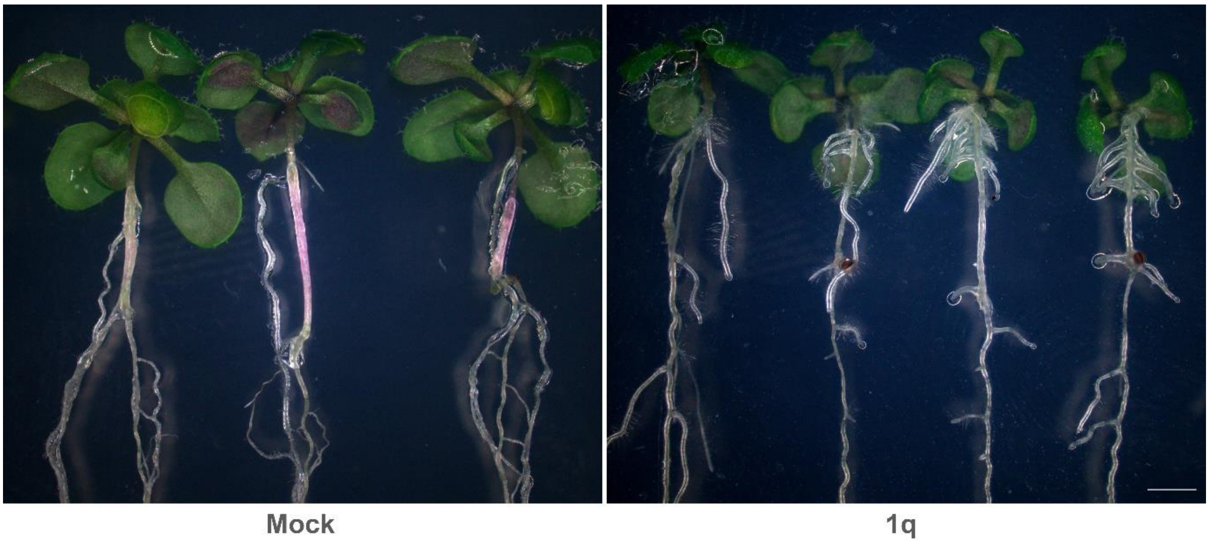


**Figure 7. Representative images of mock or 1q (10 µM) induced AR in *Arabidopsis*.** The treatments were applied for 1.5 h to shoots of 3-day-old etiolated seedlings, and images were taken two weeks later. Bar = 2 mm. Of note, these images were taken from a different experiment than the one described in Fig 2e in the main text.

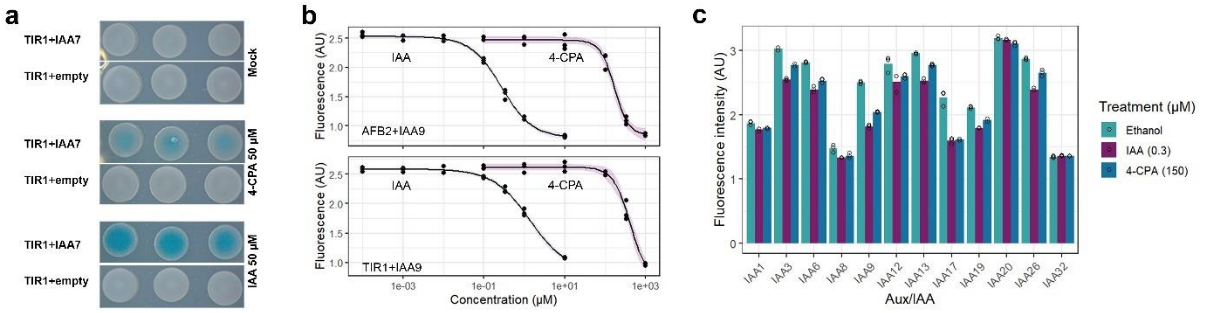


**Figure 8. 4-CPA is a weaker receptors binder compared to IAA.**

**a**, Qualitative Y2H to detect auxin activity, using TIR1 receptor (bait) and IAA7 degron (prey) and auxin at concentration shown. Three colonies per strain and treatment are shown. DMSO (0.1%) was used as Mock. **b**, Quantitative Y2H to detect auxin activity. Curve response testing fluorescence signal of YFP-tagged degron (IAA9) with AFB2 (up) or TIR1 (down) to increasing concentrations of IAA (black) or 4-CPA (pink). n = 3 biological replicates. **c**, Quantitative Y2H assay using TIR1 and YFP-tagged Aux/IAAs and the indicated auxin or Ethanol (0.1%) as mock. IAA and 4-CPA concentrations are $EC_{50}$ values calculated from the data presented in **b**. n = 3 biological replicates. Shown are means of fluorescence intensity. Error-bars represent standard errors.

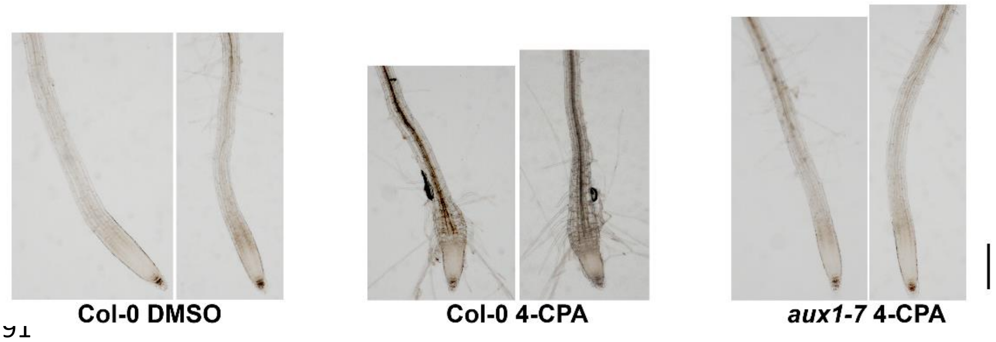


**Figure 9. Shoot application of 4-CPA inhibits root growth, in AUX1 dependent manner.**

Stereoscope images of *Arabidopsis* root tips 3 days after etiolated-shoot application (in a split-dish experiment) of 4-CPA (10 µM) or DMSO (0.1%) for 6 hours. Bar = 0.2 mm.

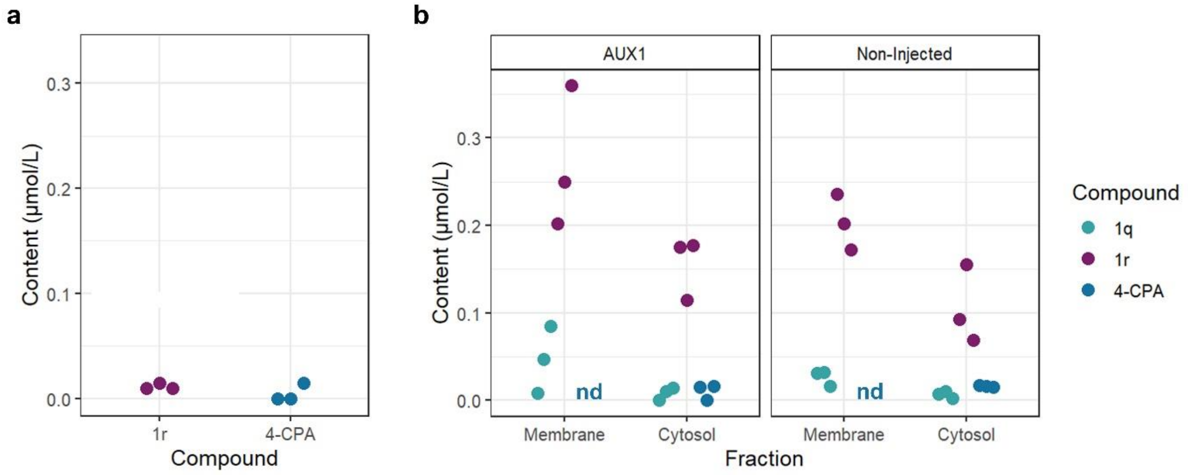


**Figure 10. AUX1 does not import 1r and 1q into *Xenopus laevis* oocytes.**

**a**, HPLC-MS/MS quantification of cytosolic **1r** and 4-CPA of AUX1-expressing oocytes following 30 min incubation with **1r**, n = 3. **b**, HPLC-MS/MS quantification of **1q**, **1r** and 4-CPA in cytosolic or membrane fractions of AUX1-expressing (left) or non-expressing (right) oocytes following 30 min incubation with **1q**. nd stands for not detected, n = 3.

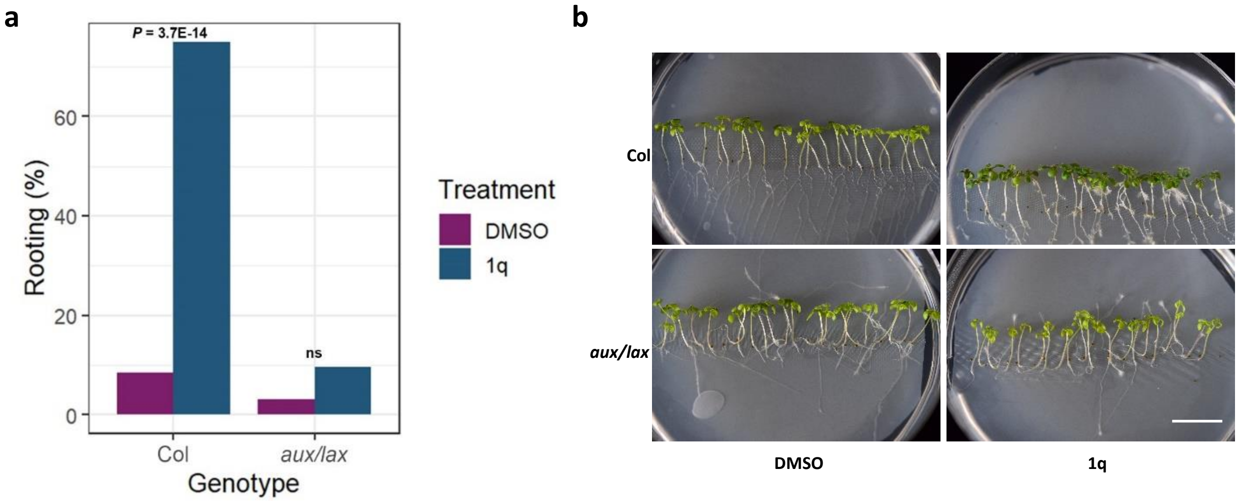


**Figure 11. AUX/LAX importers are required for 1q rooting activity.**

**a,** Percentages of two-week-seedlings (*Arabidopsis*) that developed adventitious roots in response to 10 µM **1q** or 0.1% for DMSO applied specifically to shoots for 1.5 h via a split-dish). Shown are p-values of Fisher's exact test testing the hypothesis that **1q** treatment results with higher rooting percentages (a one-sided test), ns stands for not significant. n = 71 (Col + DMSO), 48 (Col + **1q**), 64 (*aux/lax* + DMSO), and 62 (*aux/lax* + **1q**). **b**, Representative images of plants 2 weeks after induction, bar = 1.5 cm.

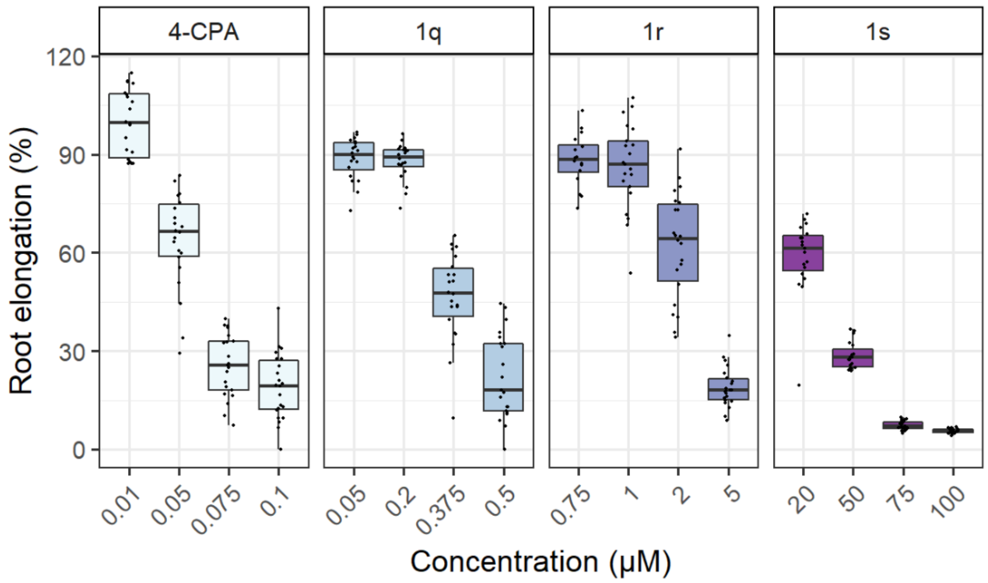


**Figure 12.** ***Arabidopsis*** **root elongation curve response to 4-CPA, 4-CPA-L-tryptophan-OMe (1q), 4-CPA-L-tryptophan (1r), and 4-CPA-D-tryptophan-OMe (1s).**

Presented is relative root elongation after 3-days incubation with the indicated compound. DMSO (0.1%) was used as mock. n > 16 plants per sample. Box-plot elements definition - center line, median; box limits, upper and lower quartiles; whiskers, 1.5x interquartile range.

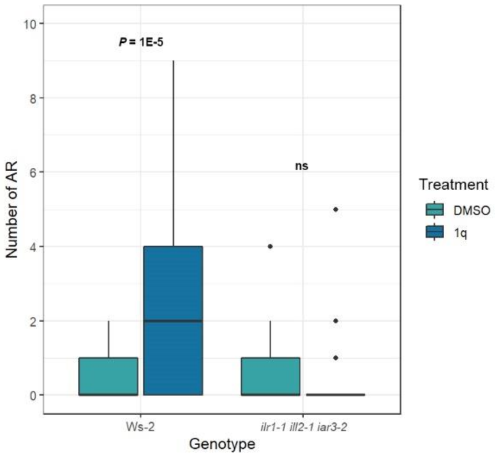


**Figure 13. Number of adventitious roots in response to 1q induction.**

Shoot specific application of 10 µM for **1q** or 0.1% DMSO in a split-dish for 1.5 h. ARs were quantified 14 d after treatment. n > 43 plants per sample. Two-sided Mann-Whitney U test p-value is presented. ns stands for not significant. Box-plot elements definition - center line, median; box limits, upper and lower quartiles; whiskers, 1.5x interquartile range; points, outliers.

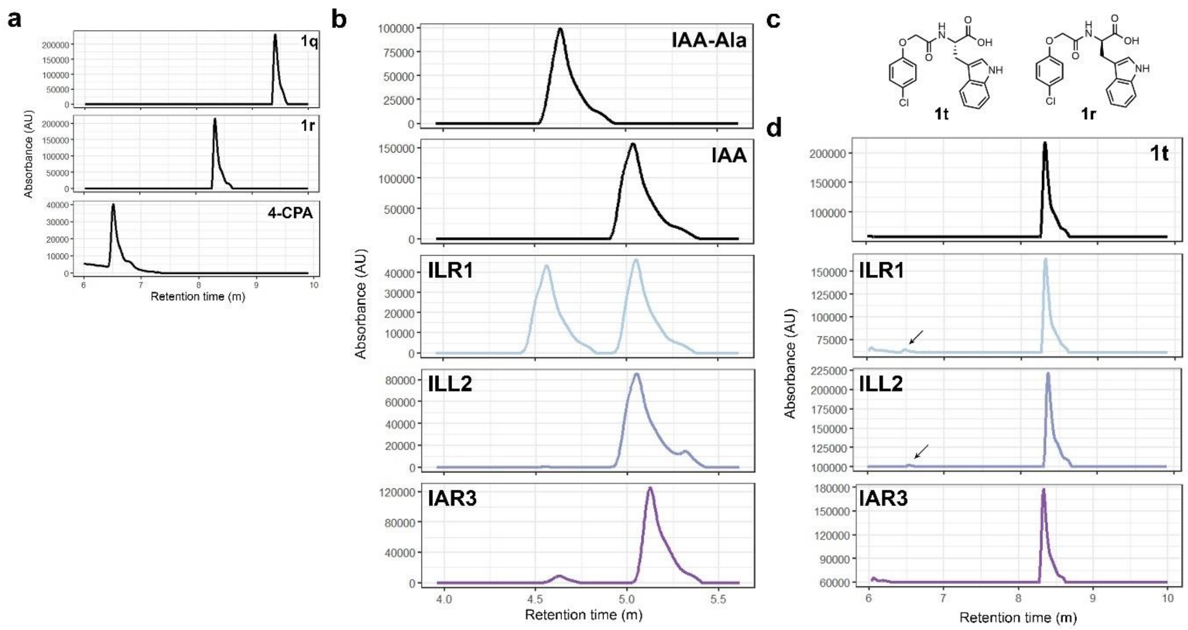


**Figure 14. *In-vitro* evaluation of amidohydrolases activities.**

**a**, HPLC-MS chromatographs of the indicated compounds. **b**, Hydrolytic activity of the indicated enzymes against IAA-Ala. This assay severs as control for enzymatic activity. **c**, Chemical structures of the indicated enantiomers **1t** and **1r**. **d**, Evaluation of hydrolytic activity towards **1t**.

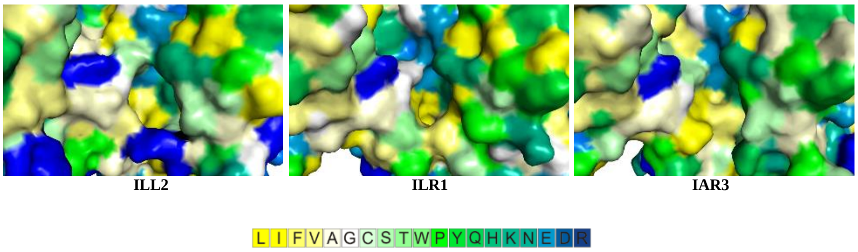


**Figure 15. Ligand binding pocket of the tested amidohydrolases.**

Amino acids are color-coded according to the Kessel/Ben-Tal hydrophobicity scale[1] (ranging from most hydrophobic (yellow) to most hydrophilic (blue)).

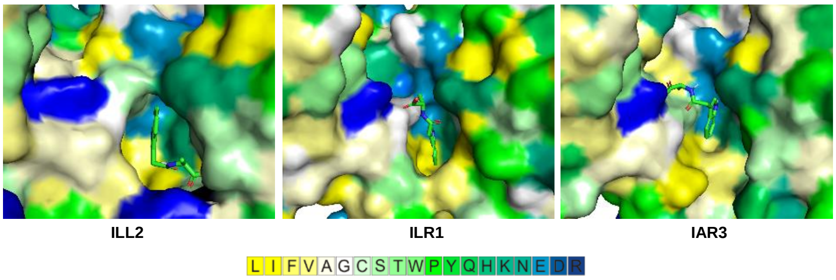


**Figure 16. Docking calculations of the indicated amidohydrolase with IAA-Ala.**

Amino acids are color-coded according to residues hydrophobicity, as described in Supporting Fig. 12.

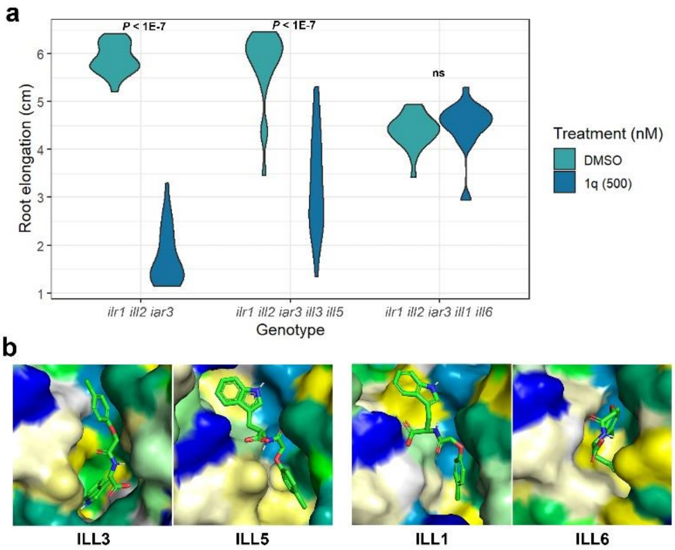


**Figure 17. Characterization of *ilr1/ills* higher order mutants.**

**a**, Root elongation of *ilr1-1 ill2-1 iar3-2*, *ilr1-1 ill2-1 iar3-2 ill3-1 ill5-1*, and *ilr1-1 ill2-1 iar3-2 ill1-1 ill6-1* after 7 days of treatment. n > 22 plants per sample. Two-sided Tukey's HSD p-values are presented. **b**, Docking calculations of **1r** with the indicated enzymes. Amino acids are color-coded according to residues hydrophobicity, as described in Supporting Fig. 12.

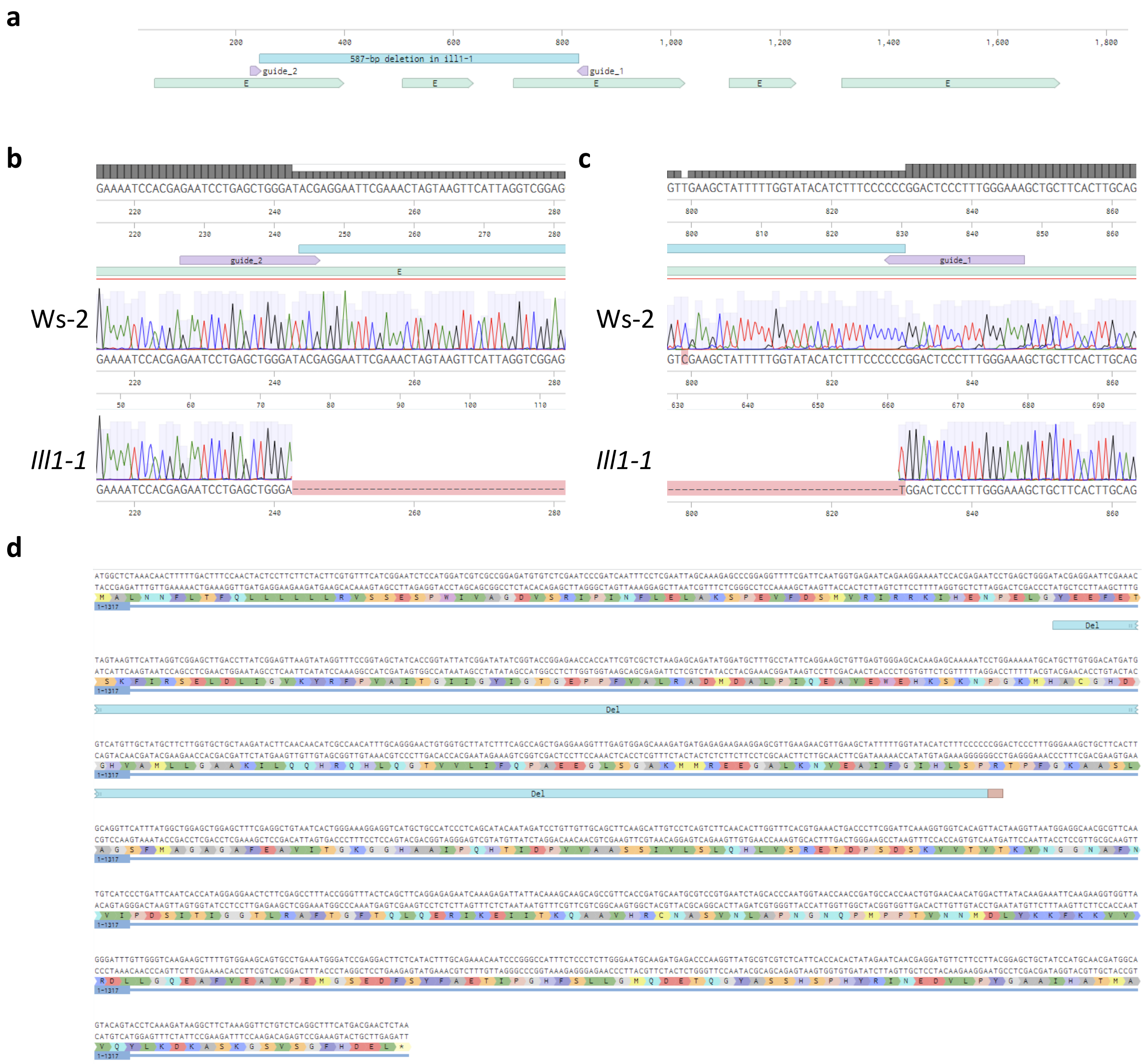


**Figure 18. Characterization of the mutated *ill1* allele generated with CRISPR-Cas9.**

**a**, Illustration of *ILL1* gene annotated with the deletion in *ill1-1* (cyan), and the guides to target Cas9 (purple). E stands for exons. **b-c**, Sanger-sequence chromatogram of Ws-2 and *ill1-1* aligned to Col-0 reference genome showing the deletion 5' (**b**) and 3' (**c**). **d**, Protein sequence of ILL1 (Col-0) annotated with the deletion (cyan) and R-to-W substitution (brown) in *ill1-1*.

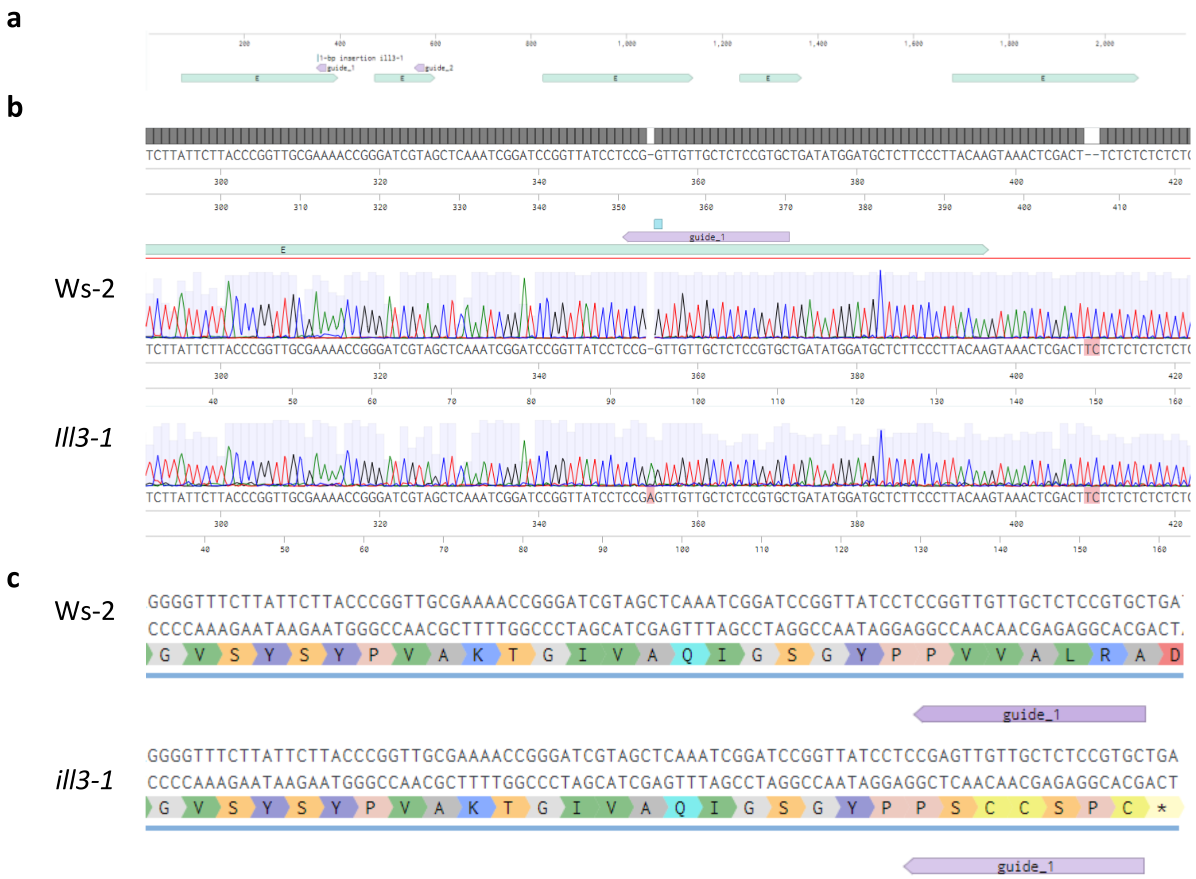


**Figure 19. Characterization of the mutated *ill3* allele generated with CRISPR-Cas9.**

**a**, Illustration of *ILL3* gene annotated with the 1-bp insertion in *ill3-1* (cyan), and the guides to target Cas9 (purple). E stands for exons. **b**, Sanger-sequence chromatogram of Ws-2 and *ill1-3* aligned to Col-0 reference genome showing 1-bp insertion. **c**, Protein sequences of ILL3 in Ws-2 and *ill3-1*, showing an early stop-codon in *ill3-1* as a result of a frameshift mutation.

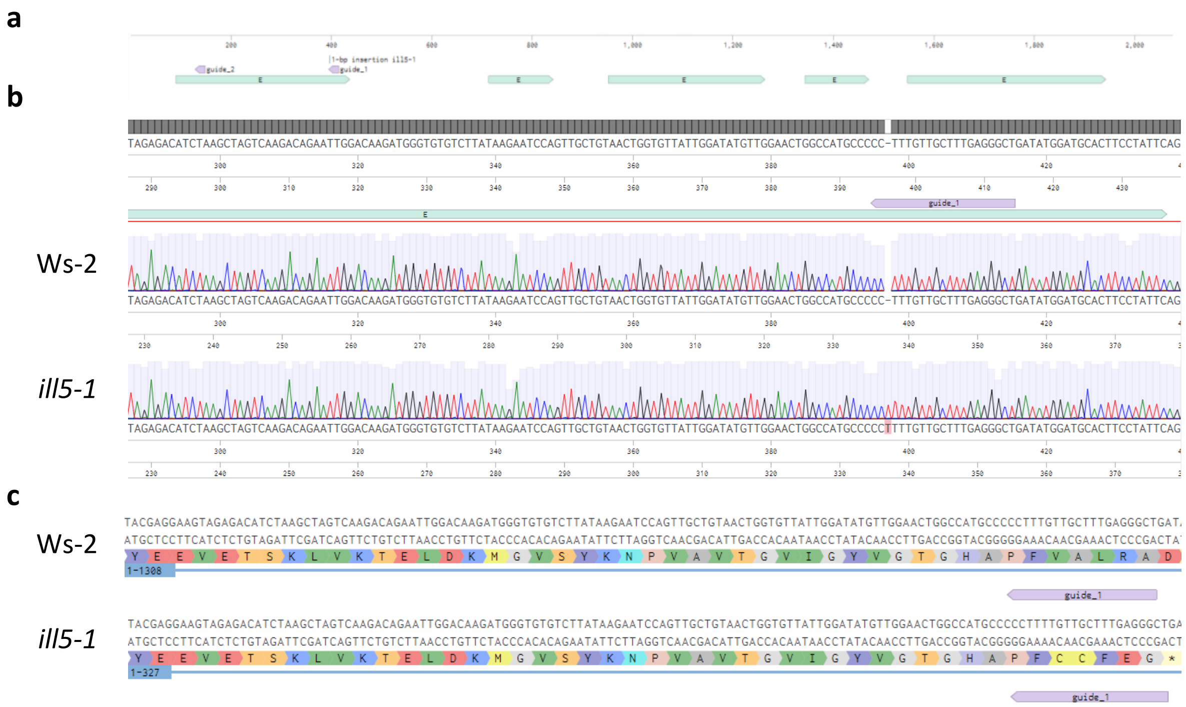


**Figure 20. Characterization of the mutated *ill5* allele generated with CRISPR-Cas9.**

**a**, Illustration of *ILL5* gene annotated with the 1-bp insertion in *ill5-1* (cyan), and the guides to target Cas9 (purple). E stands for exons. **b**, Sanger-sequence chromatogram of Ws-2 and *ill5-1* aligned to Col-0 reference genome showing 1-bp insertion. **c**, Protein sequences of ILL5 in Ws-2 and *ill5-1*, showing an early stop-codon in *ill5-1* as a result of a frameshift mutation.

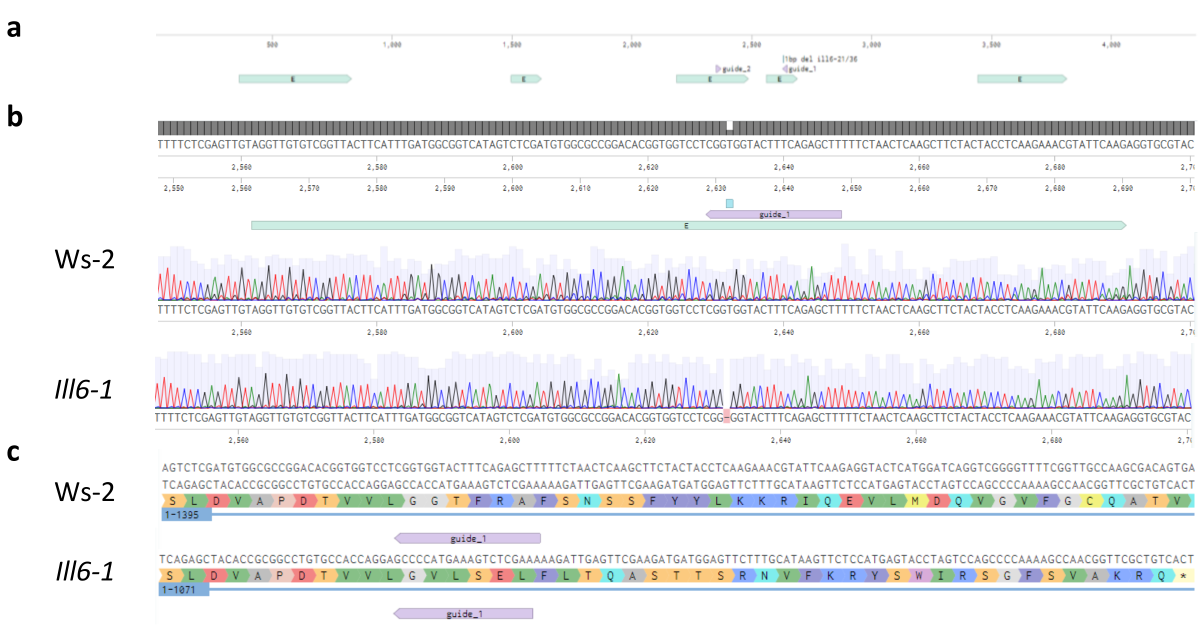


**Figure 21. Characterization of the mutated *ill6* allele generated with CRISPR-Cas9.**

**a**, Illustration of *ILL6* gene annotated with the 1-bp insertion as appeared in *ill6-1* (cyan), and the guides to target Cas9 (purple). E stands for exons. **b**, Sanger-sequence chromatogram of Ws-2 and *ill6-1* aligned to Col-0 reference genome showing 1-bp deletion. **c**, Protein sequences of ILL6 in Ws-2 and *ill6-1*, showing an early stop-codon in *ill6-1* as a result of a frameshift mutation.

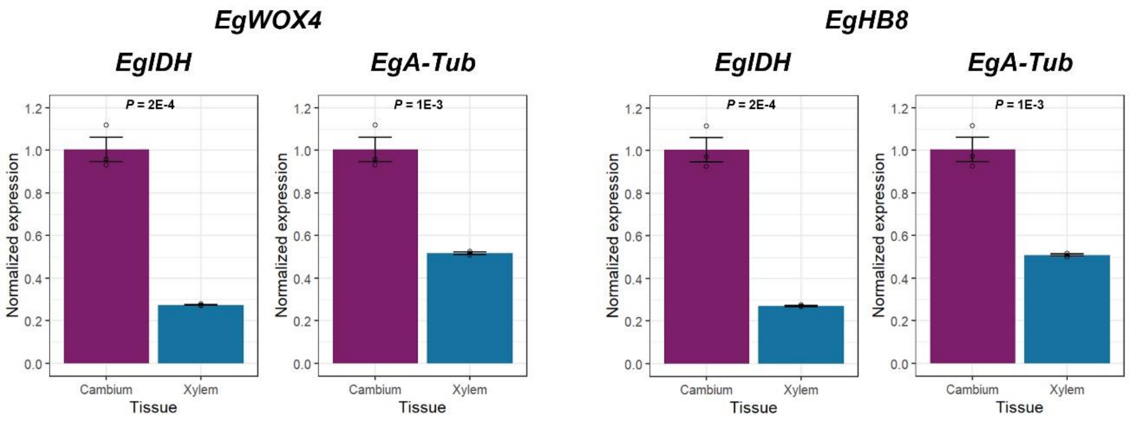


**Figure 22. Validation of the cambium enrichment procedure.**

Quantitative real-time PCR assay examining the expression of two cambium markers; *EgWOX4* (*left*) and *EgHB8* (*right*), in manually harvested cambium or xylem tissues. *EgIDH* and *EgA-Tub* were used to normalize expression. Two-sided Student's t test p-values are shown, n = 3 technical replicates. Shown are means of normalized expression. Error-bars represent standard errors.

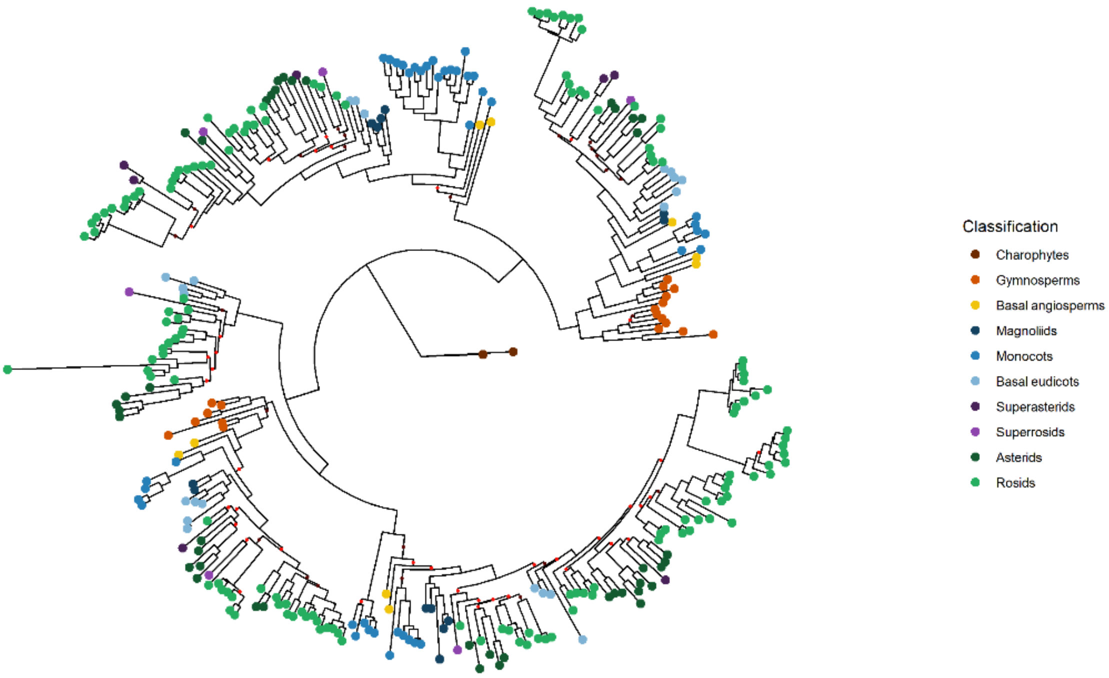


**Figure 23. Alternative phylogenetic tree based on MSA generated with automatic MAFFT strategy.**

Branches are annotated in brown or red for bootstrap values lower than 85 or 70, respectively.

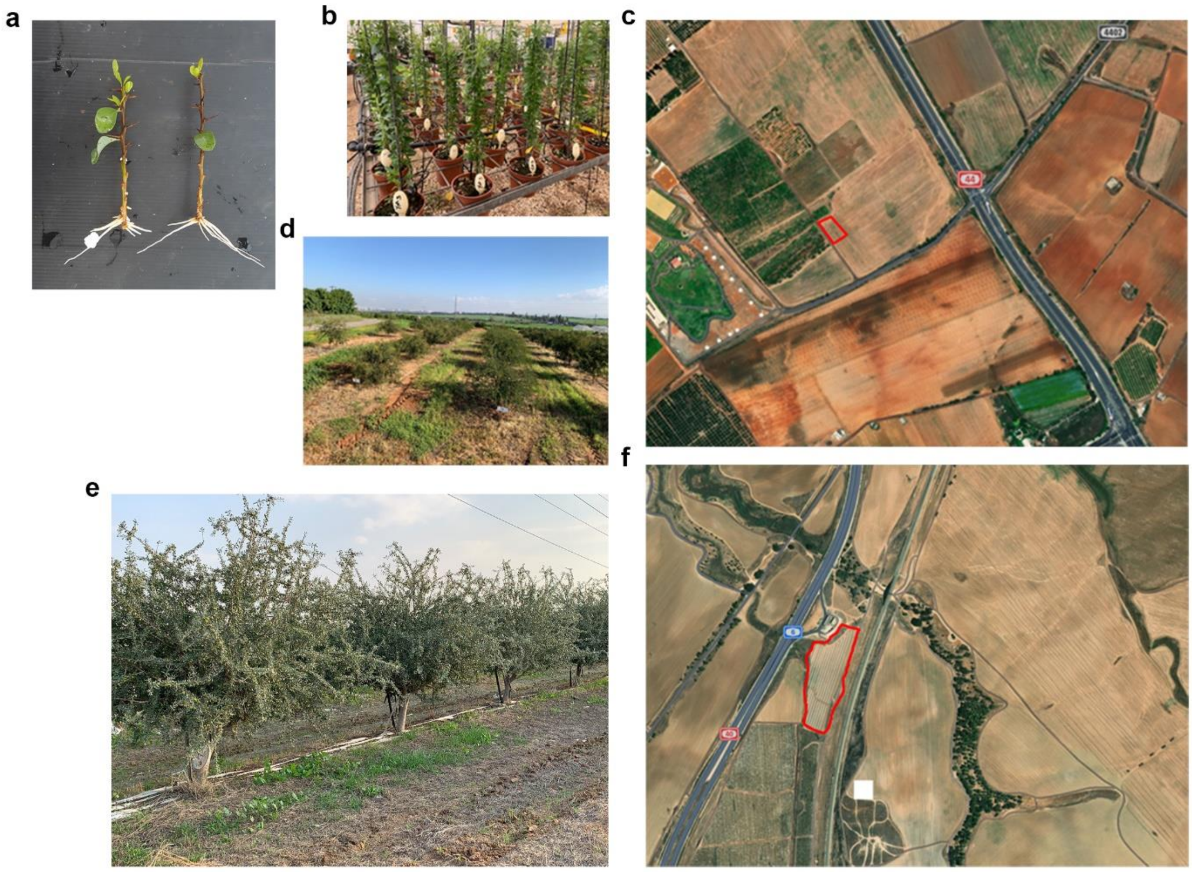


**Figure 24. Propagation of selected argan clones.**

A survey of historic argan plantations in Israel revealed that only ~10% of the trees provide yields at an economically viable level. Cuttings from select trees were obtained and propagated. **a**, A picture of rooted argan cuttings from the select trees treated with K-IBA + **1q** after 2 months in a rooting table. **b**, A picture of rooted argan clones grown in the greenhouse at Volcani Center (plants are ca. 8 months old). **c**, An aerial image of an experimental plot of argan clones established in Volcani Center (2021). The plot includes 19 unique clones in 5 replicates. **d**, The trees in **c**, 1.5 years after plantation. **e**, A picture of an earlier argan plot (planted on 2019) which includes 7 clones and planted in Beit-Kama, Israel. Picture was taken in Nov. 2021. **f**, An aerial image of the Beit-Kama plot which includes 250 Argan trees of the 7 clones.

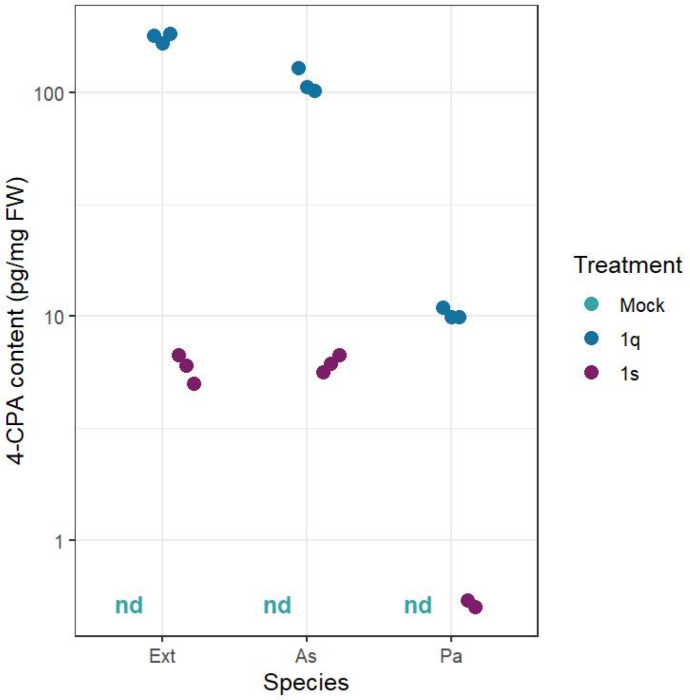


**Figure 25. 4-CPA release from 1q is enzymatically regulated in cuttings of *Eucalyptus x trabutii* (Ext), *Argania spinosa* ARS7 line (As), and *Populus alba* (Pa).** HPLC-MS/MS quantification of 4-CPA 24 h after basal application with the indicated enantiomer (100 µM) + K-IBA basal treatment (6,000 ppm), or K-IBA (6,000 ppm) + DMSO (0.1%) as mock. Each sample is composed of 3 replicates, extracted from a pool of 20 cutting-bases harvested together. Data presented in logarithmic scale. nd stands for not detected. Of note, 4-CPA was not detected in one **1s** replicate.

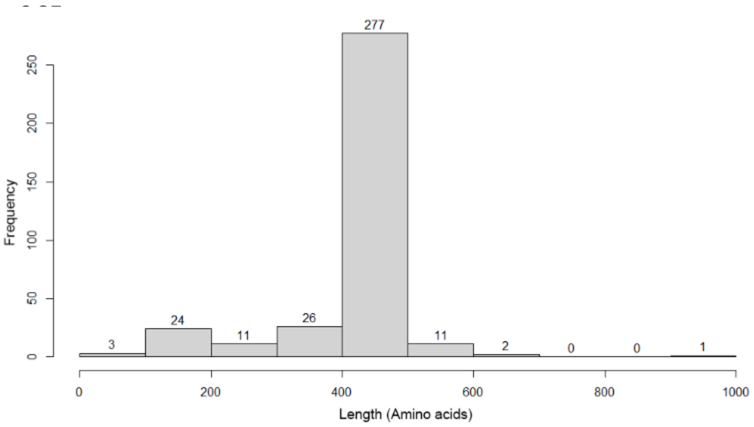


**Figure 26. Distribution of the obtained 355 ILR1/ILL sequences as a function of proteins length.** The majority of the sequences (78%) are of 400–500 amino acids. Sequences of 350–550 amino acids were selected for phylogenetic analysis.

### Table 1. Primers used for qPCR in *E. grandis*

| Name | Sequence |
|---|---|
| EgIDH_F | TGGAACTGTTGAGTCTGG |
| EgIDH_R | TTAGGACCATGAATGAGGAG |
| Tubulin_F | TCGCGCTGTGTTTGTGGATCTG |
| Tubulin_R | GCTGCTCAGGGTGAAAGAGCTG |
| HB8_F | ATGGAGGAGAACGATAGGCTGC |
| HB8_R | GGTTGCGTTTTGTGTCTGCTGG |
| WOX4_F | GGCAGAAGCAGAAGCGCAACAG |
| WOX4_R | AAAGTTACAGTGGCGATGGTGG |

### Table 2. Target sequences used to clone sgDNA against ILL1,3,5 and 6.

| sgDNA Name | Location | sgDNA sequence | Primer sequence to generate Golden Gate compatible sgDNA together with – tgtggtctcaAGCGTAATGCCAACTTTGTAC as reverse |
|---|---|---|---|
| ILL6_1 | 3rd exon | AAGCTCTGAAAGTACCACCG | tgtggtctcaATTGAGCTCTGAAAGTACCACCGgttttagagctagaaatagcaag |
| ILL6_2 | 4th exon | GGGATGTGGAATTTTCCGGG | tgtggtctcaATTGGGATGTGGAATTTTCCGGGgttttagagctagaaatagcaag |
| ILL1_1 | 3rd exon | TTTCCCAAAGGGAGTCCGGG | tgtggtctcaATTGTTCCCAAAGGGAGTCCGGGgttttagagctagaaatagcaag |
| ILL1_2 | 1st exon | GAATCCTGAGCTGGGATACG | tgtggtctcaATTGAATCCTGAGCTGGGATACGgttttagagctagaaatagcaag |
| ILL5_1 | 1st exon | AGCCCTCAAAGCAACAAAGG | tgtggtctcaATTGGCCCTCAAAGCAACAAAGGgttttagagctagaaatagcaag |
| ILL5_2 | 1st exon | GACACGAGTTTAGTAAGTGA | tgtggtctcaATTGACACGAGTTTAGTAAGTGAgttttagagctagaaatagcaag |
| ILL3_1 | 1st exon | GCACGGAGAGCAACAACCGG | tgtggtctcaATTGCACGGAGAGCAACAACCGGgttttagagctagaaatagcaag |
| ILL3_2 | 2nd exon | TAAGCAATTTAGCAGCACCA | tgtggtctcaATTGAAGCAATTTAGCAGCACCAgttttagagctagaaatagcaag |

### Table 3. Synthetic methods and yields.

| Compd # | Method | Yield (%) | Compd # | Method | Yield (%) |
|---|---|---|---|---|---|
| **1a** | A | 96 | **4f** | B | 50 |
| **2a** | A | 81 | **1g** | A | 45 |
| **3a** | A | 92 | **2g** | A | 46 |
| **4a** | B | 83 | **3g** | A | 57 |
| **1b** | A | 73 | **4g** | B | 56 |
| **2b** | A | 68 | **1h** | C | 97 |
| **3b** | A | 56 | **2h** | C | 97 |
| **4b** | B | 61 | **3h** | C | 91 |
| **1c** | A | 77 | **4h** | C | 83 |
| **2c** | A | 74 | **1i** | A | 90 |
| **3c** | A | 43 | **1j** | A | 85 |
| **1d** | A | 35 | **1k** | A | 87 |
| **2d** | A | 44 | **1l** | A | 76 |
| **3d** | A | 54 | **1m** | A | 95 |
| **1e** | A | 56 | **1n** | A | 85 |
| **2e** | A | 63 | **1o** | A | 95 |
| **3e** | A | 64 | **1p** | A | 76 |
| **4e** | B | 47 | **1q** | A | 88 |
| **1f** | A | 77 | **1s** | A | 90 |
| **2f** | A | 65 | | | |
| **3f** | A | 75 | | | |

### $^{1}H$- and $^{13}C$-NMR and mass-spectrometry data

#### Compound 1a

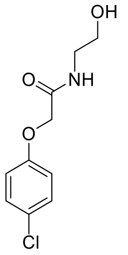

$^{1}H$ NMR (400 MHz, $CDCl_3$): δ, 7.28 (d, *J* = 9.2 Hz 2H), 7.00 (s, 1H), 6.86 (d, *J* = 9.0 Hz, 2H), 4.48 (s, 2H), 3.77 (d, *J* = 5.0 Hz, 2H), 3.53 (q, *J* = 5.3 Hz, 2H); $^{13}C$ NMR (101 MHz, $CDCl_3$): δ 168.9, 155.7, 129.7, 127.2, 116.0, 67.5, 61.8, 41.8. HR-MS(ESI) calcd. for formula $C_{10}H_{12}ClNO_3Na[M+Na]^+$: 252.0403; Found: 252.0408.

#### Compound 2a

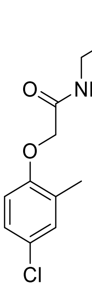

$^{1}H$ NMR (400 MHz, $CDCl_3$): δ 7.11–7.15 (m, 2H), 6.99 (br S, 1H), 6.68 (d, *J* = 8.6 Hz, 1H), 4.48 (s, 2H), 3.76 (q, *J* = 4.8 Hz, 2H), 3.52 (d, *J* = 5.3 Hz, 2H), 2.51 (t, *J* = 4.9 Hz, 1H), 2.25 (s, 3H); $^{13}C$ NMR (101 MHz, $CDCl_3$): δ 169.2, 154.1, 131.0, 128.6, 126.9 (2), 112.8, 67.8, 62.1, 41.9, 16.4. HR-MS(ESI) calcd. for formula $C_{11}H_{14}ClNO_3Na[M+Na]^+$: 266.0560; Found: 266.0557.

#### Compound 3a

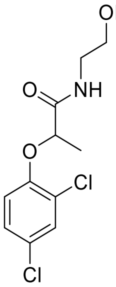

$^{1}H$ NMR (400 MHz, $CDCl_3$): δ 7.39 (d, J = 2.5 Hz, 1H), 7.18 (dd, J = 2.6 and *J* = 8.8 Hz, 1H), 7.13 (br S, 1H), 6.84 (d, *J* = 8.8 Hz, 1H), 4.68 (q, *J* = 6.7 Hz, 1H), 3.73 (t, *J* = 4.8 Hz, 2H), 3.50 - 3.45 (m, 2H), 2.52 (s, 1H), 1.62 (d, *J* = 6.8 Hz, 3H); $^{13}C$ NMR (101 MHz, $CDCl_3$): δ 172.3 ,151.4, 130.5, 128.1, 127.7, 124.87, 116.3, 62.2, 42.2, 18,6. HR-MS(ESI) calcd. for formula $C_{11}H_{13}Cl_2NO_3Na[M+Na]^+$: 300.0170; Found: 300.0169.

#### Compound 4a

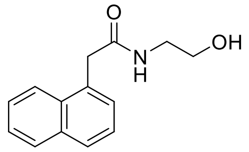

$^{1}H$ NMR (400 MHz, $CDCl_3$): δ 7.83–7.96 (m, 3H), 7.41 – 7.58 (m, 4H), 5.71 (br s, 1H), 4.06 (s, 2H), 3.57 (t, *J* = 4.4 Hz, 2H), 3.28 (q, *J* = 5.2 Hz, 2H), 2.41 (s, 1H); $^{13}C$ NMR (101 MHz, $CDCl_3$): δ 172.4, 134.0, 132.0, 130.8, 128.9, 128.6, 128.5, 126.9, 126.2, 125.7, 123.6, 62.3, 42.7, 41.6. HR-MS(ESI) calcd. for formula $C_{14}H_{15}NO_2Na[M+Na]^+$: 252.1000; Found: 252.0998.

**Compound 1b** 296

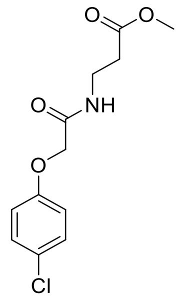


$^1$H NMR (400 MHz, $CDCl_3$): δ 7.24 (d, *J* = 9.0 Hz, 2H), 7.09 (br s, 1H), 6.83 (d, *J* = 9.0 Hz, 2H), 4.44 (s, 2H), 3.66 (s, 3H), 3.59 (q, *J* = 6.1 Hz, 2H), 2.57 (t, *J* = 6.1 Hz, 2H); $^{13}$C NMR (101 MHz, $CDCl_3$): δ 172.7, 167.8, 155.7, 155.7, 129.6, 127.0, 115.9, 67.5, 51.8, 34.3, 33.6. HR-MS(ESI) calcd. for formula $C_{12}H_{14}ClNO_4Na[M+Na]^+$: 294.0509; Found: 294.0505.

**Compound 2b** 297

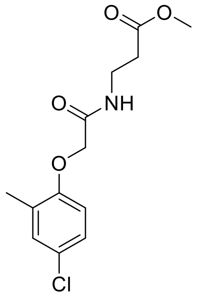


$^1$H NMR (400 MHz, $CDCl_3$): δ 7.10–7.19 (m, 3H), 6.65 (d, *J* = 8.6 Hz, 1H), 4.45 (s, 2H), 3.67 (s, 3H), 3.62 (q, *J* = 6.1 Hz, 2H), 2.58 (t, *J* = 5.8 Hz, 2H), 2.27 (s, 3H); $^{13}$C NMR (101 MHz, $CDCl_3$): δ 172.8, 167.9, 153.9, 130.8, 128.5, 126.7, 126.5, 112.3, 67.4, 51.8, 34.2, 33.6, 16.2. HR-MS(ESI) calcd. for formula $C_{13}H_{16}ClNO_4Na[M+Na]^+$: 308.0666; Found: 308.0663.

**Compound 3b** 298

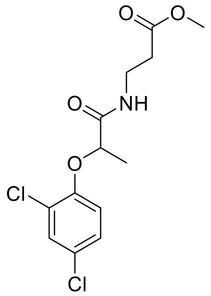


$^1$H NMR (400 MHz, $CDCl_3$): δ 7.39 (d, *J* = 2.5 Hz, 1H) 7.16 (dd, *J* = 2.5 and 8.8 Hz, 1H), 7.12 (br s, 1H), 6.80 (d, *J* = 8.8 Hz, 1H), 4.65 (q, *J* = 6.7 Hz, 1H), 3.62 (s, 3H), 3.50 - 3.60 (m, 2H), 2.53 (t, *J* = 5.8 Hz, 2H), 1.60 (d, *J* = 6.7 Hz, 3H); $^{13}$C NMR (101 MHz, $CDCl_3$): δ 172.4, 171.2, 151.4, 130.4, 127.9, 127.4, 124.6, 115.8, 76.5, 51.9, 34.7, 33.9, 18.5. HR-MS(ESI) calcd. for formula $C_{13}H_{15}Cl_2NO_4Na[M+Na]^+$: 342.0276; Found: 342.0270.

**Compound 4b** 299

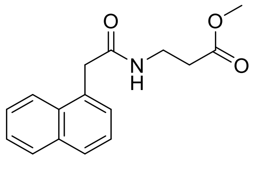


$^1$H NMR (400 MHz, $CDCl_3$): δ 7.82–7.93 (m, 3H), 7.38–7.54 (m, 4H), 5.81 (br s, 1H), 4.00 (s, 2H), 3.40 (s, 3H), 3.36–3.39 (m, 2H), 2.39 (t, *J* = 6.0 Hz, 2H); $^{13}$C NMR (101 MHz, $CDCl_3$): δ 172.3, 171.1, 151.2, 130.3, 127.8, 127.2, 124.5, 115.7, 51.8, 34.51, 33.7, 18.4. HR-MS(ESI) calcd. for formula $C_{16}H_{17}NO_3Na[M+Na]^+$: 294.1106; Found: 294.1111.

**Compound 1c** 300

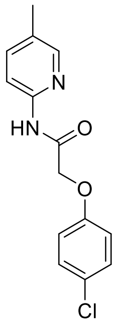


$^1$H NMR (400 MHz, $CDCl_3$): δ 8.79 (br s, 1H), 8.15 (d, *J* = 8.5 Hz, 1H), 8.13–8.14 (m, 1H), 7.55 (dd, *J* = 2.4 and 8.4 Hz, 1H), 7.29 (d, *J* = 6.8 Hz, 2H), 6.92 (d, *J* = 6.8 Hz, 2H), 4.59 (s, 2H), 2.31 (s, 3H); $^{13}$C NMR (101 MHz, $CDCl_3$): δ 166.1, 155.5, 148.1, 147.9, 139.0, 129.8 (2), 127.4 (2), 116.0, 113.7, 67.8, 17.8. HR-MS(ESI) calcd. for formula $C_{14}H_{13}ClN_2O_2Na[M+Na]^+$: 299.0563; Found: 299.0561.

301

**Compound 2c** 302

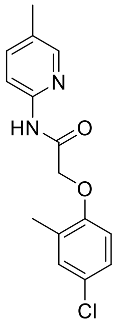


$^1$H NMR (400 MHz, $CDCl_3$): δ 8.80 (s, 1H), 8.08–8.20 (m, 2H), 7.56 (dd, *J* = 2.2 and 8.4 Hz, 1H), 7.18–7.19 (m, 1H), 7.14 (dd, *J* = 2.4 and 8.5 Hz, 1H), 6.75 (d, *J* = 8.7 Hz, 1H), 4.60 (s, 2H), 2.36 (s, 3H), 2.32 (s, 3H); $^{13}$C NMR (101 MHz, $CDCl_3$): δ 166.4, 153.9, 148.1, 147.9, 139.0, 131.0, 129.9, 128.8, 127.0, 126.8, 113.6, 112.8, 67.9, 17.9, 16.4. HR-MS(ESI) calcd. for formula $C_{15}H_{15}ClN_2O_2Na[M+Na]^+$: 313.0720; Found: 313.0725.

**Compound 3c** 303

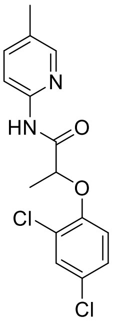


$^1$H NMR (400 MHz, $CDCl_3$): δ 9.01 (br s, 1H), 8.14 (m, 2H), 7.54 (dd, *J* = 2.1, and 8.1 Hz, 1H), 7.42 (d, *J* = 2.5 Hz, 1H), 7.19 (dd, *J* = 2.5 and 8.8 Hz, 1H), 6.87 – 6.92 (m, 1H), 4.77 (q, *J* = 6.7 Hz, 1H), 2.30 (s, 3H), 1.69 (d, *J* = 6.8 Hz, 3H); $^{13}$C NMR (101 MHz, $CDCl_3$): δ 169.6, 151.2, 148.3, 147.6, 139.1, 130.4, 129.8, 127.9, 127.8, 125.1, 116.6, 113.7, 76.9, 18.4, 17.8. HR-MS(ESI) calcd. for formula $C_{15}H_{14}Cl_2N_2O_2Na[M+Na]^+$: 347.0330; Found: 347.0331.

**Compound 1d** 304

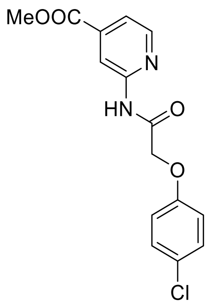


$^1$H NMR (400 MHz, $CDCl_3$): δ 9.06 (s, 1H), 8.88 (d, *J* = 16.9 Hz, 1H), 8.63–8.40 (m, 1H), 7.65–7.89 (m, 1H), 7.60–7.36 (m, 2H), 6.84–7.17 (m, 2H), 4.71 (s, 2H), 4.04 (s, 3H); $^{13}$C NMR (101 MHz, $CDCl_3$): δ 166.3, 165.2, 155.3, 151.1, 148.6, 139.8, 129.7, 127.4, 119.8, 115.9, 113.5, 67.5, 52.7. HR-MS(ESI) calcd. for formula $C_{15}H_{13}ClN_2O_4Na[M+Na]^+$: 343.0462; Found: 343.0466.

**Compound 2d** 305

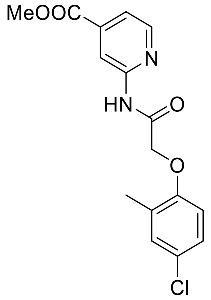


$^1$H NMR (400 MHz, $CDCl_3$): δ 8.96 (s, 1H), 8.79 (s, 1H), 8.45 (d, *J* = 5.1 Hz, 1H), 7.66 (dd, *J* = 1.3 and 5.1 Hz, 1H), 7.11–7.29 (m, 2H), 6.76 (d, *J* = 8.6 Hz, 1H), 4.64 (s, 2H), 3.97 (s, 3H), 2.37 (s, 3H); $^{13}$C NMR (101 MHz, $CDCl_3$): δ 166.6, 165.3, 153.7, 151.1, 148.8, 139.9, 131.1, 128.7, 127.1, 126.8, 119.9, 113.5, 112.8, 67.8, 52.8, 16.4. HR-MS(ESI) calcd. for formula $C_{11}H_{13}Cl_2NO_3Na[M+Na]^+$: 300.0170; Found: 300.0169. HR-MS(ESI) calcd. for formula $C_{16}H_{15}ClN_2O_4Na[M+Na]^+$: 357.0618; Found: 357.0612.

306

307

308

309

**Compound 3d** 310

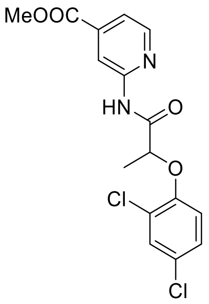

$^1$H NMR (400 MHz, $CDCl_3$): δ 9.15 (br s, 1H), 8.76 (s, 1H), 8.44 (d, *J* = 5.1 Hz, 1H), 7.54–7.76 (m, 1H), 7.34–7.51 (m, 1H), 7.20 (ddd, *J* = 1.1, 2.5 and 8.8 Hz, 1H), 6.92 (d, *J* = 8.8 Hz, 1H), 4.82 (q, *J* = 6.8 Hz, 1H), 3.95 (s, 3H), 1.71 (d, *J* = 6.7 Hz, 4H); $^{13}$C NMR (101 MHz, $CDCl_3$): δ 169.8, 165.3, 151.40, 151.1, 148.9, 139.8, 130.5, 128.1, 128.0, 125.1, 119.7, 116.6, 113.5, 76.9, 52.7, 18. HR-MS(ESI) calcd. for formula $C_{16}H_{15}Cl_2N_2O_4[M+H]^+$: 369.0409; Found: 369.0402.

**Compound 1e** 311

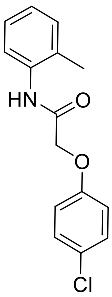

$^1$H NMR (400 MHz, $CDCl_3$): δ 8.17 (br s, 1H), 7.96 (d, *J* = 8.0 Hz, 1H), 7.30 (d, *J* = 8.9 Hz, 2H), 7.19 - 7.26 (m, 2H), 7.10 (t, *J* = 7.3 Hz, 1H), 6.92 (d, *J* = 8.9 Hz, 2H), 4.63 (s, 2H), 2.23 (s, 3H); $^{13}$C NMR (101 MHz, $CDCl_3$): δ 165.7, 155.5, 134.6, 130.6, 129.9, 128.6, 127.5, 127.0, 125.5, 122.4, 116.0, 67.9, 17.4. HR-MS(ESI) calcd. for formula $C_{15}H_{14}ClNO_2Na[M+Na]^+$: 298.0611; Found: 298.0608.

**Compound 2e** 312

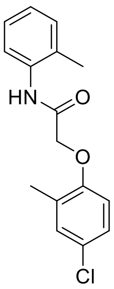

$^1$H NMR (400 MHz, $CDCl_3$): δ 8.24 (br s, 1H), 8.05 (d, *J* = 7.36 Hz, 1H), 7.15 - 7.27 (m, 4H), 7.10 (dt, *J* = 1.1 and 7.3 Hz, 1H), 6.77 (d, *J* = 8.6 Hz, 1H), 4.63 (s, 2H), 2.33 (s, 3H), 2.27 (s, 3H); $^{13}$C NMR (101 MHz, $CDCl_3$): δ 165.7, 153.7, 134.8, 131.0, 130.5, 128.1, 127.8, 127.0 (2), 125.3, 121.9, 112.5, 67.7, 17.5, 16.4. HR-MS(ESI) calcd. for formula $C_{16}H_{16}ClNO_2Na[M+Na]^+$: 312.0767; Found: 312.0763.

**Compound 3e** 313

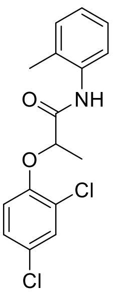

$^1$H NMR (400 MHz, $CDCl_3$): δ 8.36 (br s, 1H), 7.98 (d, *J* = 7.6 Hz, 1H), 7.44 (d, J = 2.6 Hz, 1H), 7.18 - 7.25 (m, 3H), 7.09 (dt, *J* = 1.1 and 7.5 Hz, 1H), 6.92 (d, J = 8.8 Hz, 1H), 4.90 (q, *J* = 6.7 Hz, 1H), 2.24 (s, 3H), 1.72 (d, *J* = 6.7Hz, 3H); $^{13}$C NMR (101 MHz, $CDCl_3$): δ 169.0, 151.0, 135.0, 130.7, 130.6, 128.7, 128.2, 127.7, 127.0, 125.4, 124.6, 122.3, 115.8, 76.4, 18.5, 17.7. HR-MS(ESI) calcd. for formula $C_{16}H_{15}Cl_2NO_2Na[M+Na]^+$: 346.0378; Found: 346.0376.

**Compound 4e** 314

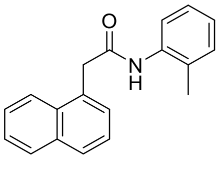

$^1$H NMR (400 MHz, $CDCl_3$): δ 8.05 (d, *J* = 7.8 Hz, 1H), 7.89 - 7.93 (m, 2H), 7.82 (d, *J* = 8.0, Hz, 1H), 7.49 – 7.59 (m, 4H), 7.12 – 7.17 (m, 1H), 6.97 (m, 2H), 6.82 (br s, 1H), 4.24 (s, 2H), 1.51 (s, 3H); $^{13}$C NMR (101 MHz, $CDCl_3$): δ 168.9, 135.4, 135.1, 134.0, 132.0, 130.9, 130.2, 128.9, 128.6, 128.2, 127.2, 126.7, 126.5, 125.7, 124.9, 123.8, 122.0, 42.8, 16.7. HR-MS(ESI) calcd.

for formula $C_{19}H_{17}Cl_2NONa[M+Na]^+$: 298.1208; Found: 298.1201.

**Compound 1f** 315

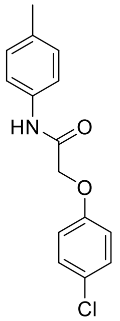


$^{1}H$ NMR (400 MHz, $CDCl_3$): δ 8.18 (s, 1H), 7.44 – 7.55 (m, 2H), 7.26 – 7.38 (m, 2H), 7.17 (d, $J$ = 8.2 Hz, 2H), 6.91 – 7.01 (m, 2H), 4.58 (s, 2H), 2.34 (s, 3H); $^{13}C$ NMR (101 MHz, $CDCl_3$): δ 165.6, 155.6, 134.7, 134.1, 129.8, 129.6, 127.4, 120.2, 116.1, 67.8, 20.9. HR-MS(ESI) calcd. for formula $C_{15}H_{14}ClNO_2Na[M+Na]^+$: 298.0611; Found: 298.0613.

**Compound 2f** 316

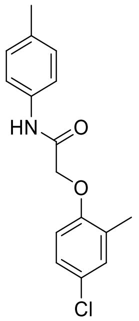


$^{1}H$ NMR (400 MHz, $CDCl_3$): δ 8.21 (br s, 1H), 7.46 (d, $J$ = 8.4 Hz, 2H), 7.18 (m, 4H), 6.77 (d, $J$ = 8.6 Hz, 1H), 4.58 (s, 2H), 2.35 (s, 6H); $^{13}C$ NMR (101 MHz, $CDCl_3$): δ 165.8, 153.8, 134.7, 134.1, 131.0, 129.6, 128.4, 127.0, 120.0, 113.0, 68.0, 20.9, 16.3. HR-MS(ESI) calcd. for formula $C_{16}H_{16}ClNO_2Na[M+Na]^+$: 312.0767; Found: 312.0770.

**Compound 3f** 317

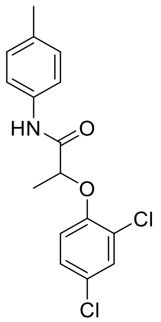


$^{1}H$ NMR (400 MHz, $CDCl_3$): δ 8.51 (br s, 1H), 7.39 – 7.53 (m, 3H), 7.23 (dd, $J$ = 2.5 and 8.8 Hz, 1H), 7.16 (d, $J$ = 8.2 Hz, 2H), 6.94 (d, $J$ = 8.8 Hz, 1H). 4.80 (q, $J$ = 6.7 Hz, 1H), 2.34 (s, 3H), 1.72 (d, $J$ = 6.7 Hz, 3H); $^{13}C$ NMR (101 MHz, $CDCl_3$): δ 168.8, 151.2, 134.5, 134.4, 130.3, 129.6, 128.1, 127.7, 124.5, 119.8, 116.2, 77.0, 20.9, 18.4. HR-MS(ESI) calcd. for formula $C_{16}H_{15}Cl_2NO_2Na[M+Na]^+$: 346.0378; Found: 346.0379.

**Compound 4f** 318

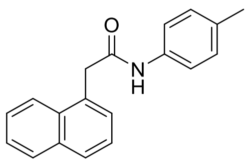


$^{1}H$ NMR (400 MHz, $CDCl_3$): δ 8.04 (dd, $J$ = 2.8 and 6.0 Hz, 1H), 7.87-7.93 (m, 2H), 7.48 – 7.60 (m, 4H), 7.20 (d, $J$ = 8.4 Hz, 2H), 7.04 (d, $J$ = 8.2 Hz, 2H), 4.17 (s, 2H), 2.27 (s, 3H); $^{13}C$ NMR (101 MHz, $CDCl_3$): δ 169.1, 134.9, 134.1, 134.0, 132.1, 130.8, 129.3, 128.9, 128.8, 128.5, 127.0, 126.3, 125.7, 123.7, 120.1, 42.7, 20.8. HR-MS(ESI) calcd. for formula $C_{19}H_{18}NO[M+H]^+$: 276.1388; Found: 276.1386.

319

320

321

322

**Compound 1g** 323

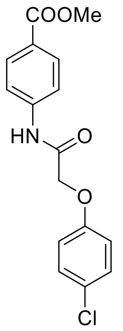

$^{1}$H NMR (400 MHz, $CDCl_3$): δ 8.35 (br s, 1H), 8.03 (d, *J* = 8.8 Hz, 2H), 7.67 (d, *J* = 8.8 Hz, 2H), 7.30 (d, *J* = 9.0 Hz, 2H), 6.93 (d, *J* = 9.0 Hz, 2H), 4.60 (s, 2H), 3.91 (s, 3H); $^{13}$C NMR (101 MHz, $CDCl_3$): δ 166.6, 166.1, 155.5, 140.9, 131.1, 130.1, 127.9, 126.6, 119.3, 116.3, 68.0, 52.2. HR-MS(ESI) calcd. for formula $C_{16}H_{14}ClNO_4Na[M+Na]^+$: 342.0509; Found: 342.0503.

**Compound 2g** 324

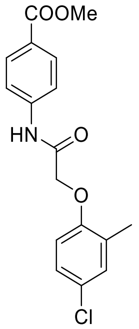

$^{1}$H NMR (400 MHz, $CDCl_3$): δ 8.43 (br s, 1H), 8.01 – 8.11 (m, 2H), 7.65 – 7.73 (m, 2H), 7.12 – 7.25 (m, 2H), 6.77 (d, *J* = 8.6 Hz, 1H), 4.60 (s, 2H), 3.92 (s, 3H), 2.36 (s, 3H); $^{13}$C NMR (101 MHz, $CDCl_3$): δ 166.4, 166.2, 153.7, 140.8, 131.1, 130.9, 128.4, 127.3, 127.0, 126.3, 119.0, 113.1, 68.0, 52.1, 16.3. HR-MS(ESI) calcd. for formula $C_{17}H_{16}ClNO_4Na[M+Na]^+$: 356.0666; Found: 356.0663.

**Compound 3g** 325

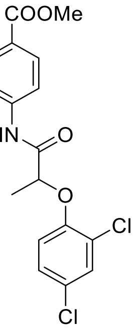

$^{1}$H NMR (400 MHz, $CDCl_3$): δ 8.77 (br s, 1H), 8.02 (d, *J* = 8.7 Hz, 2H), 7.67 (d, *J* = 8.7 Hz, 2H), 7.45 (d, *J* = 2.5 Hz, 1H), 7.23 (dd, *J* = 2.5 and 8.7 Hz, 1H), 6.92 (d, *J* = 8.8 Hz, 1H), 4.81 (q, *J* = 6.7 Hz, 1H), 3.91 (s, 3H), 1.71 (d, *J* = 6.7 Hz, 3H); $^{13}$C NMR (101 MHz, $CDCl_3$): δ 169.2, 166.5, 150.9, 141.1, 130.9, 130.4, 128.2, 128.0, 126.2, 124.61, 119.0, 116.4, 52.1, 30.9, 18.3. HR-MS(ESI) calcd. for formula $C_{17}H_{15}Cl_2NO_4Na[M+Na]^+$: 390.0276; Found: 390.0274.

**Compound 4g** 326

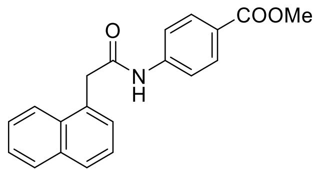

$^{1}$H NMR (400 MHz, $CDCl_3$): δ 7.98 - 8.09 (m, 1H), 7.88 - 8.04 (m, 4H), 7.48 – 7.61 (m, 4H), 7.40 (d, *J* = 8.7 Hz, 2H), 7.25 (br s, 1H), 4.21 (s, 2H), 3.87 (s, 3H); $^{13}$C NMR (101 MHz, $CDCl_3$): δ 169.2, 166.5, 141.6, 134.1, 132.0, 130.6, 130.2, 129.1, 129.0, 128.6, 127.2, 126.5, 125.7, 123.5, 118.8, 52.0, 43.0. HR-MS(ESI) calcd. for formula $C_{20}H_{17}NO_3Na[M+Na]^+$: 342.1106; Found: 342.1110.

**Compound 1h** 327

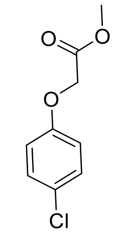

$^{1}$H NMR (400 MHz, DMSO-$d_6$): δ 7.32 (d, *J* = 9.0 Hz, 2H), 6.96 (d, *J* = 9.1 Hz, 2H), 4.81 (s, 2H), 3.69 (s, 3H); $^{13}$C NMR (101 MHz, DMSO- $d_6$): δ 169.5, 156.9, 129.6 (2), 125.4, 116.7 (2), 65.2, 52.3. HR-MS(ESI) calcd. for formula $C_9H_9ClO_3[M]^+$: 200.0240; Found: 200.0247.

**Compound 2h** 328

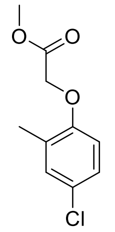

$^{1}$H NMR (400 MHz, $CDCl_3$): δ 7.13 (d, *J* = 2.1 Hz, 1H), 7.06 (dd, *J* = 2.4 and 8.6 Hz, 1H), 6.60 (d, *J* = 8.6 Hz, 1H), 4.63 (s, 2H), 3.80 (s, 3H), 2.26 (s, 3H); $^{13}$C NMR (101 MHz, $CDCl_3$): δ 169.4, 154.8, 131.0, 129.4, 126.5, 112.4, 65.9, 52.4, 16.3. HR-MS(ESI) calcd. for formula $C_{10}H_{11}ClO_3[M]^+$: 214.0397; Found: 214.0404.

**Compound 3h** 329

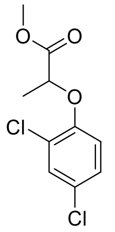

$^{1}$H NMR (400 MHz, $CDCl_3$): δ 7.37 (d, *J* = 2.5 Hz, 1H), 7.13 (dd, *J* = 2.5 and 8.8 Hz, 1H), 6.76 (d, *J* = 8.8 Hz, 1H), 4.72 (q, *J* = 6.8 Hz, 1H), 3.75 (s, 3H), 1.66 (d, *J* = 6.8 Hz, 3H); $^{13}$C NMR (101 MHz, $CDCl_3$): δ 171.9, 152.4, 130.4, 127.7, 127.3, 124.9, 116.2, 74.5, 52.6, 18.6. HR-MS(ESI) calcd. for formula $C_{10}H_{10}Cl_2O_3[M]^+$: 248.0007; Found: 248.0012.

**Compound 4h** 330

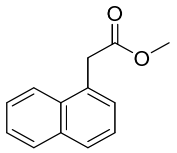

$^{1}$H NMR (400 MHz, $CDCl_3$): δ 8.00 (d, *J* = 8.2 Hz, 1H), 7.87 (d, *J* = 7.7 Hz, 1H), 7.80 (d, *J* = 7.2 Hz, 1H), 7.42 – 7.57 (m, 4H), 4.09 (s, 2H), 3.69 (s, 3H); $^{13}$C NMR (101 MHz, $CDCl_3$): δ 172.1, 133.8, 132.1, 130.5, 128.7, 128.1, 128.0, 126.4, 125.8, 125.5, 123.8, 52.2, 39.0. HR-MS(ESI) calcd. for formula $C_{13}H_{12}O_2[M]^+$: 200.0837; Found: 200.0836.

**Compound 1i** 331

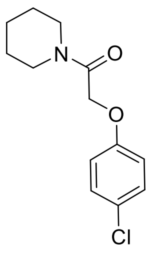

$^{1}$H NMR (400 MHz, $CDCl_3$): 7.257 (d, *J* = 9.0 Hz, 2H), 6.88 (d, *J* = 9.0 Hz, 2H), 4.66 (s, 2H), 3.55 (t, *J* = 7.1 Hz, 2H), 3.47 (t, *J* = 7.1 Hz, 2H), 1.54–1.65 (m, 6H); $^{13}$C NMR (101 MHz, $CDCl_3$): δ 165.9, 129.6, 127.0, 116.1, 68.1, 46.5, 43.4, 26.6, 25.7, 24.6. HR-MS(ESI) calcd. for formula $C_{13}H_{16}ClNO_2Na[M+Na]^+$: 276.0767; Found: 276.0765.

332

**Compound 1j** 333

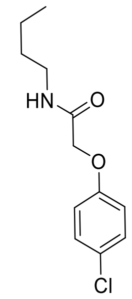

$^{1}H$ NMR (400 MHz, $CDCl_3$): 7.27 (d, *J* = 9.04 Hz, 2H), 6.85 (d, *J* = 9.04 Hz, 2H), 4.46 (s, 2H), 3.34 (q, *J* = 7.1 Hz, 2H), 1.46–1.59 (m, 2H), 1.22–1.45 (m, 2H), 0.93 (t, *J* = 7.3 Hz, 3H); $^{13}C$ NMR (101 MHz, $CDCl_3$): δ 167.6, 155.8, 129.7, 127.1, 115.9, 67.60 38.8, 31.5, 20.0, 13.7. HR-MS(ESI) calcd. for formula $C_{12}H_{16}ClNO_2Na[M+Na]^+$: 264.0767; Found: 264.0764.

**Compound 1k**

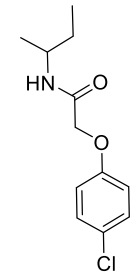

$^{1}H$ NMR (400 MHz, $CDCl_3$): δ 7.23–7.33 (m, 2H), 6.79–6.93 (m, 2H), 6.28 (s, 1H), 4.45 (s, 2H), 4.01 (ddd, *J* = 6.7, 8.6 and 13.3 Hz, 1H), 1.45–1.55 (m, 2H), 1.17 (d, *J* = 6.6 Hz, 3H), 0.90 (t, *J* = 7.4 Hz, 3H); $^{13}C$ NMR (101 MHz, $CDCl_3$): δ 167.0, 155.8, 129.7, 127.1, 116.0, 67.6, 46.4, 29.6, 20.4, 10.3. HR-MS(ESI) calcd. for formula $C_{12}H_{16}ClNO_2Na[M+Na]^+$: 264.0767; Found: 264.0770.

**Compound 1l**

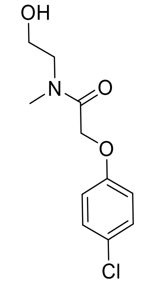

$^{1}H$ NMR (400 MHz, $CDCl_3$): 7.12–7.41 (m, 2H), 6.86 (dd, *J* = 9.0 and 11.2 Hz, 2H), 4.79 (s, 1H), 4.69 (s, 1H), 3.77 (s, 3H), 3.55 (t, *J* = 5.3 Hz, 2H), 3.46 (d, *J* = 7.3 Hz, 2H); $^{13}C$ NMR (101 MHz, $CDCl_3$): δ 169.0, 168.5, 156.5, 156.3, 129.3, 129.2, 126.5, 126.2, 115.9, 67.1, 66.7, 60.7, 59.1, 51.2, 35.6, 33.3. HR-MS(ESI) calcd. for formula $C_{11}H_{14}ClNO_3Na[M+Na]^+$: 266.0560; Found: 266.0557.

**Compound 1m**

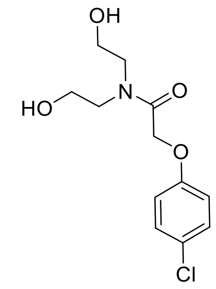

$^{1}H$ NMR (400 MHz, DMSO-$d_6$) δ 7.27 (d, *J* = 9.0 Hz, 2H), 7.01 (s, 1H), 6.90 (d, *J* = 9.0 Hz, 2H), 3.33-3.57 (m, 8H); $^{13}C$ NMR (101 MHz, DMSO-$d_6$): δ 167.8, 157.6, 135.6, 129.4, 124.6, 116.7, 66.1, 59.1, 59.0, 49.7, 48.2. HR-MS(ESI) calcd. for formula $C_{12}H_{16}ClNO_4Na[M+Na]^+$: 296.0666; Found: 296.0668.

**Compound 1n**

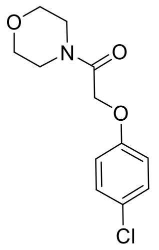


[1]H NMR (400 MHz, $CDCl_3$): 7.24 (d, *J* = 9.04 Hz, 2H), 6.88 (d, *J* = 9.0 Hz, 2H), 4.68 (s, 2H), 3.55–3.70 (m, 8H); [13]C NMR (101 MHz, $CDCl_3$): δ 166.2, 156.3, 129.6, 126.7, 115.9, 67.8, 66.8, 66.7, 45.8, 42.4. HR-MS(ESI) calcd. for formula $C_{12}H_{14}ClNO_3Na[M+Na]^+$: 278.0560; Found: 278.0562.

**Compound 1o**

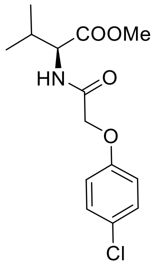


[1]H NMR (400 MHz, $CDCl_3$): 7.27 (d, *J* = 8.8 Hz, 2H), 6.98 (d, *J* = 8.5 Hz, 1H), 6.88 (d, *J* = 8.8 Hz, 2H), 4.62 (q, *J* = 5.0 Hz, 1H), 4.51 (d, *J* = 3.1 Hz, 2H), 3.75 (s, 3H), 2.29–2.11 (m, 1H), 0.93 (d, *J* = 6.9 Hz, 3H), 0.89 (d, *J* = 6.9 Hz, 3H); [13]C NMR (101 MHz, $CDCl_3$): δ 171.9, 167.8, 155.7, 129.7, 127.2, 116.1, 67.6, 56.6, 52.3, 31.3, 18.9, 17.7. HR-MS(ESI) calcd. for formula $C_{14}H_{18}ClNO_4Na[M+Na]^+$: 322.0822; Found: 322.0819.

**Compound 1p**

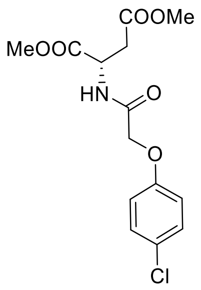


[1]H NMR (400 MHz, $CDCl_3$): δ 7.53 (d, *J* = 8.4 Hz, 1H), 7.32 – 7.22 (m, 2H), 6.83 – 6.93 (m, 2H), 4.87 – 4.99 (m, 1H), 4.44 – 4.55 (m, 2H), 3.76 (s, 3H), 3.66 (s, 3H), 3.07 (dd, *J* = 4.4 and 17.3 Hz, 1H), 2.86 (dd, *J* = 4.5 and 17.3 Hz, 1H); [13]C NMR (101 MHz, $CDCl_3$): δ 171.2, 170.6, 167.8, 155.7, 129.6, 127.1, 116.1, 67.4, 53.0, 52.1, 47.8, 35.9. HR-MS(ESI) calcd. for formula $C_{14}H_{16}ClNO_6Na[M+Na]^+$: 352.0564; Found: 352.0563.

**Compound 1q**

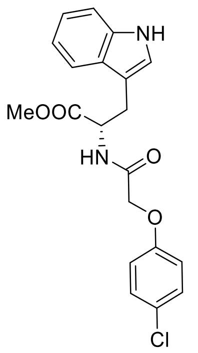


[1]H NMR (400 MHz, $CDCl_3$): δ 8.10 (br s, 1H), 7.54 (dd, *J* = 7.9 Hz, 1H), 7.37 (d, *J* = 7.9 Hz, 1H), 7.18–7.23 (m, 3H), 7.01 – 7.15 (m, 2H), 6.90 (d, *J* = 7.8 Hz, 1H), 6.65–6.67 (m, 2H), 4.97–5.02 (m, 1H), 4.43 (q, *J* = 7.2 Hz, 2H), 3.73 (s, 3H), 3.36 (d, *J* = 5.5 Hz, 2H); [13]C NMR (101 MHz, $CDCl_3$): δ 171.8, 167.7, 155.6, 136.1, 129.5, 127.4, 126.7, 122.7, 122.3, 119.8, 118.4, 116.0, 111.3, 109.6, 67.4, 52.5, 27.4. HR-MS(ESI) calcd. for formula $C_{20}H_{19}ClN_2O_4Na[M+Na]^+$: 409.0931; Found: 409.0932.

**Compound 1r**

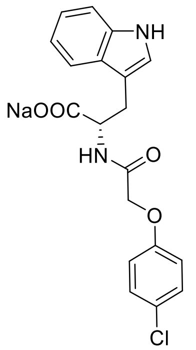

[1]H NMR (400 MHz, $D_2O$): δ 7.31 (d, *J* = 7.9 Hz, 1H), 7.19 (d, *J* = 9.0 Hz, 1H), 7.10 (d, *J* = 8.1 Hz, 1H), 6.94–6.71 (m, 5H), 6.16 (d, *J* = 8.9 Hz, 2H), 4.41 (dd, *J* = 7.6, 4.8 Hz, 1H), 4.10 (d, *J* = 15.4 Hz, 1H), 3.88 (d, *J* = 15.4 Hz, 1H), 3.13 (dd, *J* = 14.8, 4.8 Hz, 1H), 3.00 (dd, *J* = 14.9, 7.7 Hz, 1H); $^{13}$C NMR (101 MHz, $D_2O$): δ 178.0, 169.9, 155.2, 135.9, 129.3, 129.1, 127.1, 126.1, 123.8, 121.6, 119.1, 118.1, 115.8, 115.7, 111.6, 109.5, 66.5, 55.4, 27.0. HR-MS(ESI) calcd. for formula $C_{20}H_{19}ClN_2O_4Na[M+H]^+$: 395.0769; Found: 395.0770.

**Compound 1s**

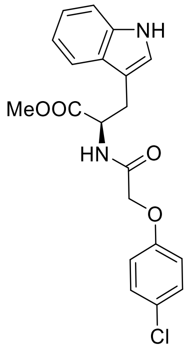

[1]H NMR (400 MHz, $CDCl_3$): δ 8.25 (br s, 1H), 7.54 (d, *J* = 0.7 and 7.9 Hz, 1H), 7.37 (d, *J* = 8.1 Hz, 1H), 7.16–7.26 (m, 3H), 7.01 – 7.15 (m, 2H), 6.90 (d, *J* = 2.3 Hz, 1H), 6.61–6.71 (m, 2H), 4.99 (dt, *J* = 5.5 and 8.0 Hz, 1H), 4.34 – 4.54 (m, 2H), 3.75 (s, 3H), 3.36 (d, *J* = 5.5 Hz, 2H); $^{13}$C NMR (101 MHz, $CDCl_3$): δ 171.9, 167.8, 155.6, 136.1, 129.5, 127.4, 126.7, 122.8, 122.3, 119.8, 118.4, 116.0, 111.4, 109.5, 67.4, 52.5, 27.4. HR-MS(ESI) calcd. for formula $C_{20}H_{19}ClN_2O_4Na[M+Na]^+$: 409.0931; Found: 409.0934.

**Compound 1t**

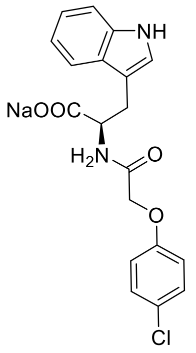

[1]H NMR (400 MHz, $D_2O$): δ 7.52 (d, *J* = 8.0 Hz, 1H), 7.37 (d, *J* = 8.0, 1H), 7.10 (t, *J* = 4.2 Hz, 1H), 7.06-6.93 (m, 4H), 6.46 (d, *J* = 9.0 Hz, 2H), 4.52-4.47 (m, 1H), 4.38 (d, *J* = 15.7 Hz, 1H), 4.21 (d, *J* = 15.7 Hz, 1H), 3.30 (dd $J_1$ = 14.8 Hz, $J_2$ = 4.6 Hz, 1H), 3.04 (dd, $J_1$ = 14.8 Hz, $J_2$ = 8.6 Hz, 1H). $^{13}$C NMR (101 MHz, $D_2O$): δ 178.0, 170.0, 155.3, 135.9, 129.3, 129.1 126.9, 126.0, 123.9, 121.5, 119.0, 118.2, 115.7, 115.6, 111.6, 109.5, 66.6, 55.1, 27.0. HR-MS(ESI) calcd. for formula $C_{20}H_{19}ClN_2O_4Na[M+H]^+$: 395.0769; Found: 395.0774.

### $^1$H- and $^{13}$C-NMR spectra

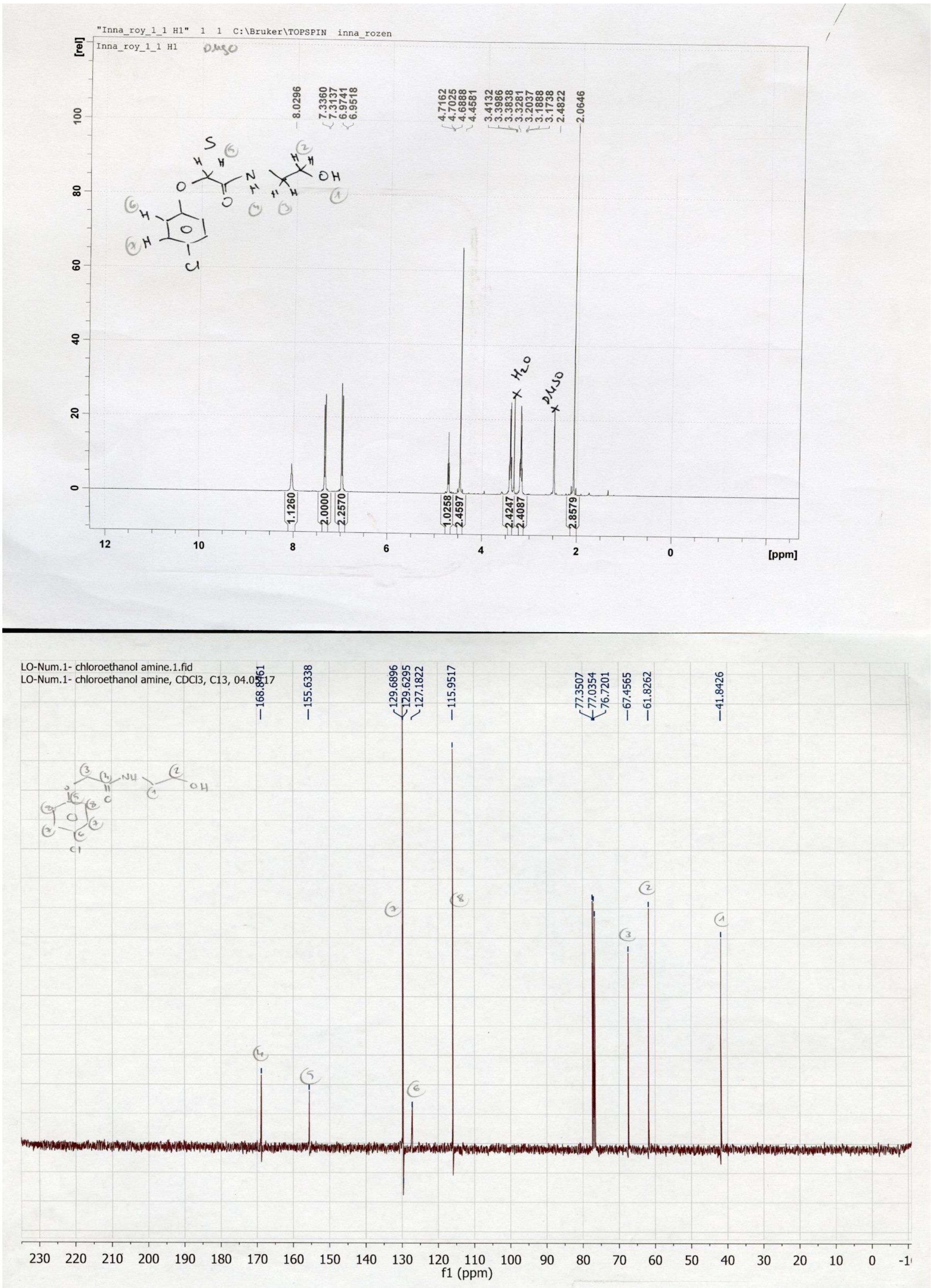


**Figure 27.** $^1$H (top) and $^{13}$C (bottom) NMR spectra of compound **1a**.

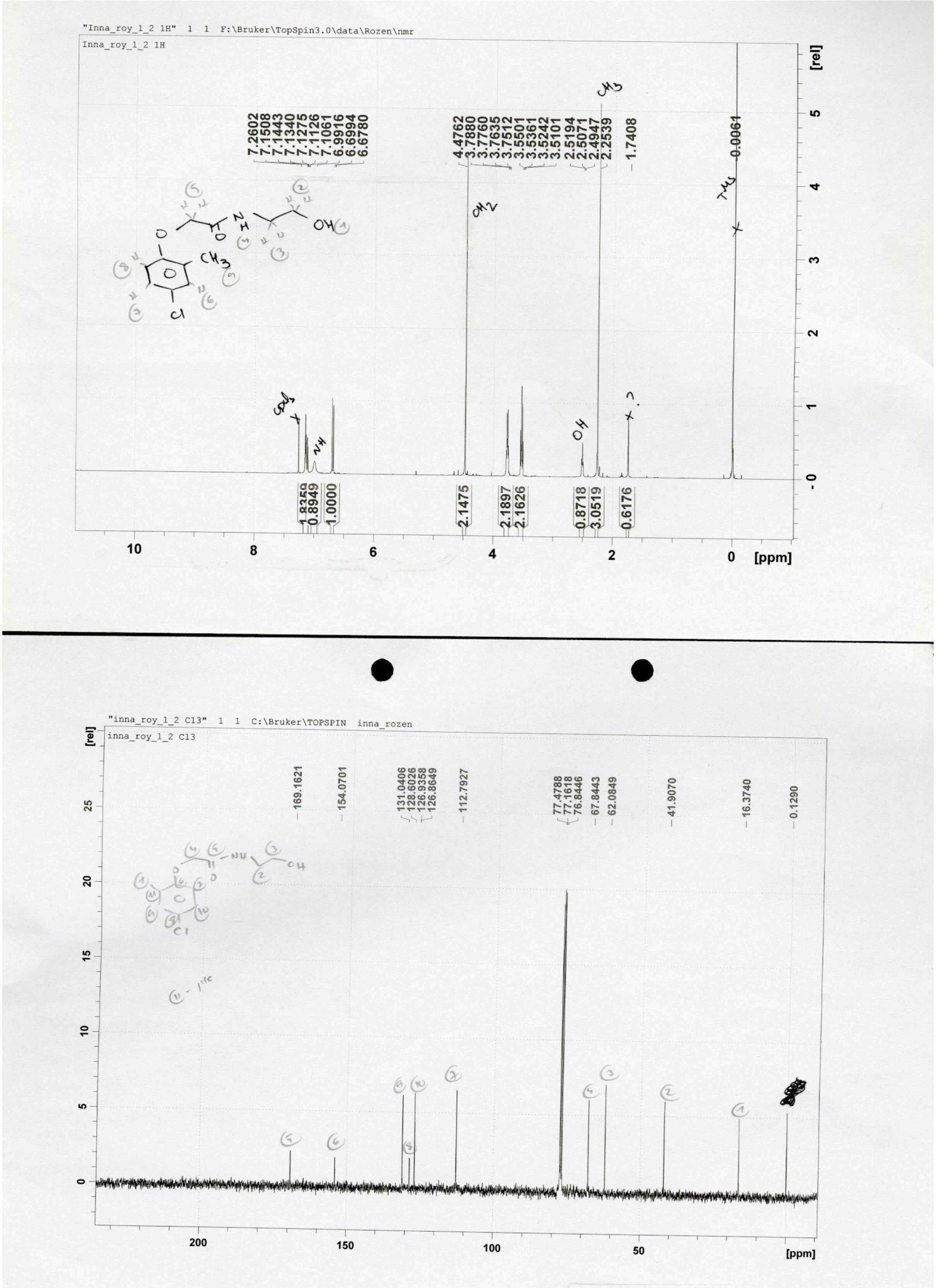


**Figure 28.** $^{1}H$ (top) and $^{13}C$ (bottom) NMR spectra of compound **2a**.

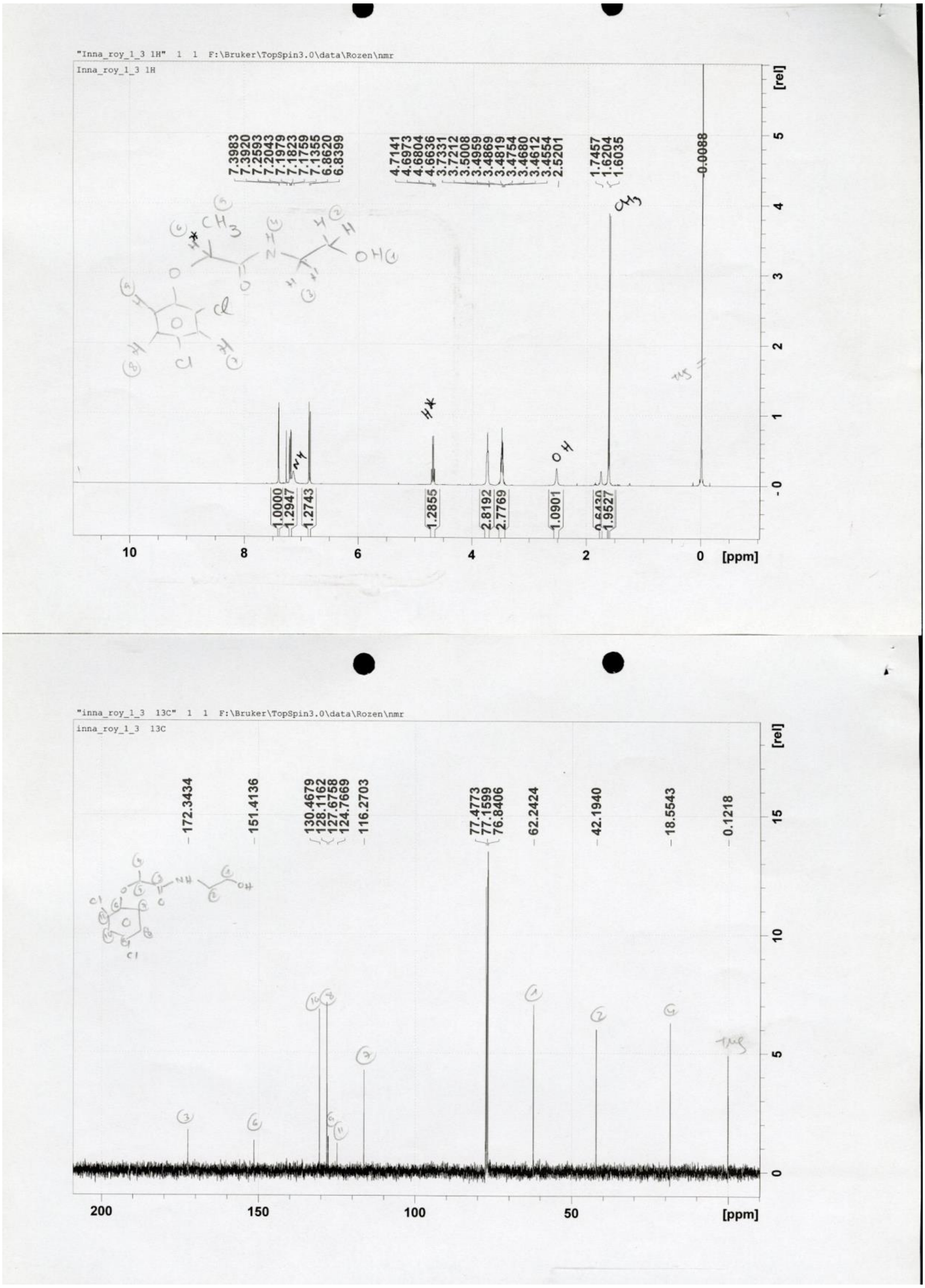

**Figure 29.** $^{1}H$ (top) and $^{13}C$ (bottom) NMR spectra of compound **3a**.

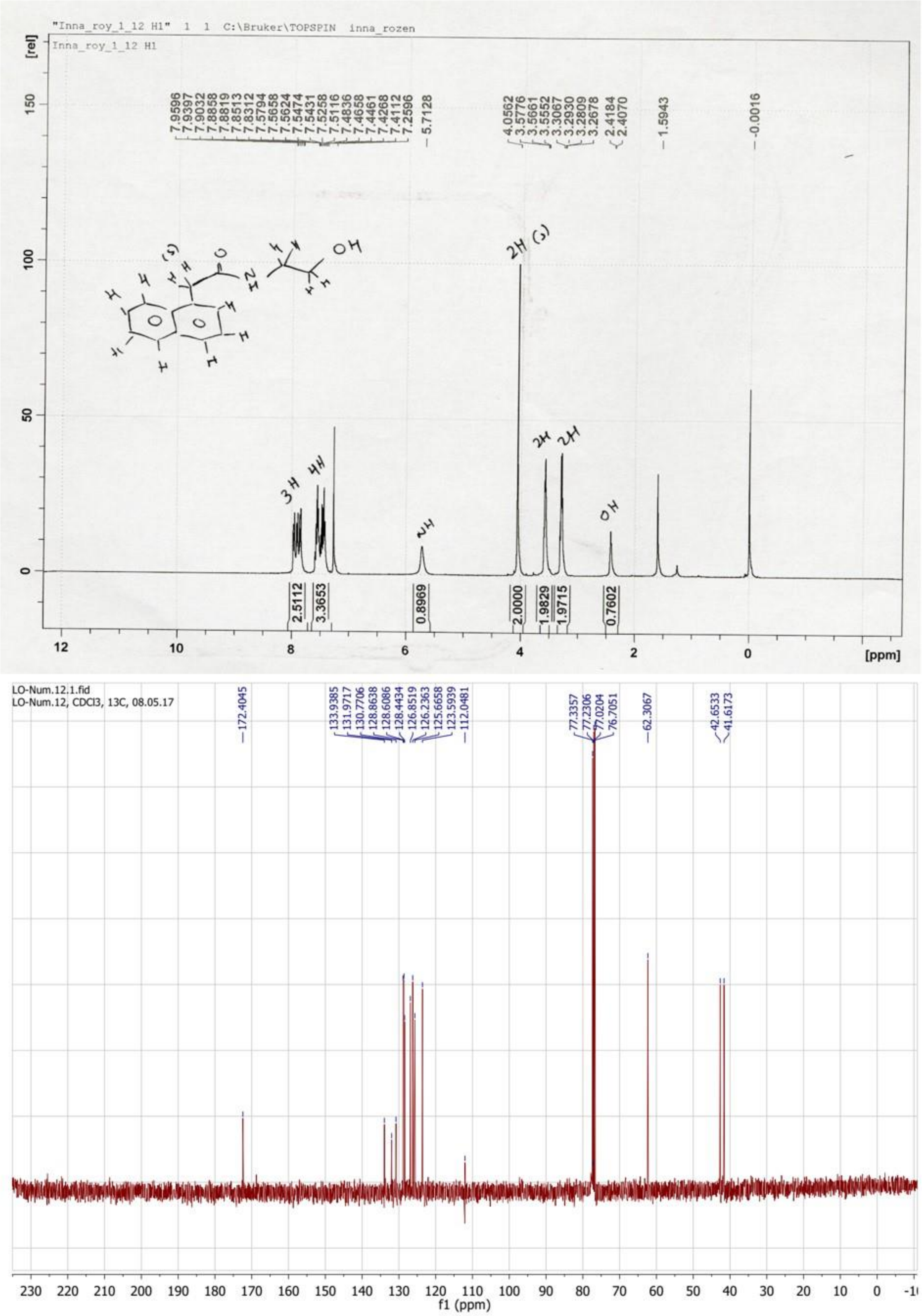


**Figure 30.** $^{1}H$ (top) and $^{13}C$ (bottom) NMR spectra of compound **4a**.

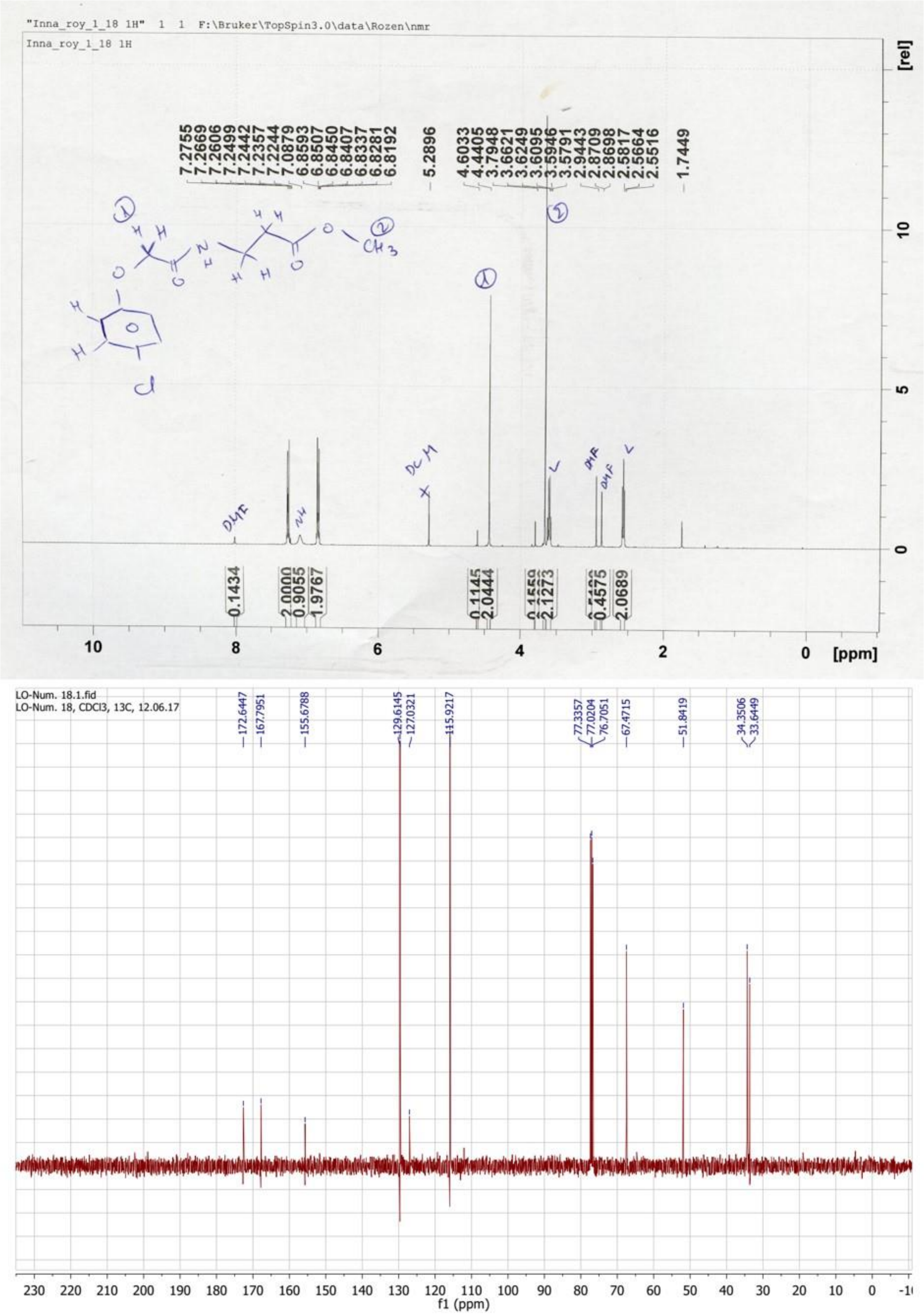


**Figure 31.** $^{1}H$ (top) and $^{13}C$ (bottom) NMR spectra of compound **1b**.

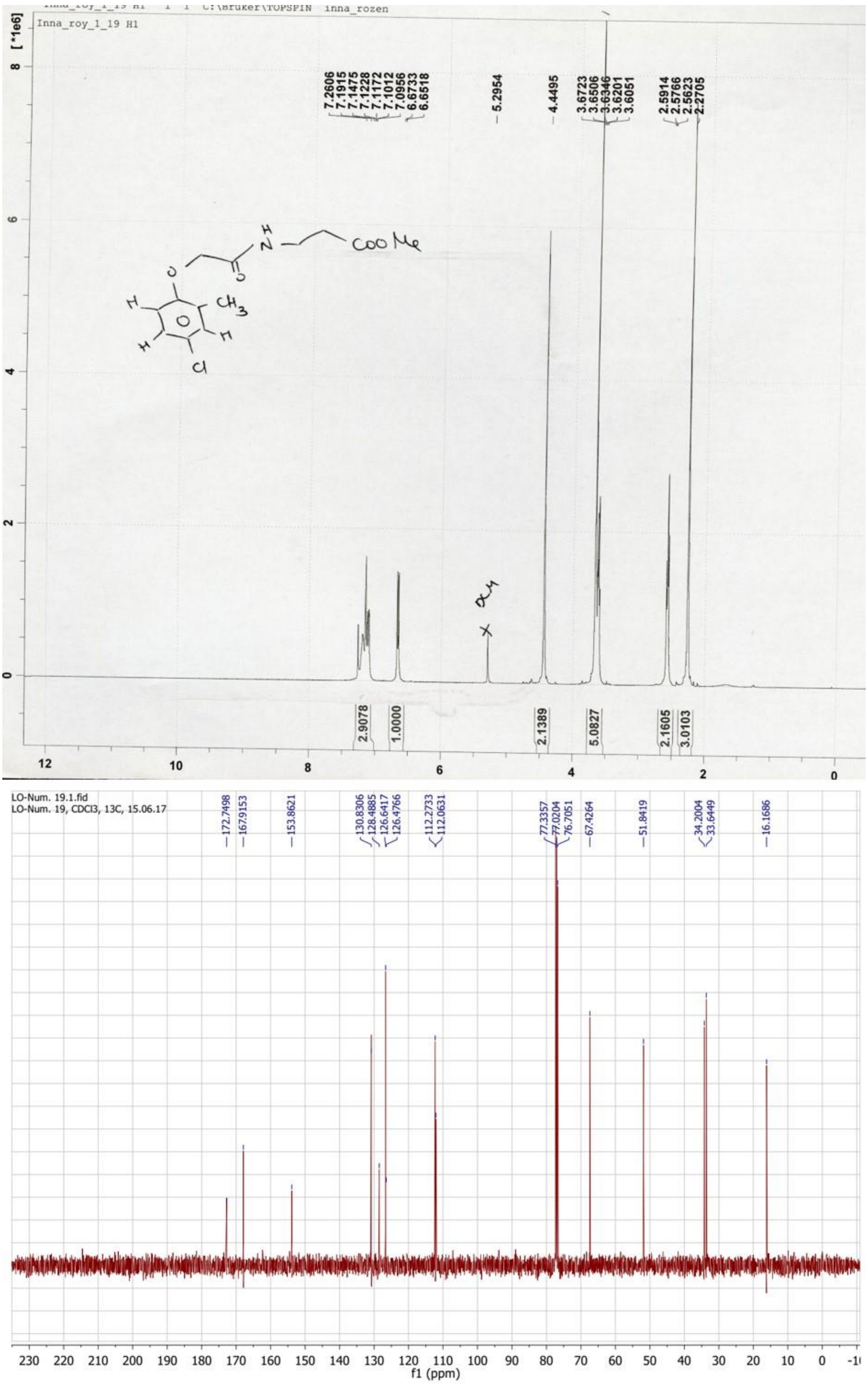


**Figure 32.** $^{1}H$ (top) and $^{13}C$ (bottom) NMR spectra of compound **2b**.

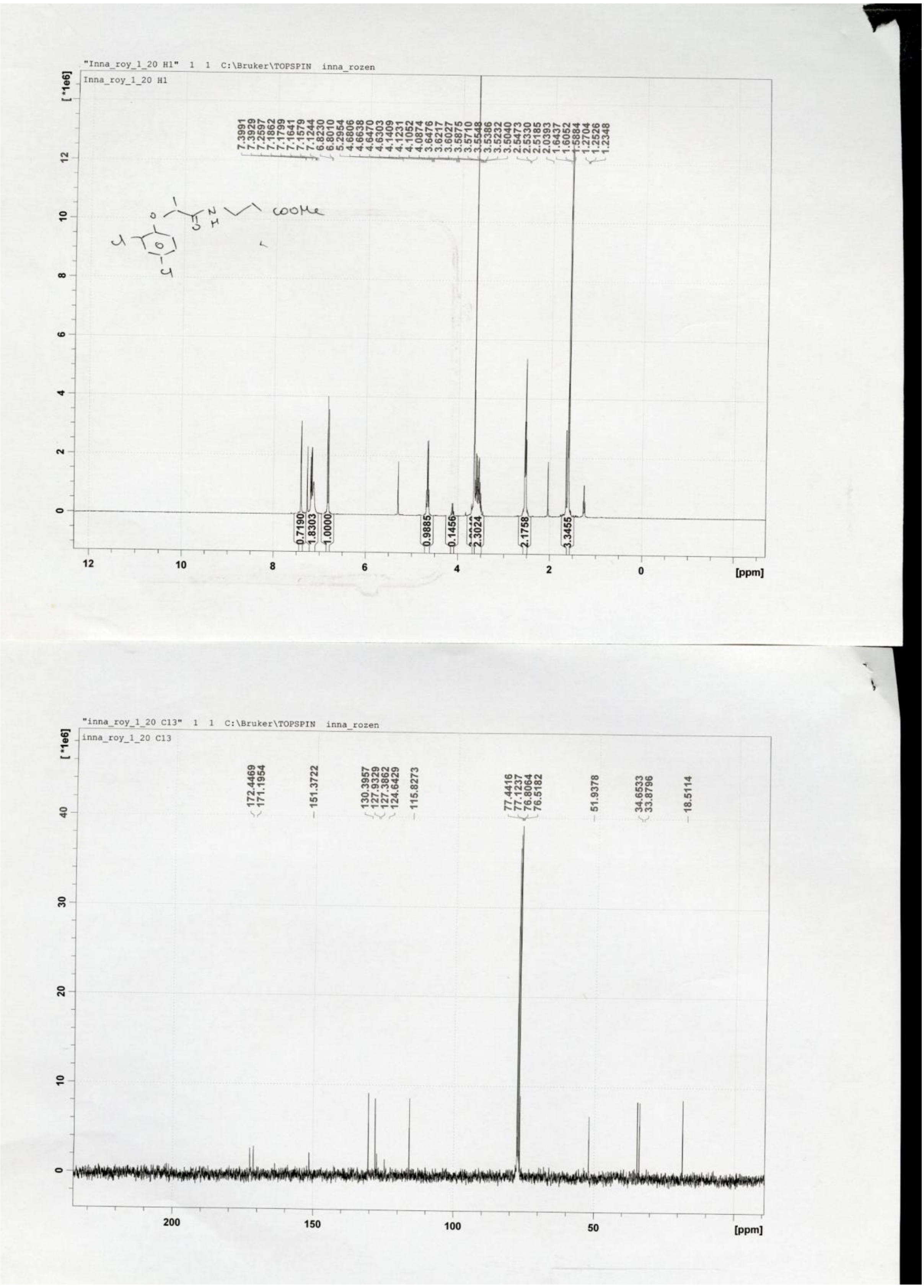


**Figure 33.** $^{1}H$ (top) and $^{13}C$ (bottom) NMR spectra of compound **3b**.

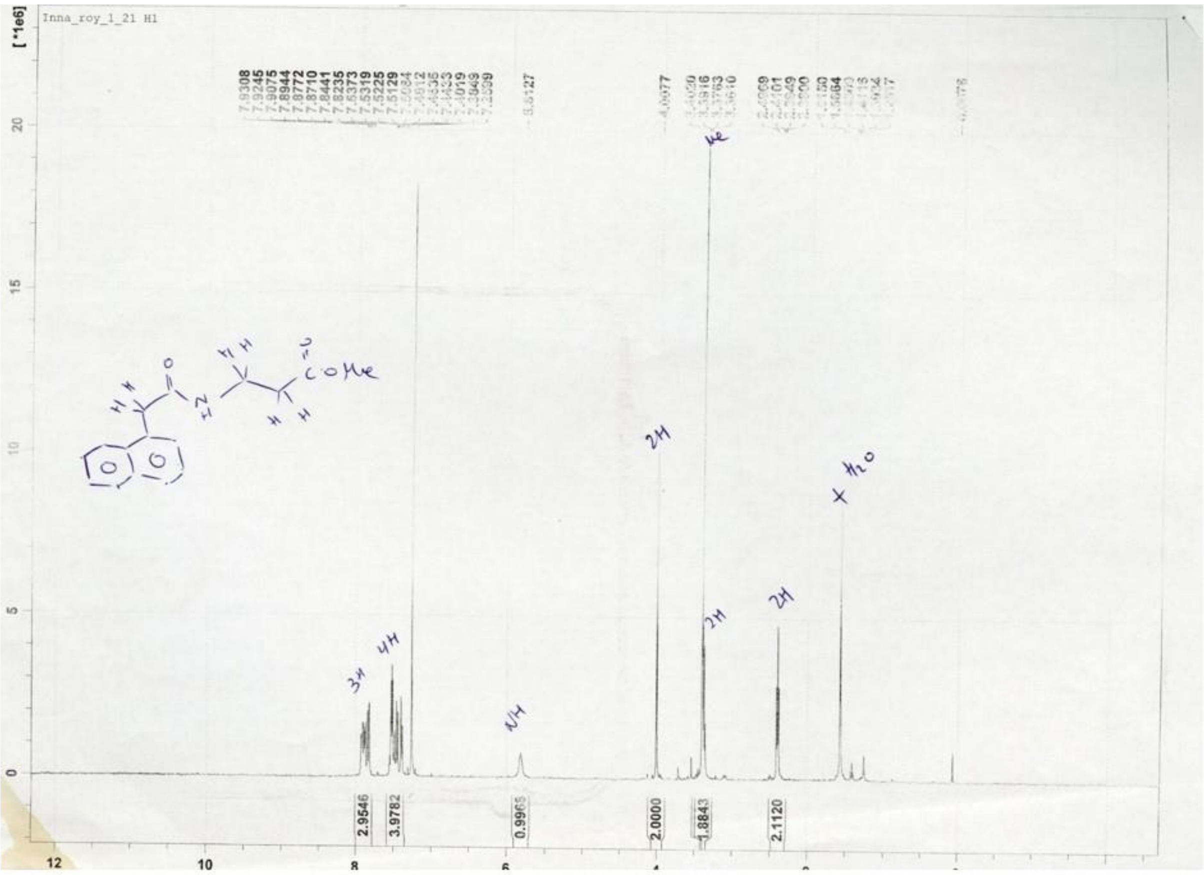

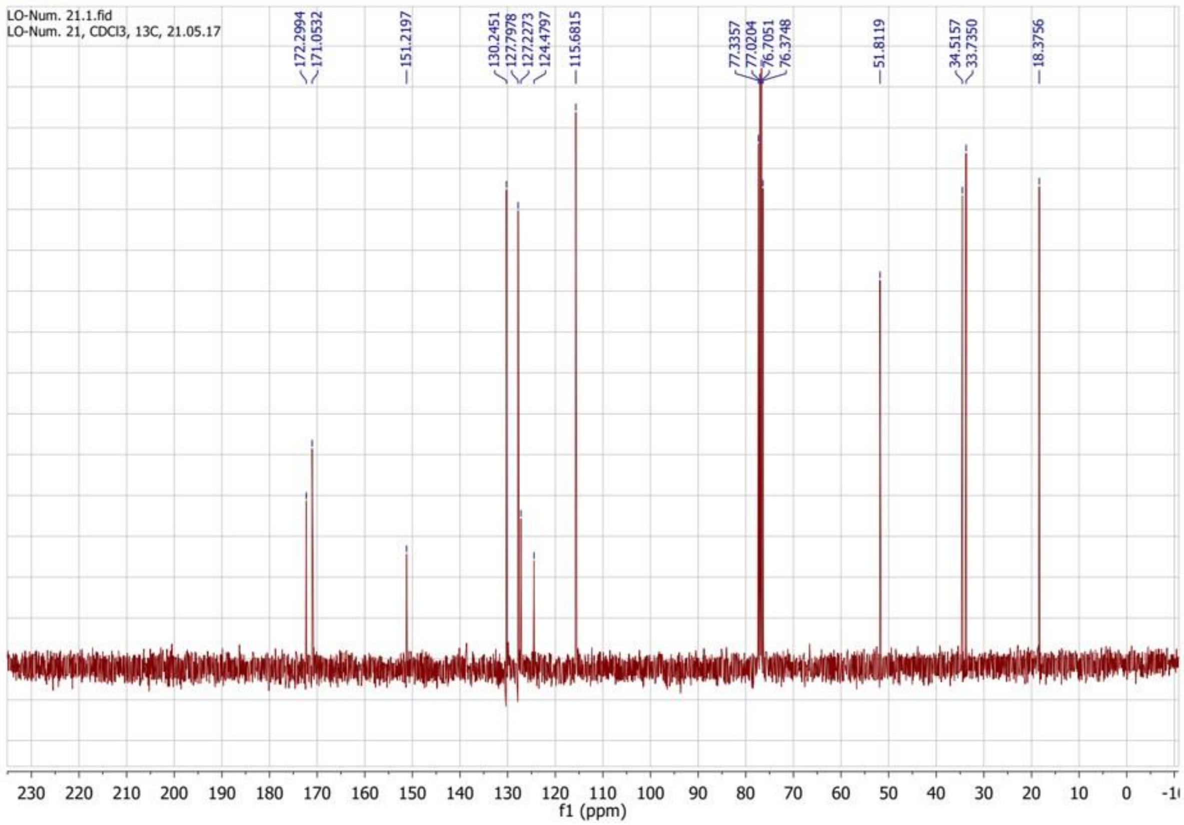

Figure 34. $^{1}H$ (top) and $^{13}C$ (bottom) NMR spectra of compound **4b**. 379

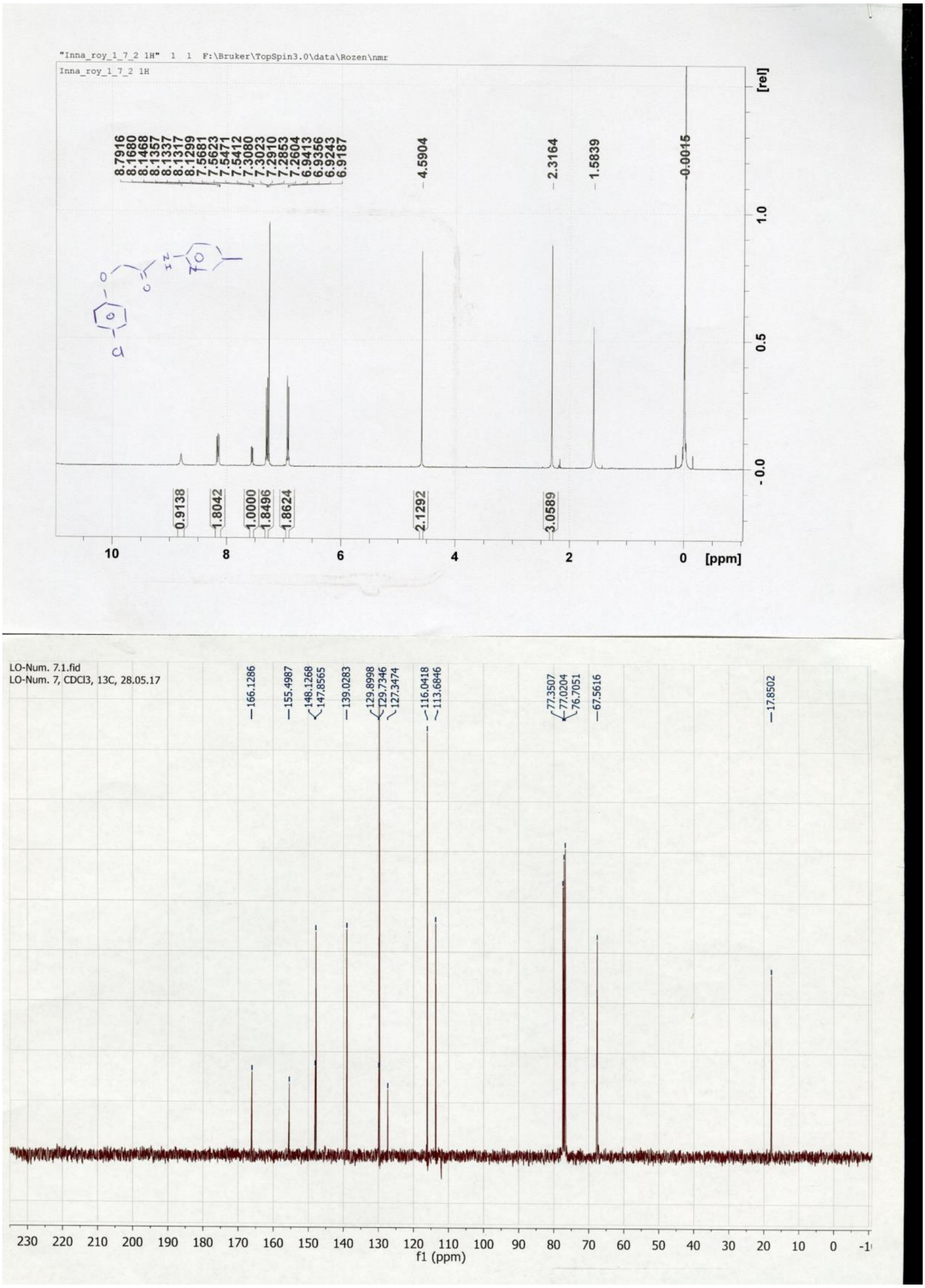


**Figure 35.** $^{1}H$ (top) and $^{13}C$ (bottom) NMR spectra of compound **1c**.

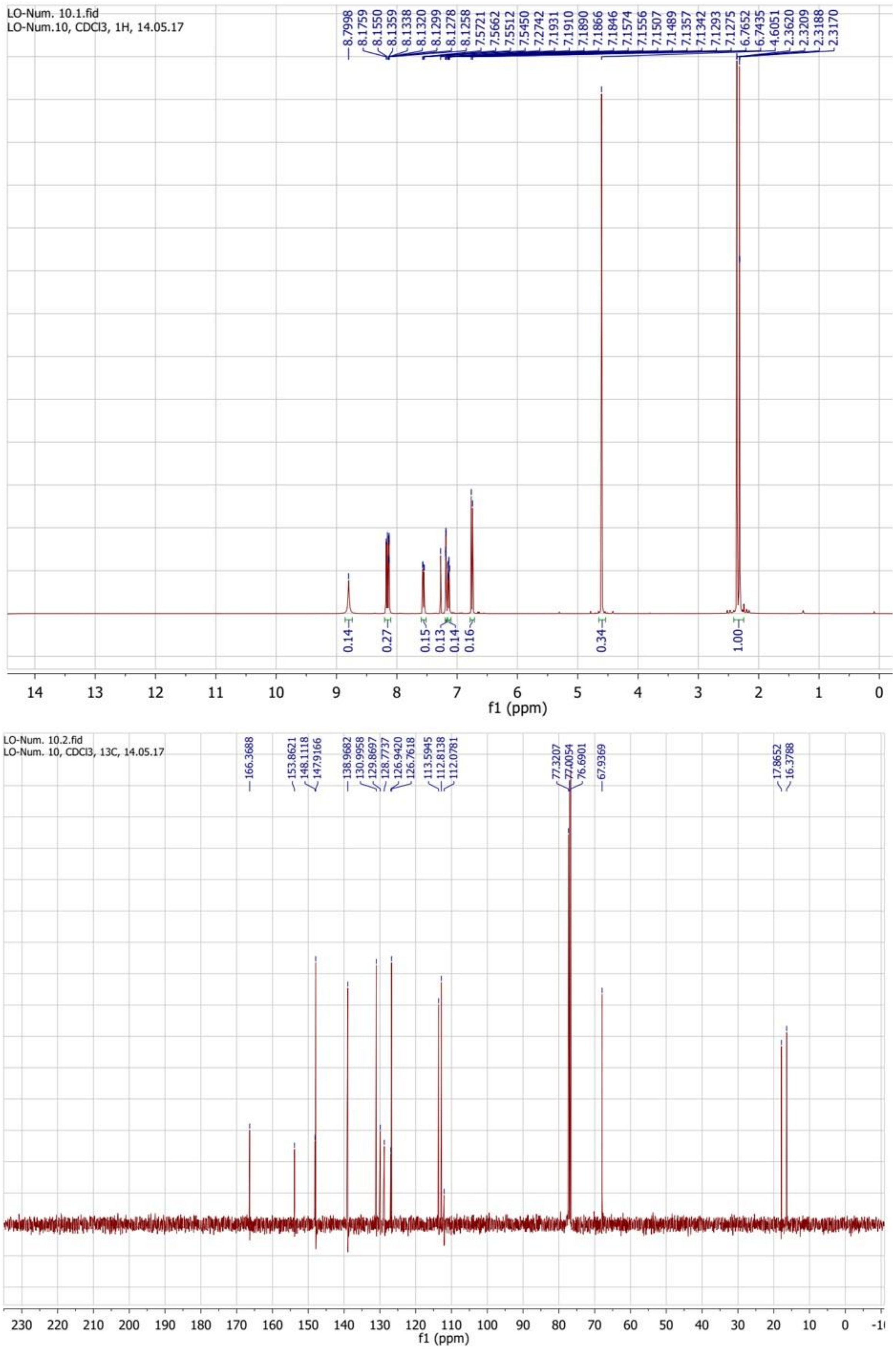


**Figure 36.** $^{1}$H (top) and $^{13}$C (bottom) NMR spectra of compound **2c**.

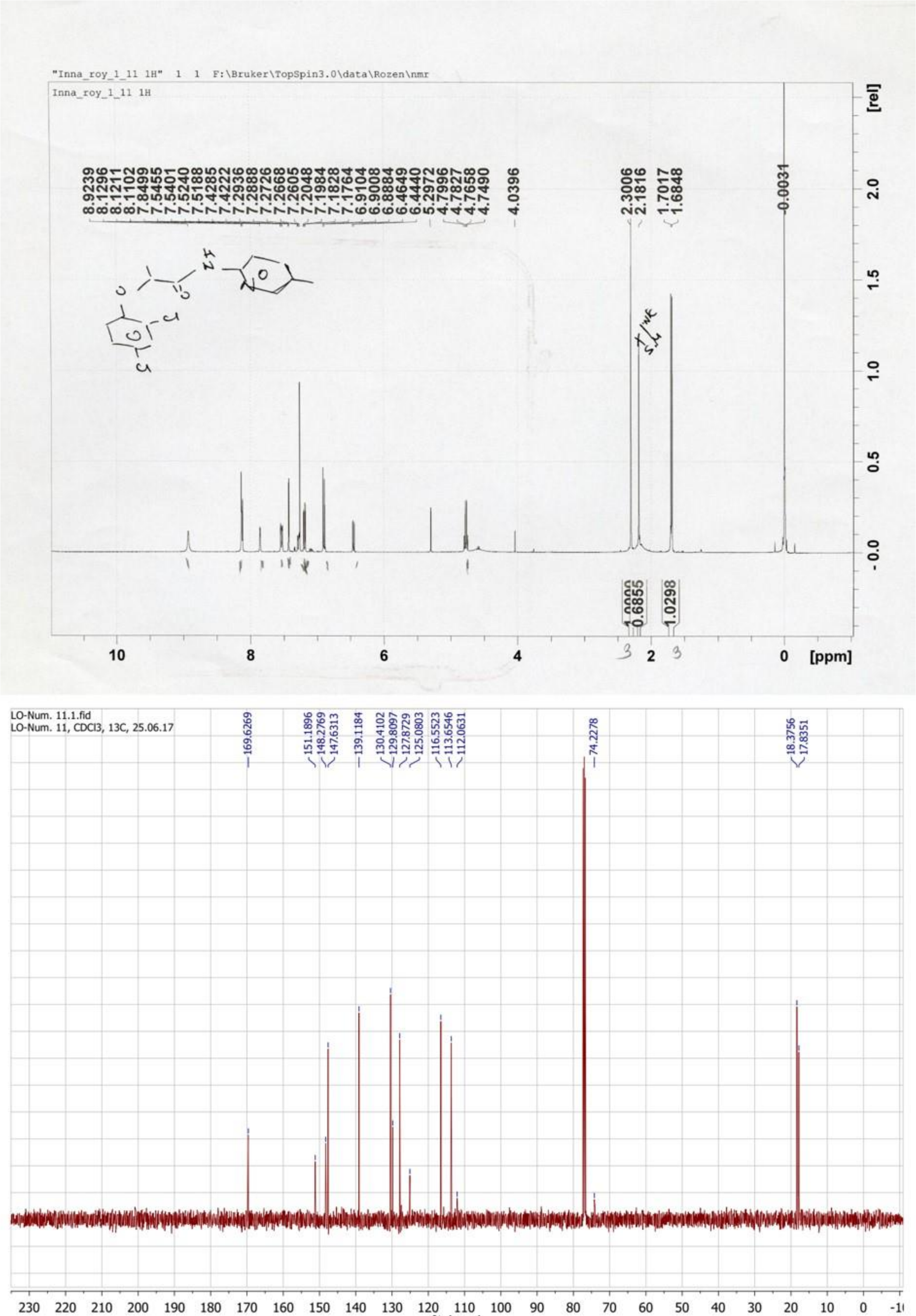


**Figure 37.** $^1$H (top) and $^{13}$C (bottom) NMR spectra of compound **3c**.

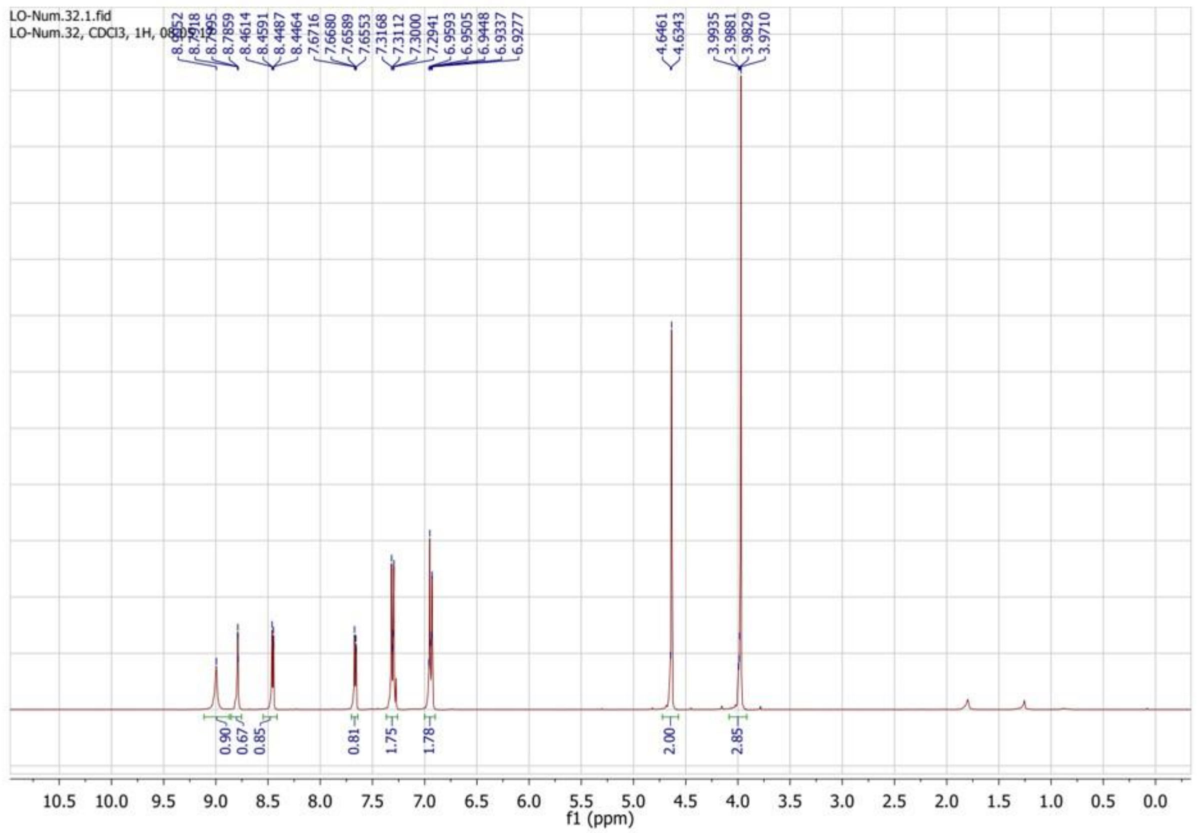


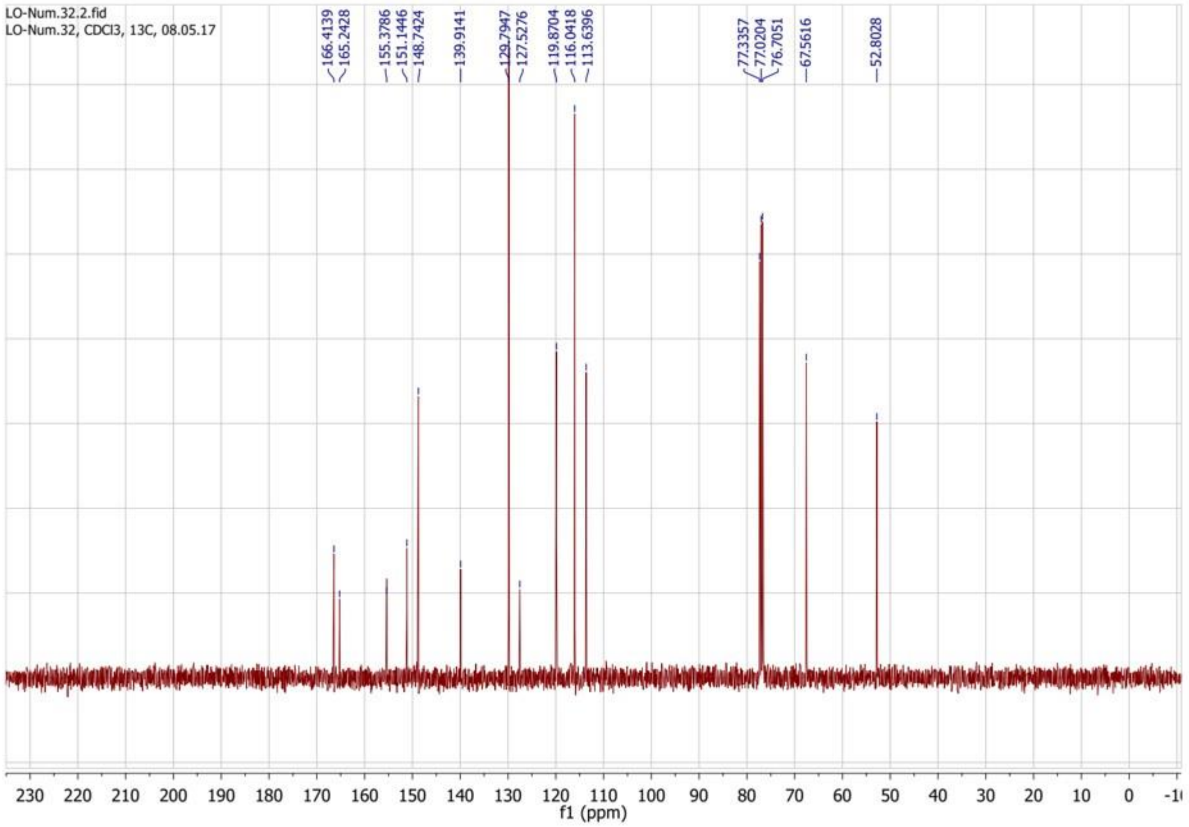


**Figure 38.** $^{1}H$ (top) and $^{13}C$ (bottom) NMR spectra of compound **1d**.

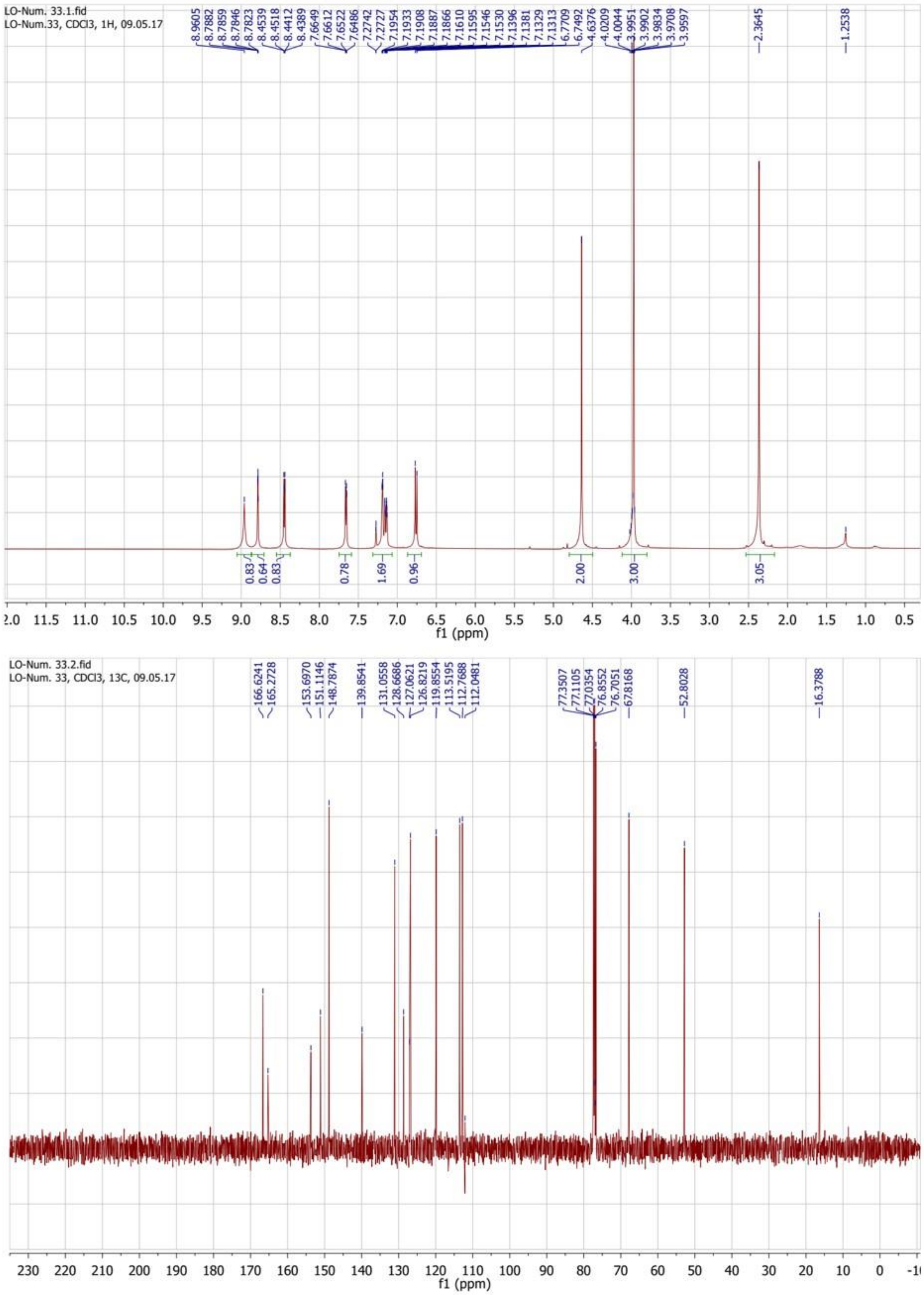


**Figure 39.** $^{1}H$ (top) and $^{13}C$ (bottom) NMR spectra of compound **2d**.

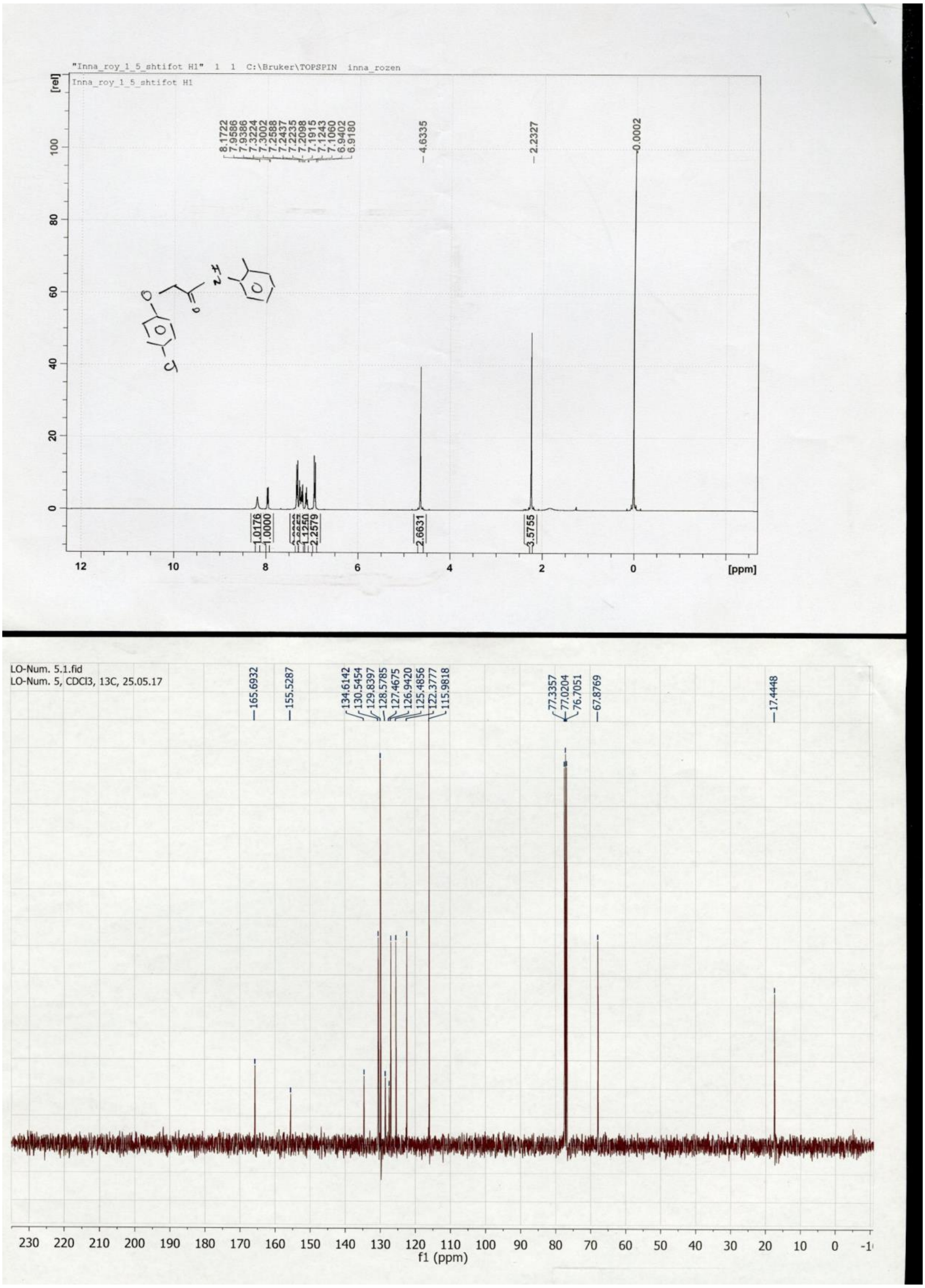


**Figure 40.** $^{1}$H (top) and $^{13}$C (bottom) NMR spectra of compound **1e**.

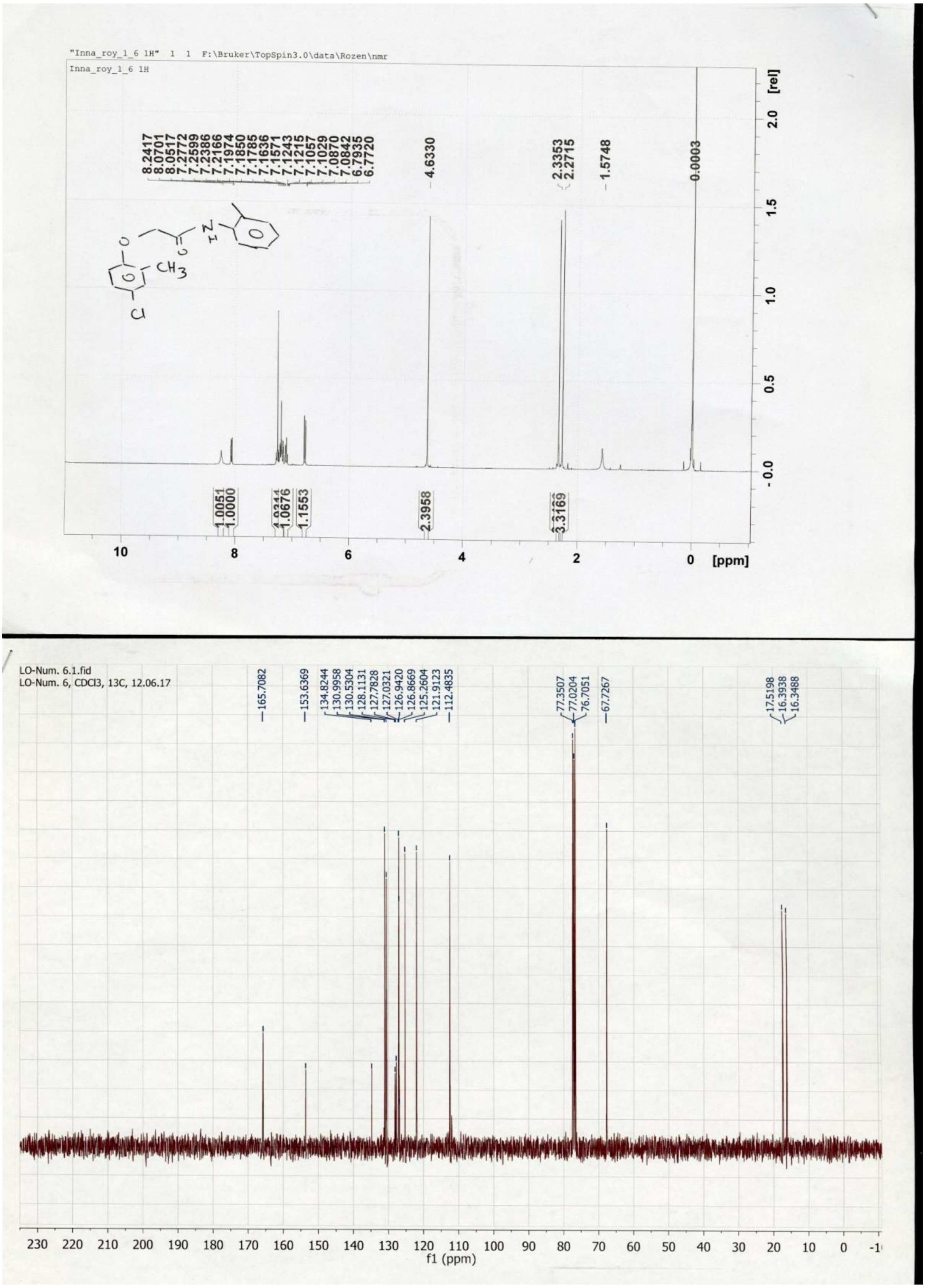


**Figure 41.** $^{1}H$ (top) and $^{13}C$ (bottom) NMR spectra of compound **2e**.

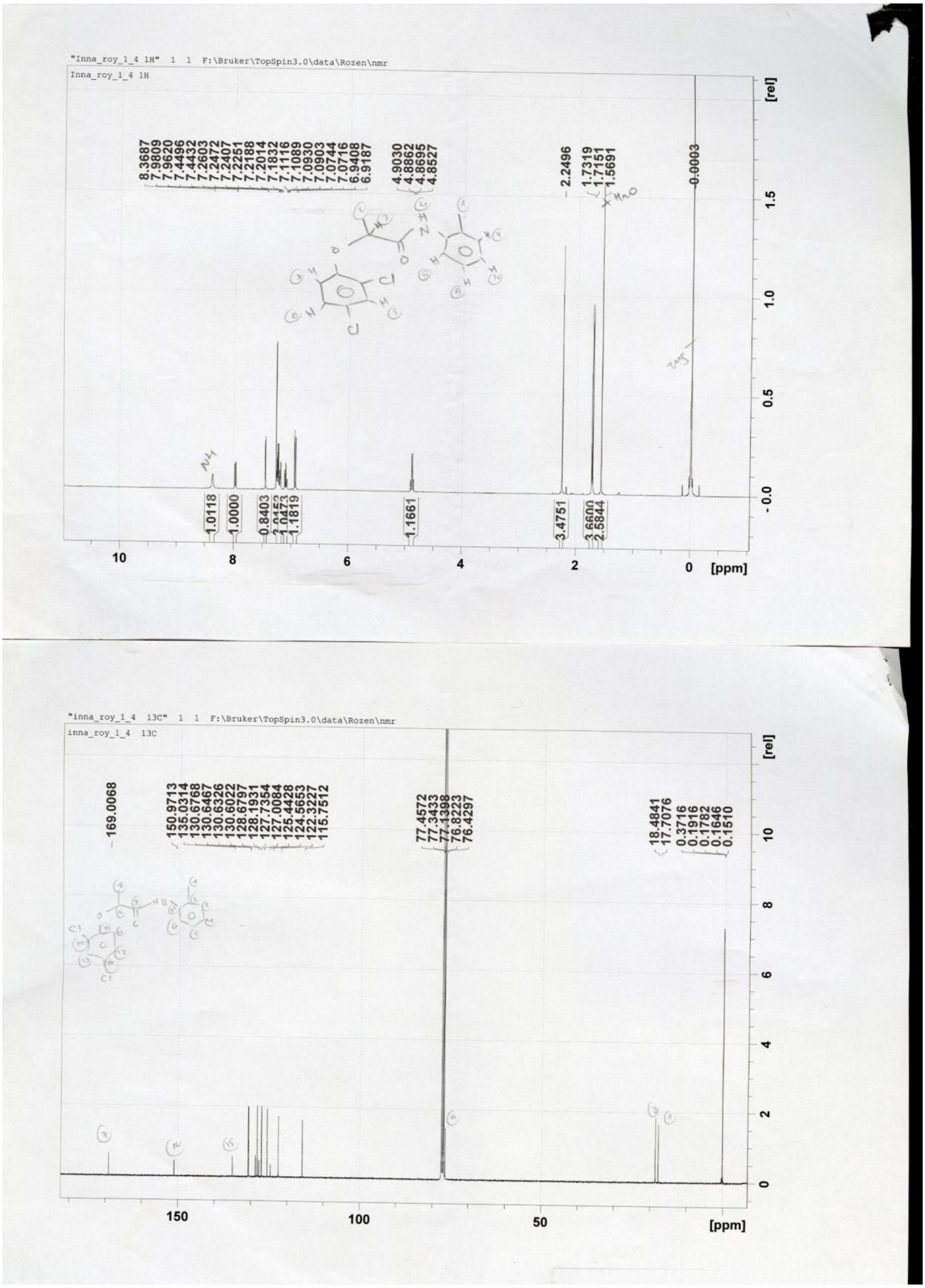


**Figure 42.** $^{1}$H (top) and $^{13}$C (bottom) NMR spectra of compound **3e**.

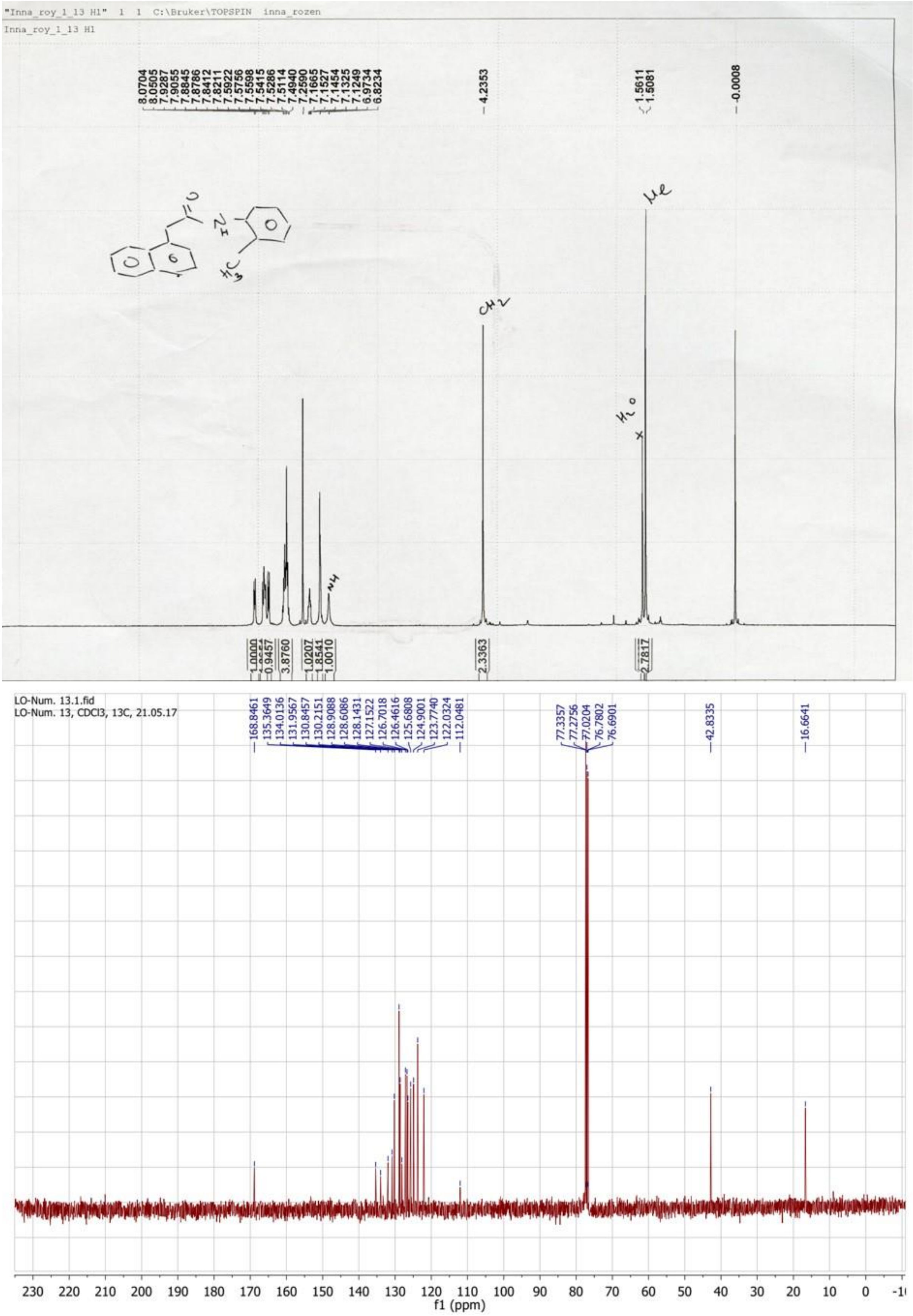


**Figure 43.** $^{1}H$ (top) and $^{13}C$ (bottom) NMR spectra of compound **4e**.

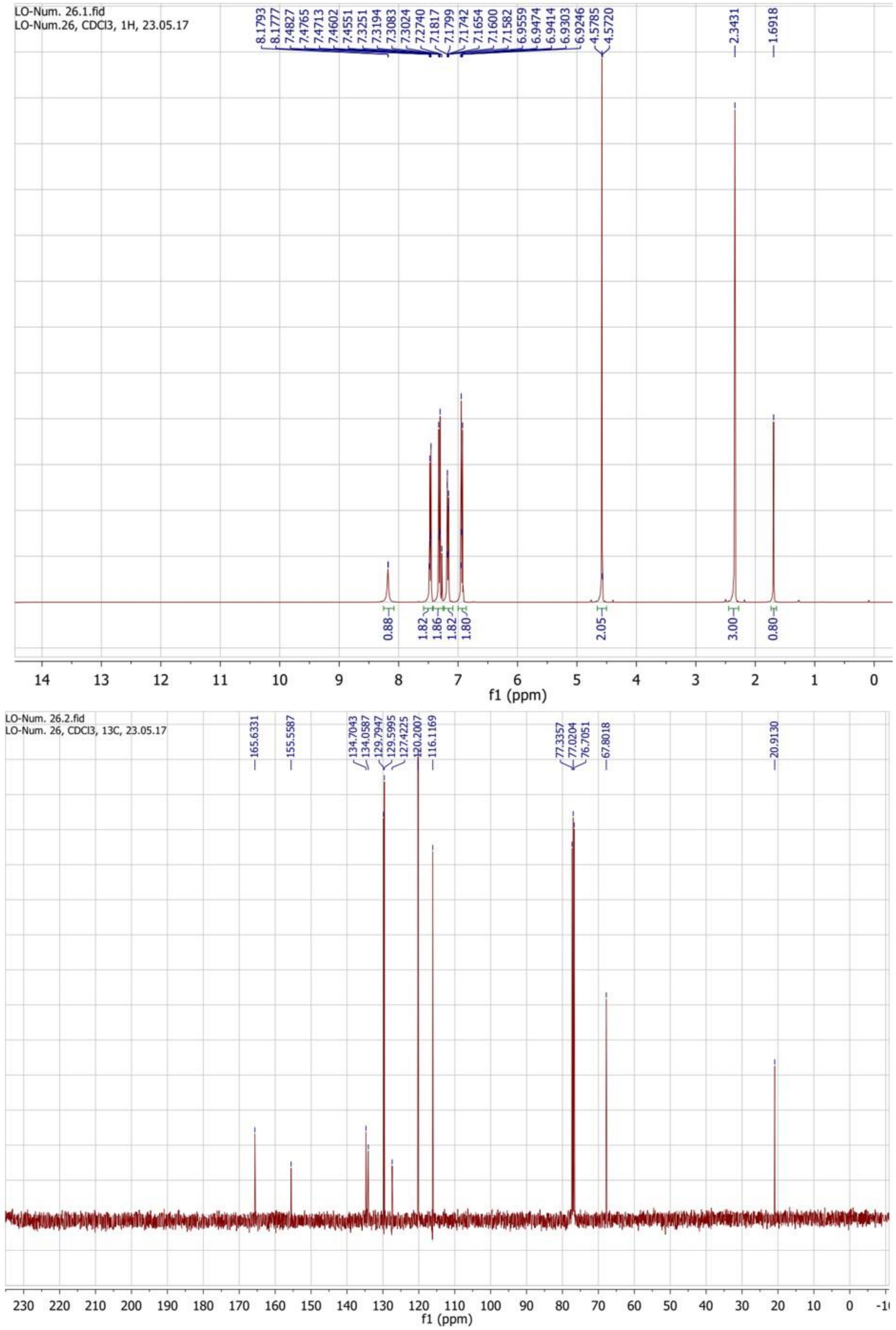


**Figure 44.** $^1$H (top) and $^{13}$C (bottom) NMR spectra of compound **1f**.

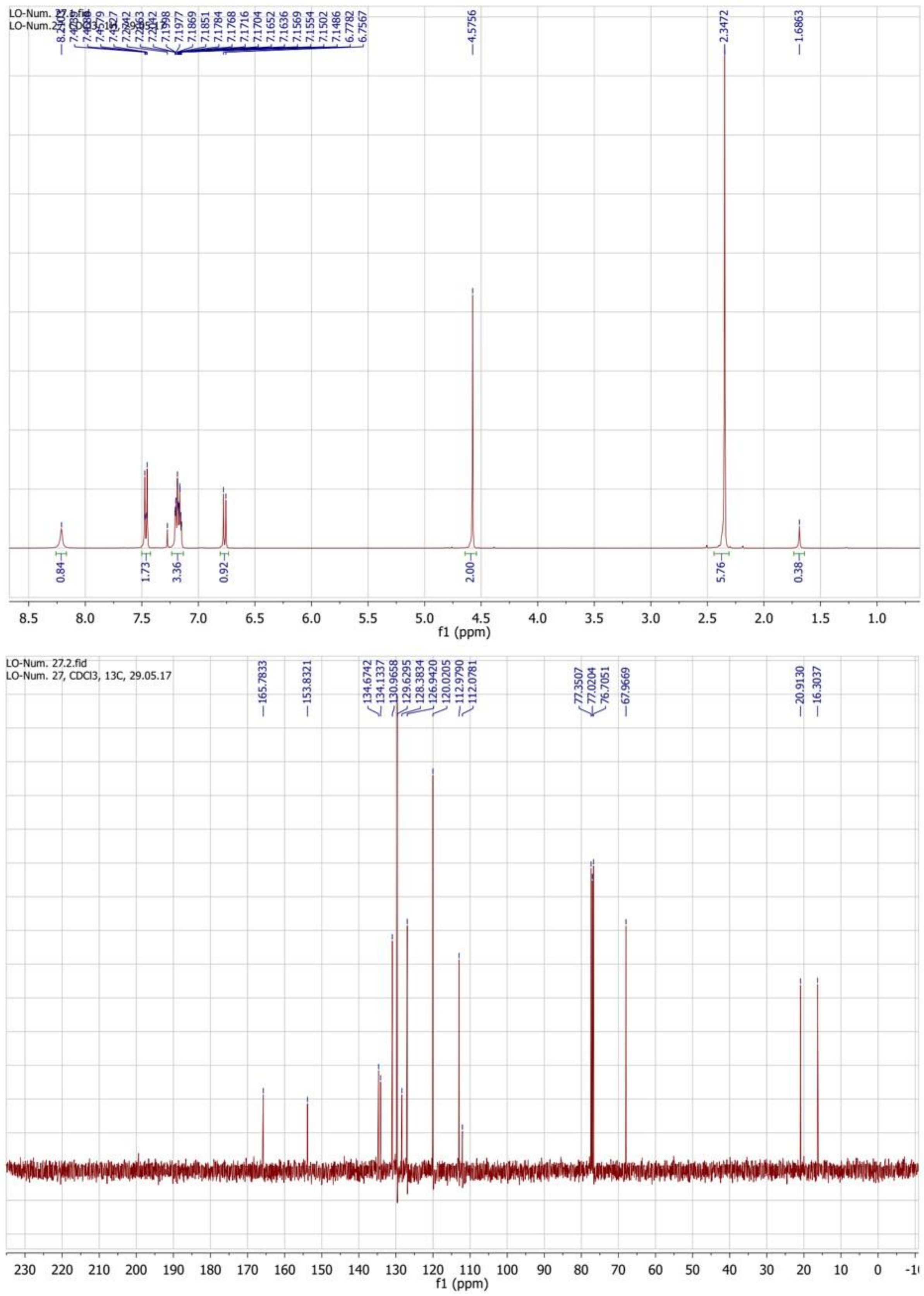


**Figure 45.** $^{1}H$ (top) and $^{13}C$ (bottom) NMR spectra of compound **2f**.

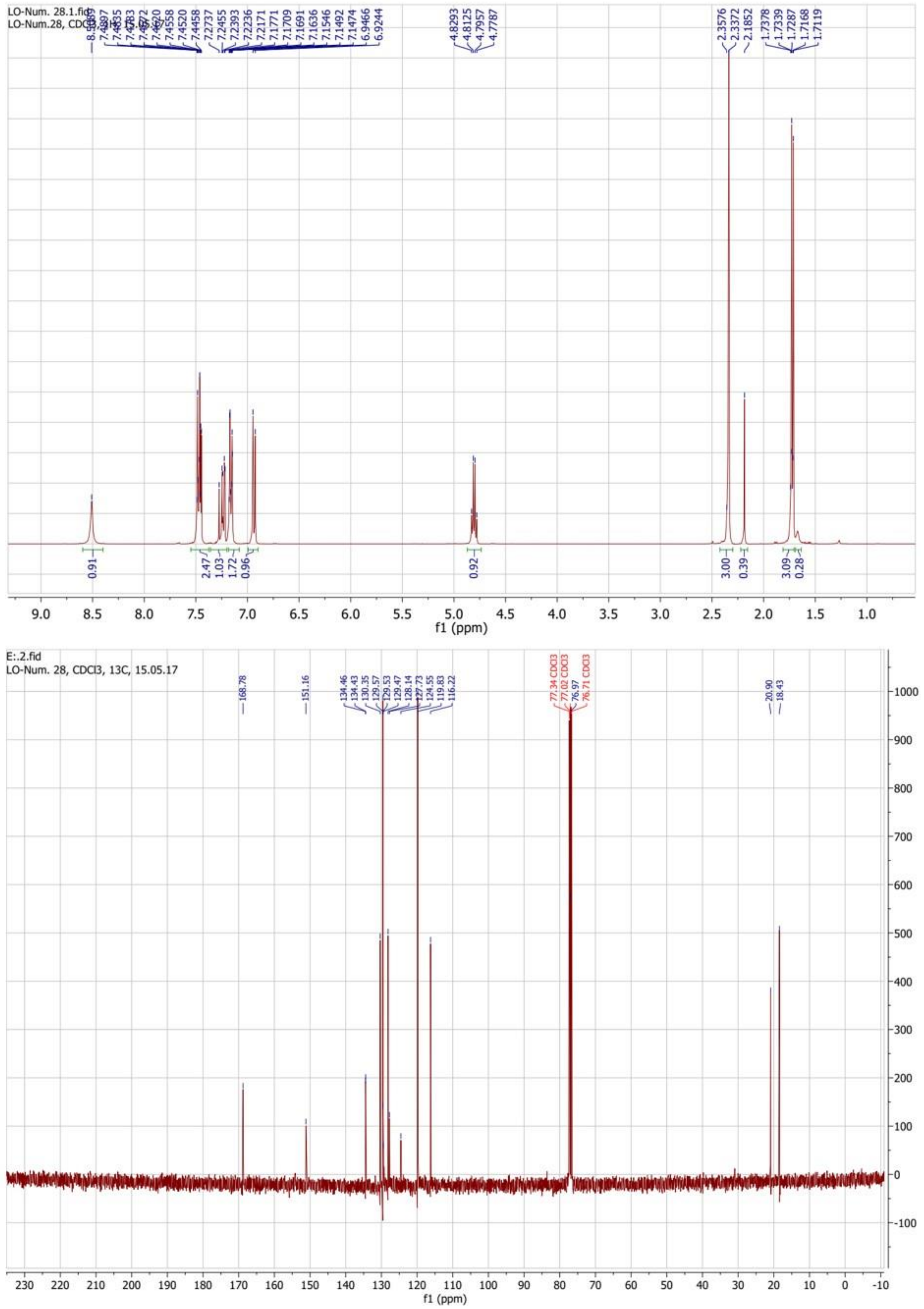


**Figure 46.** $^{1}$H (top) and $^{13}$C (bottom) NMR spectra of compound **3f**.

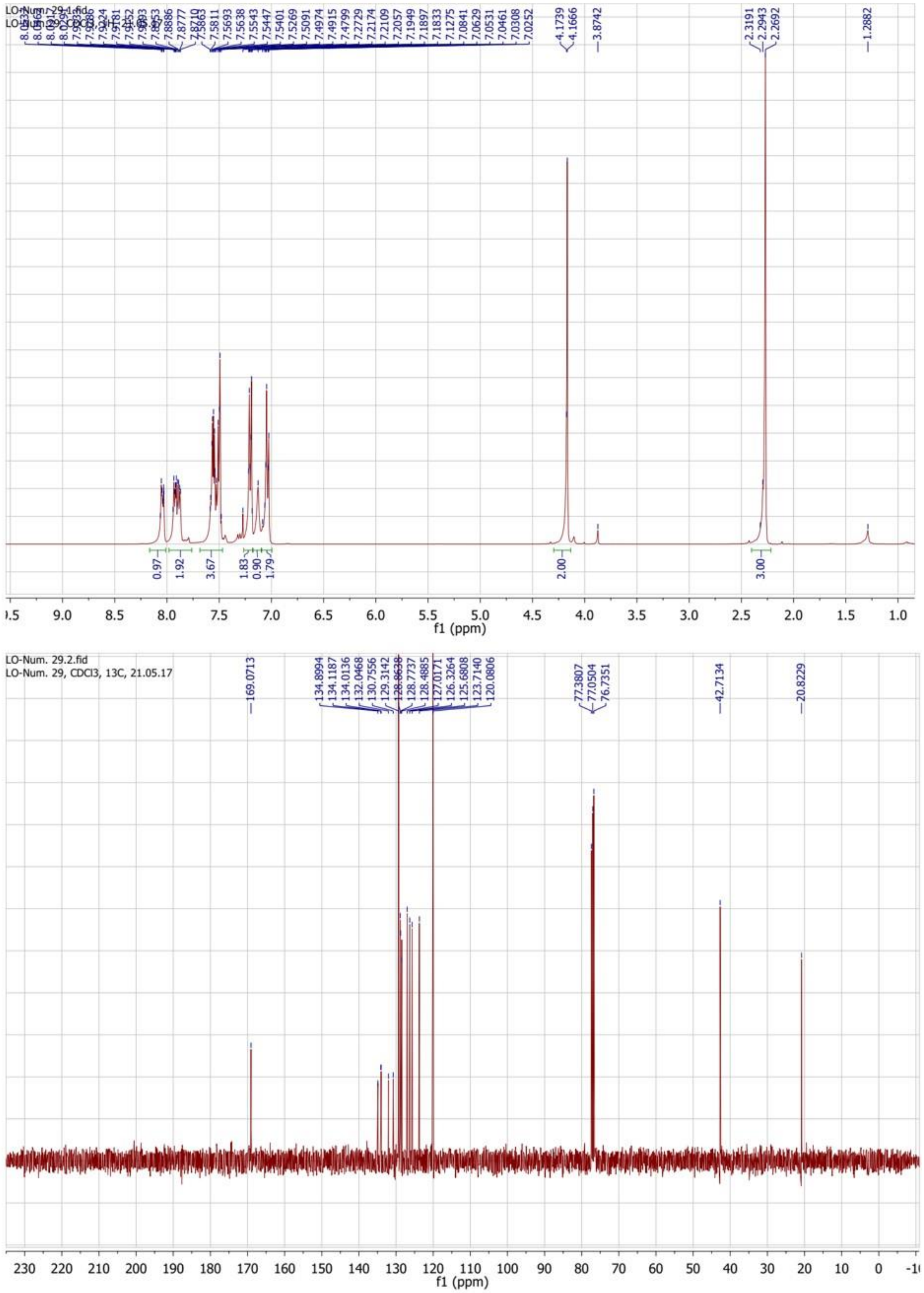


**Figure 47.** $^1$H (top) and $^{13}$C (bottom) NMR spectra of compound **4f**.

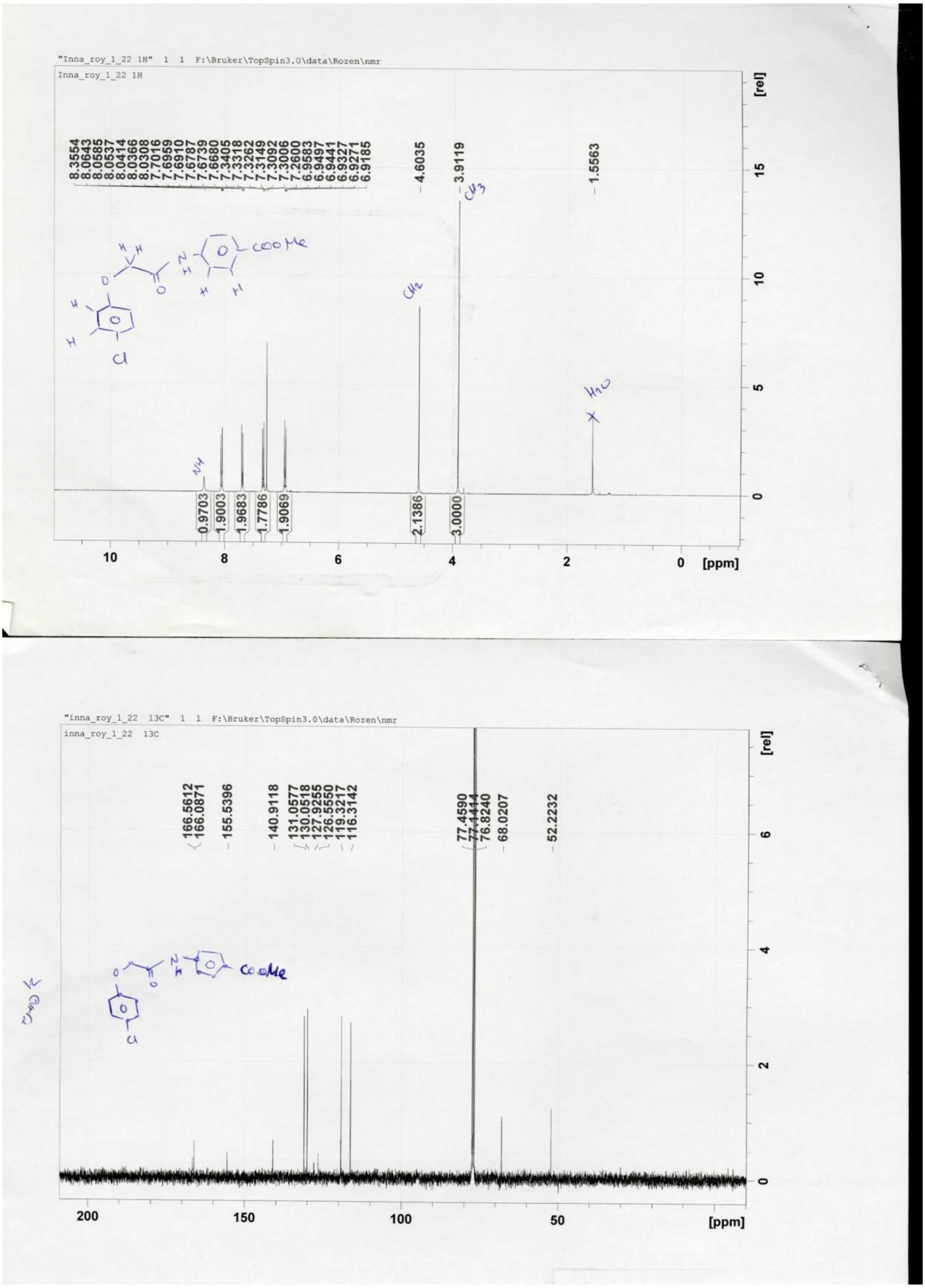

**Figure 48.** $^{1}$H (top) and $^{13}$C (bottom) NMR spectra of compound **1g**.

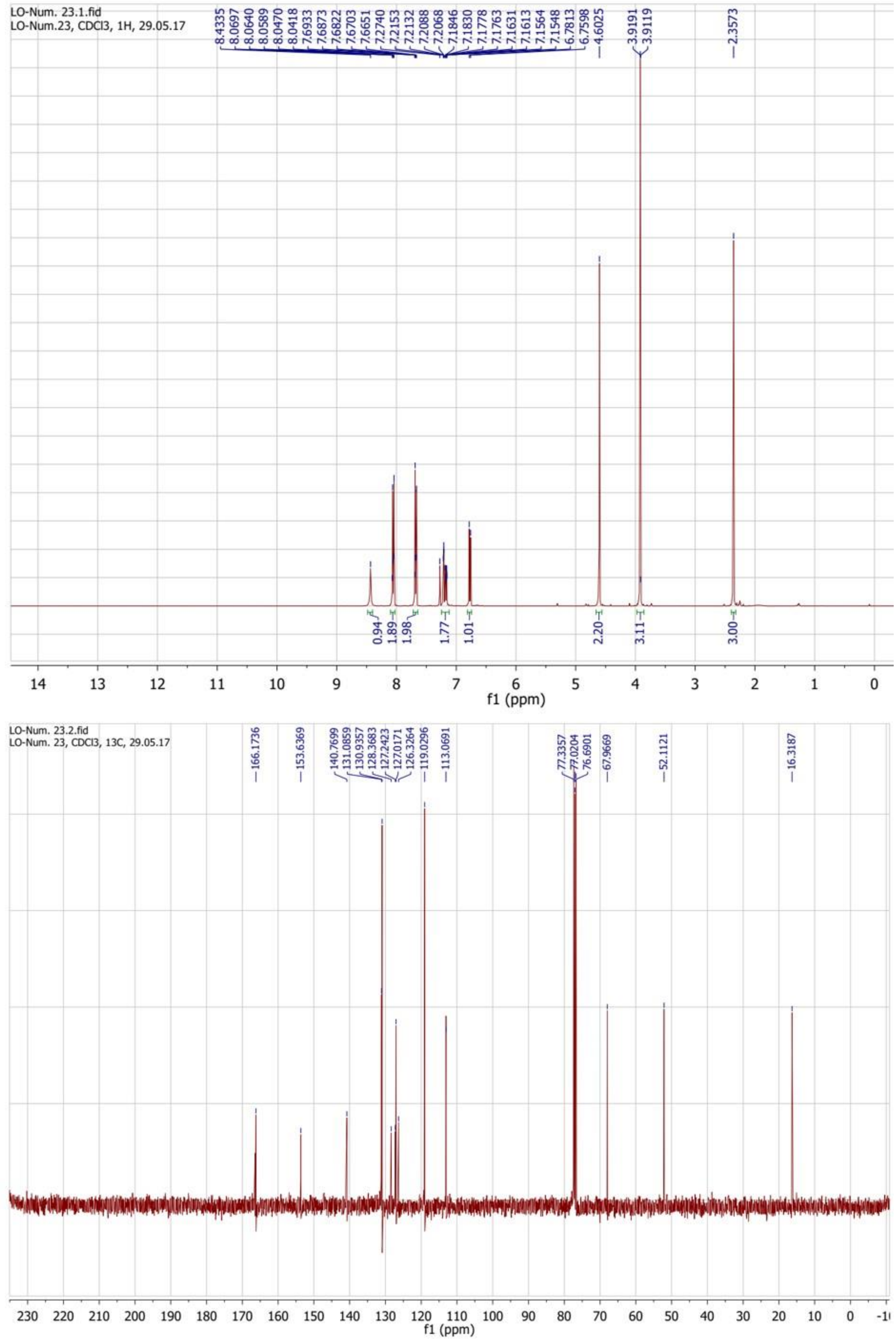


**Figure 49.** $^1$H (top) and $^{13}$C (bottom) NMR spectra of compound **2g**.

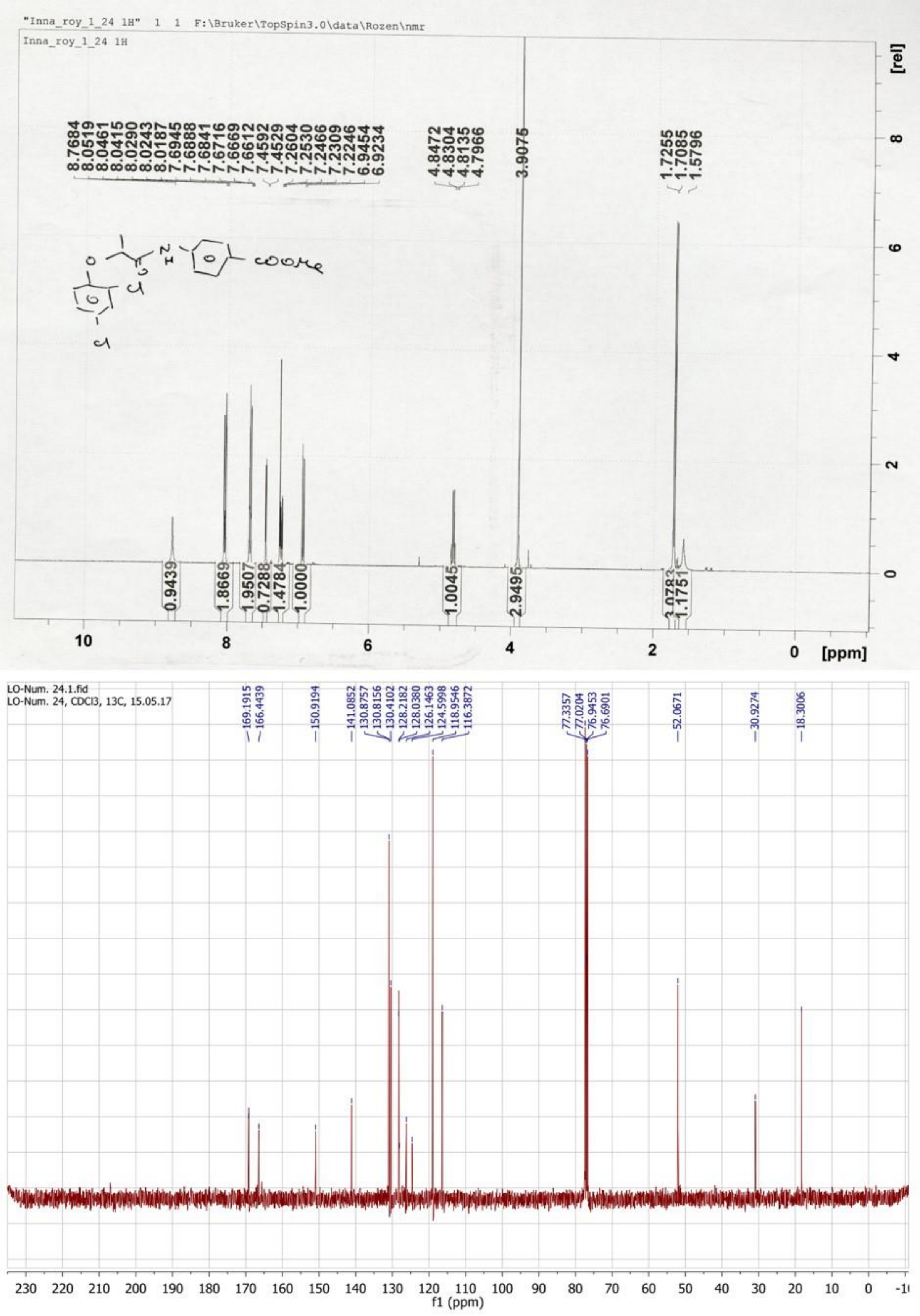


**Figure 50.** $^{1}H$ (top) and $^{13}C$ (bottom) NMR spectra of compound **3g**.

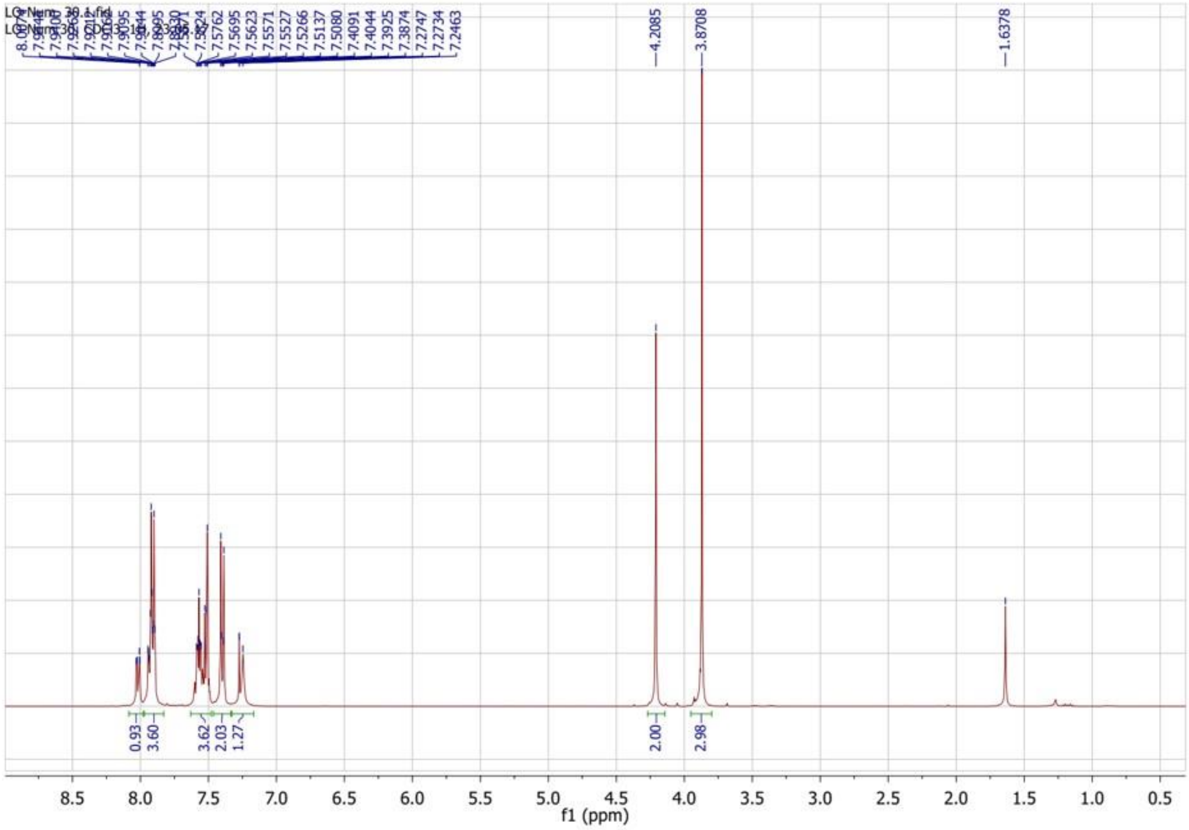


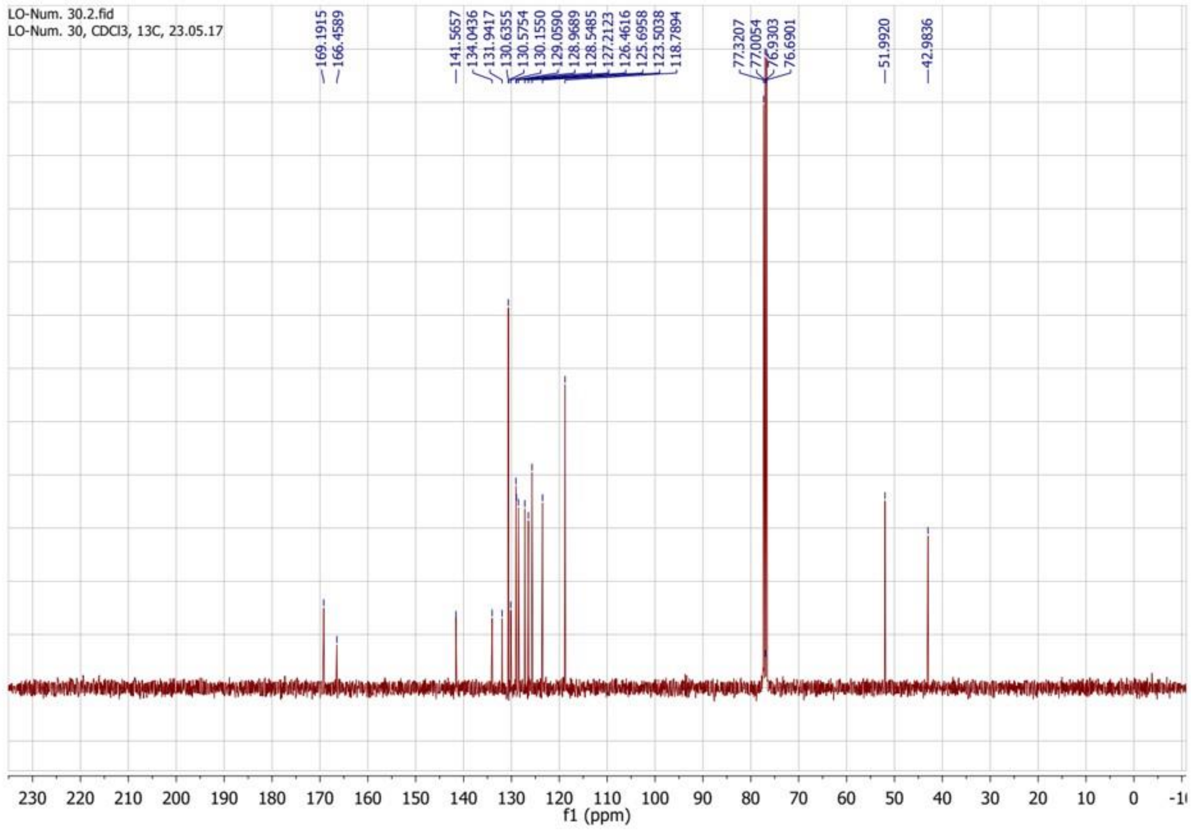


**Figure 51.** $^{1}H$ (top) and $^{13}C$ (bottom) NMR spectra of compound **4g**.

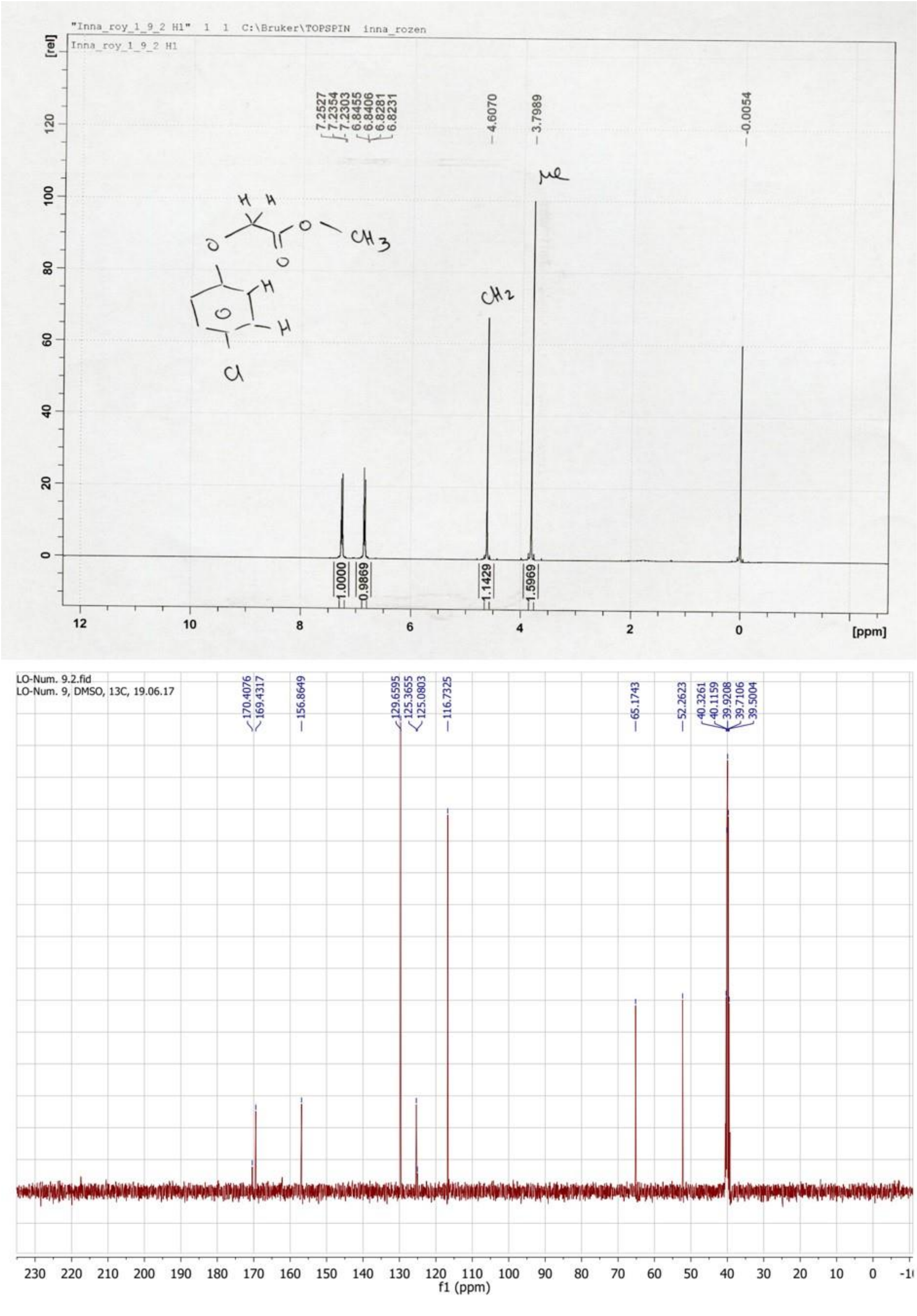


**Figure 52.** $^1H$ (top) and $^{13}C$ (bottom) NMR spectra of compound **1h**.

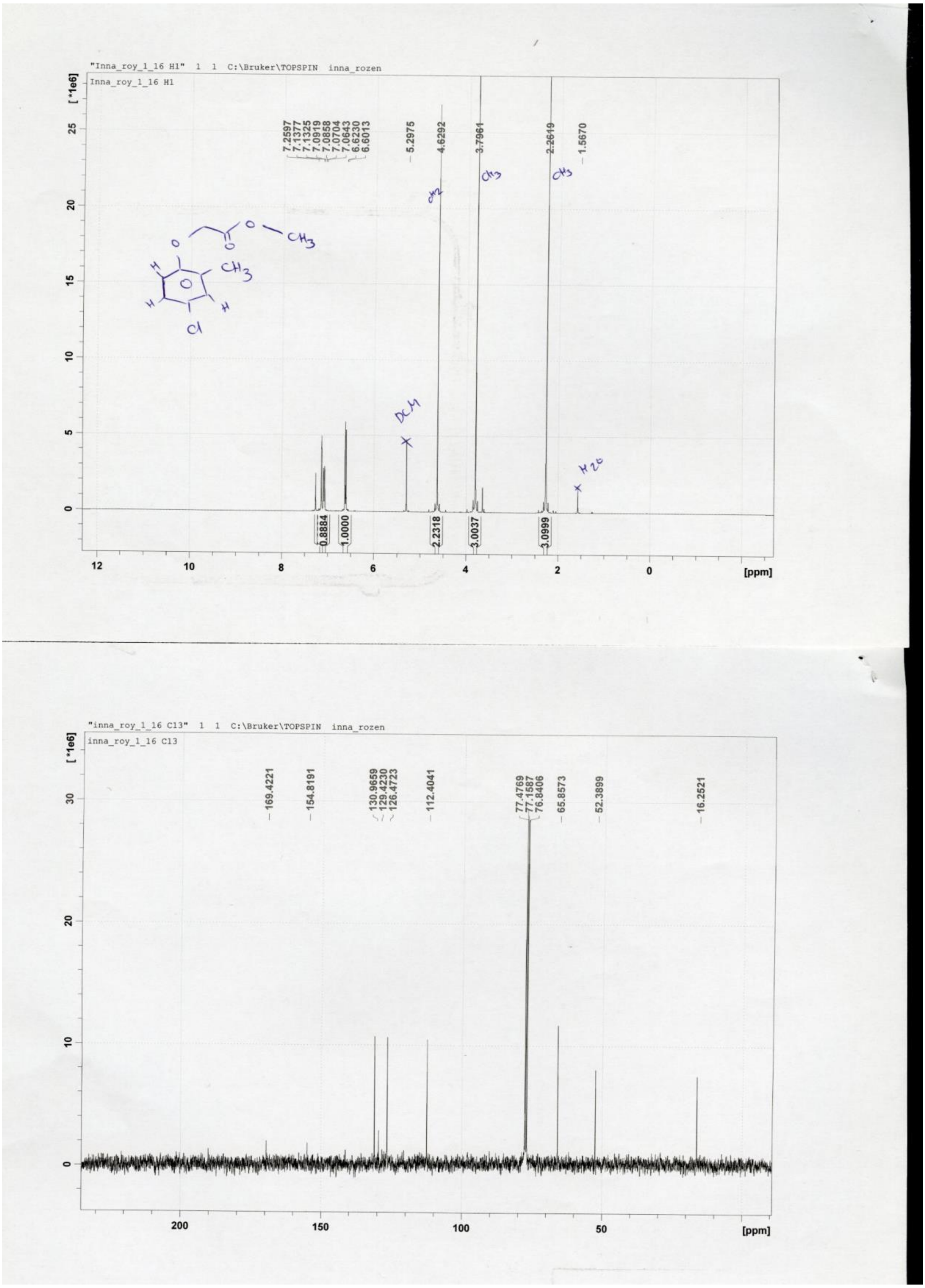


**Figure 53.** $^1$H (top) and $^{13}$C (bottom) NMR spectra of compound **2h**.

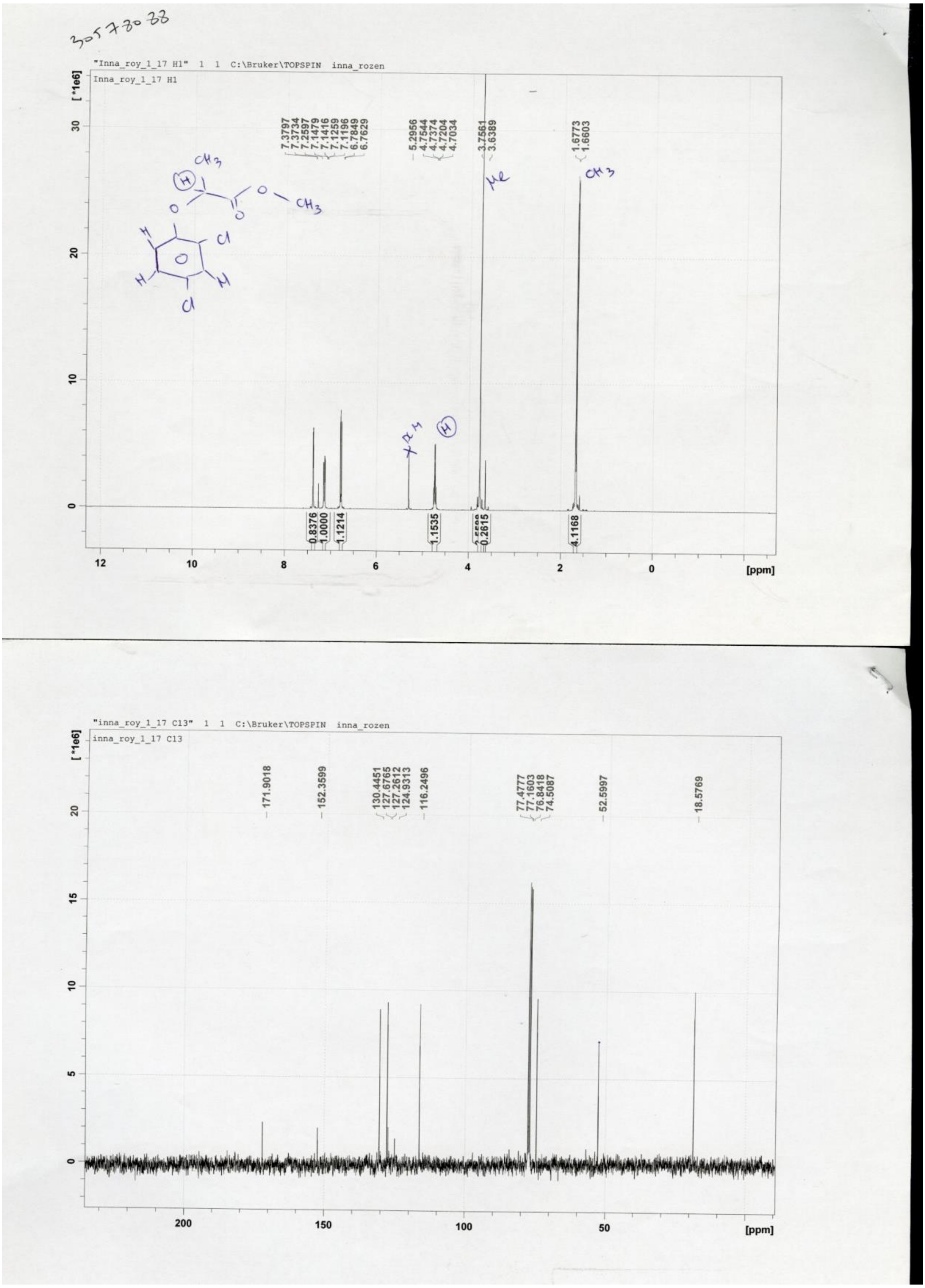


**Figure 54.** $^{1}H$ (top) and $^{13}C$ (bottom) NMR spectra of compound **3h**.

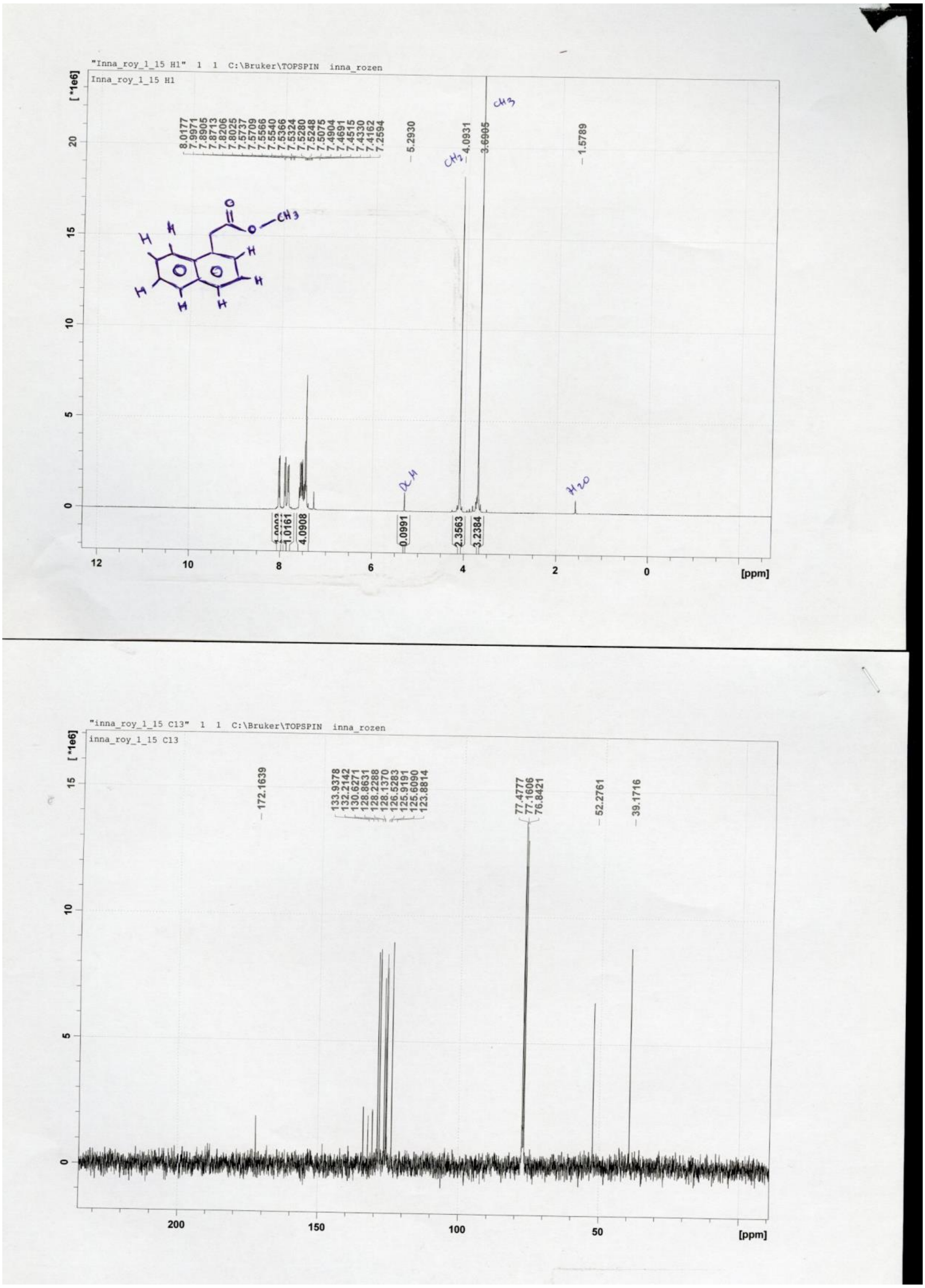


**Figure 55.** $^1$H (top) and $^{13}$C (bottom) NMR spectra of compound **4h**.

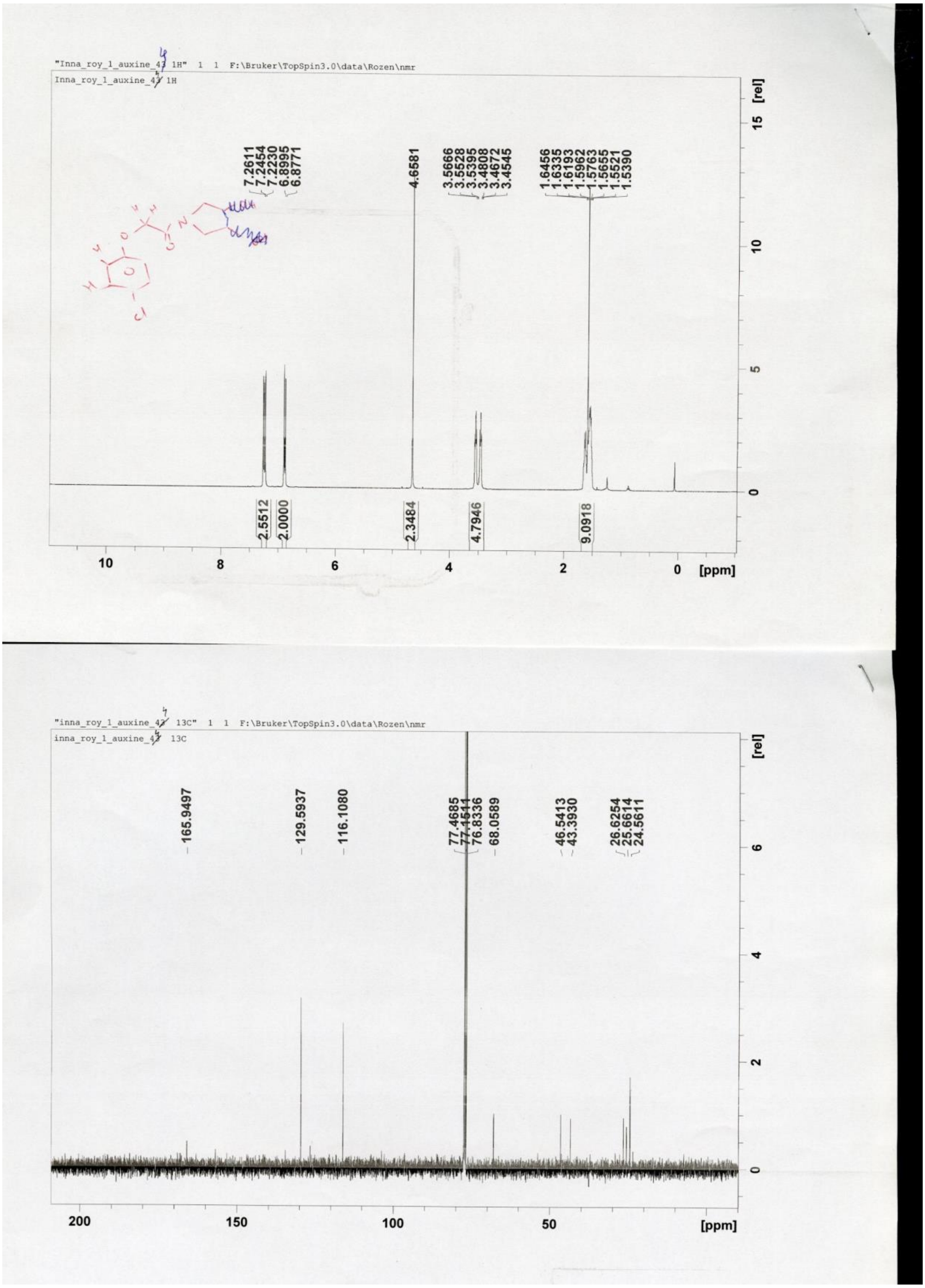


**Figure 56.** $^{1}H$ (top) and $^{13}C$ (bottom) NMR spectra of compound **1i**.

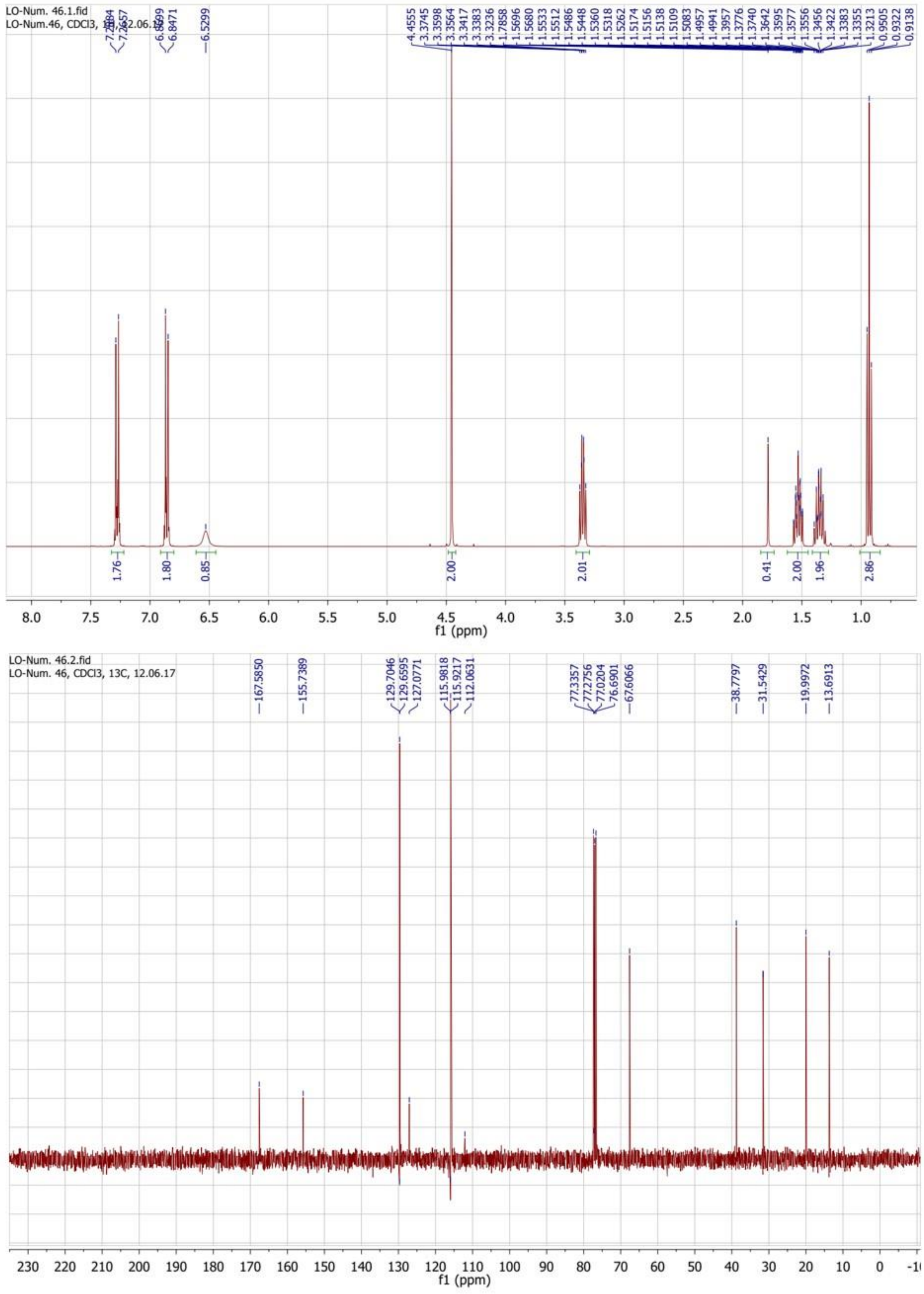


**Figure 57.** $^{1}H$ (top) and $^{13}C$ (bottom) NMR spectra of compound **1j**.

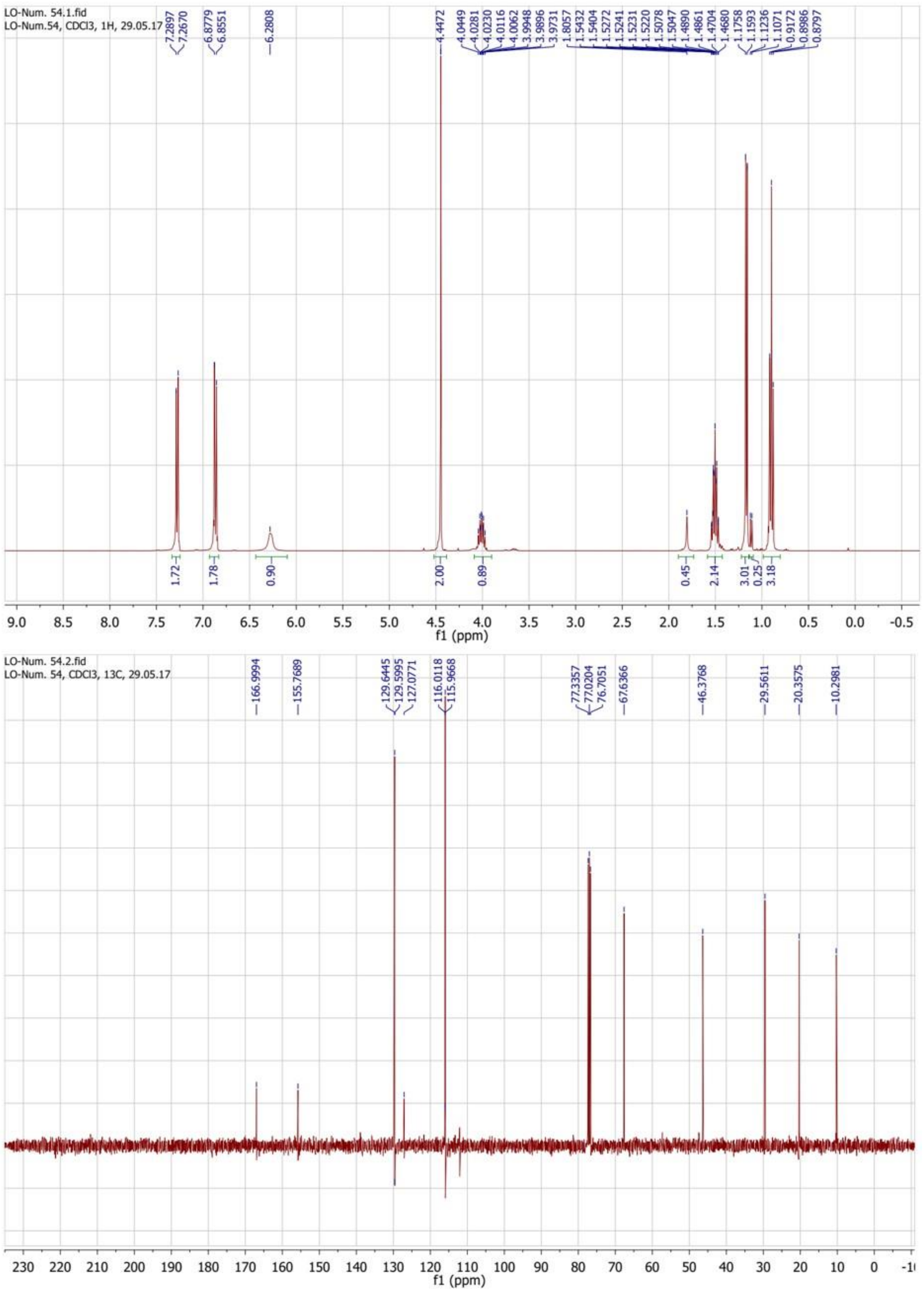


**Figure 58.** $^{1}H$ (top) and $^{13}C$ (bottom) NMR spectra of compound **1k**.

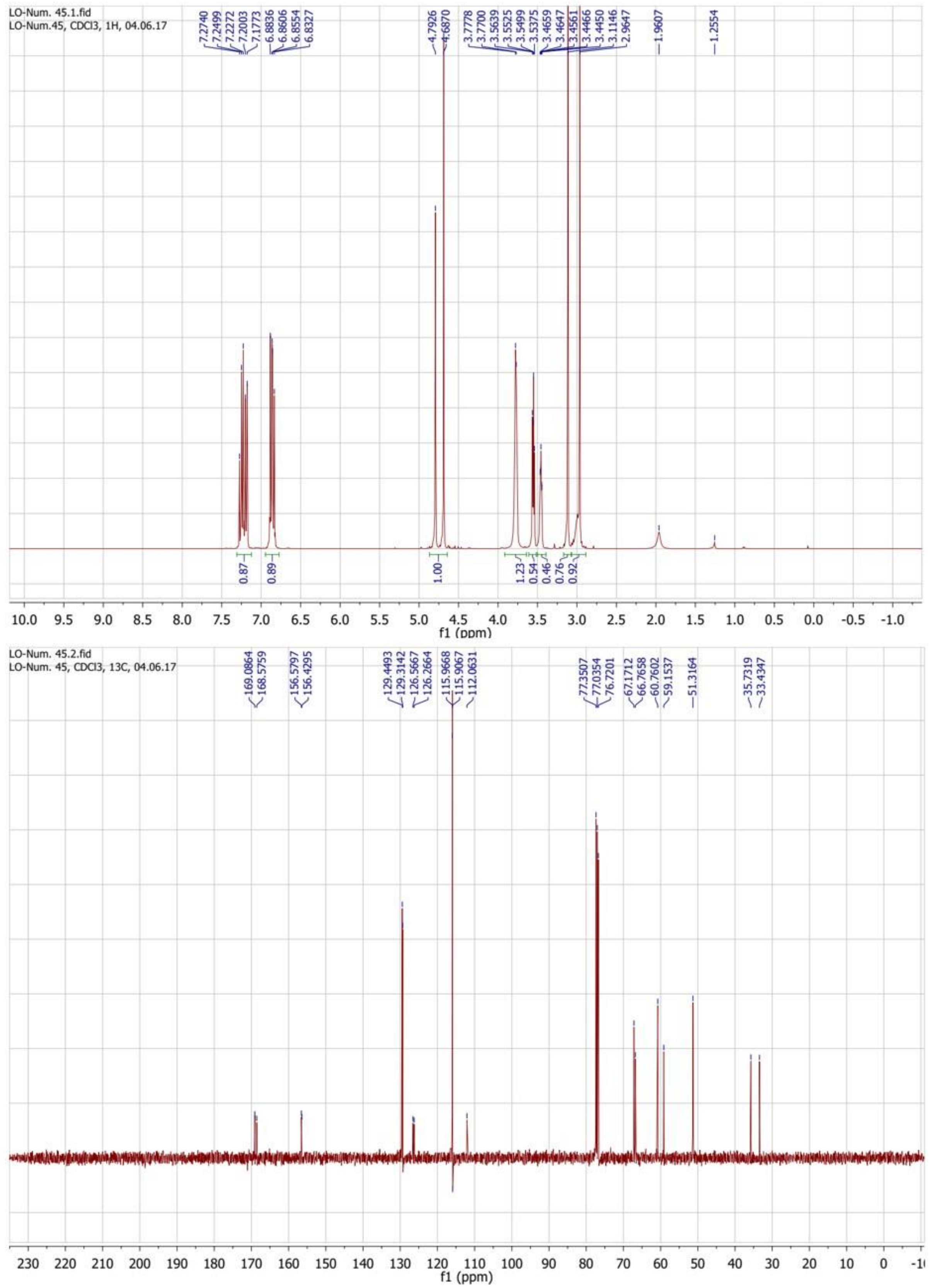


**Figure 59.** $^{1}H$ (top) and $^{13}C$ (bottom) NMR spectra of compound **1l**.

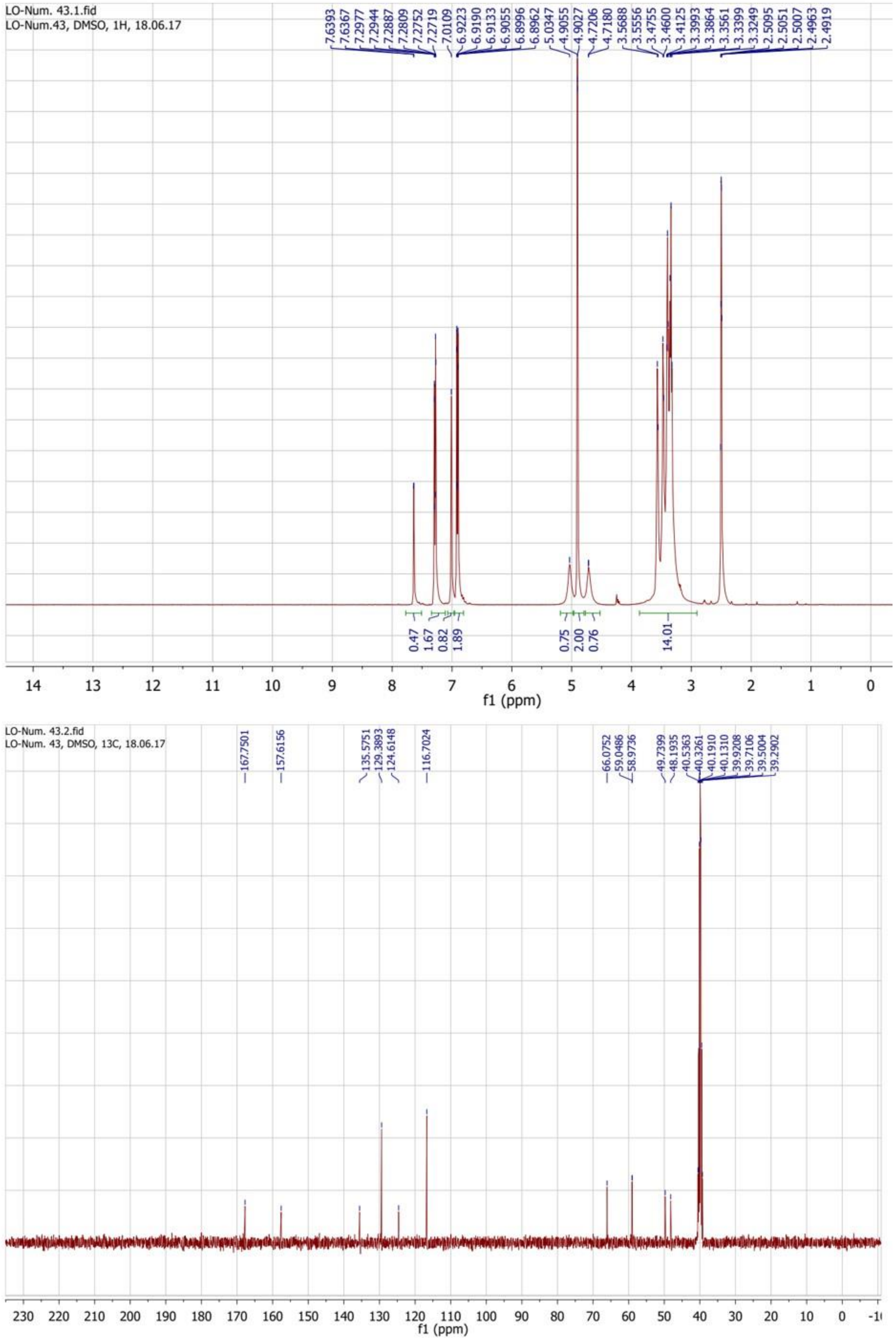


**Figure 60.** $^{1}H$ (top) and $^{13}C$ (bottom) NMR spectra of compound **1m**.

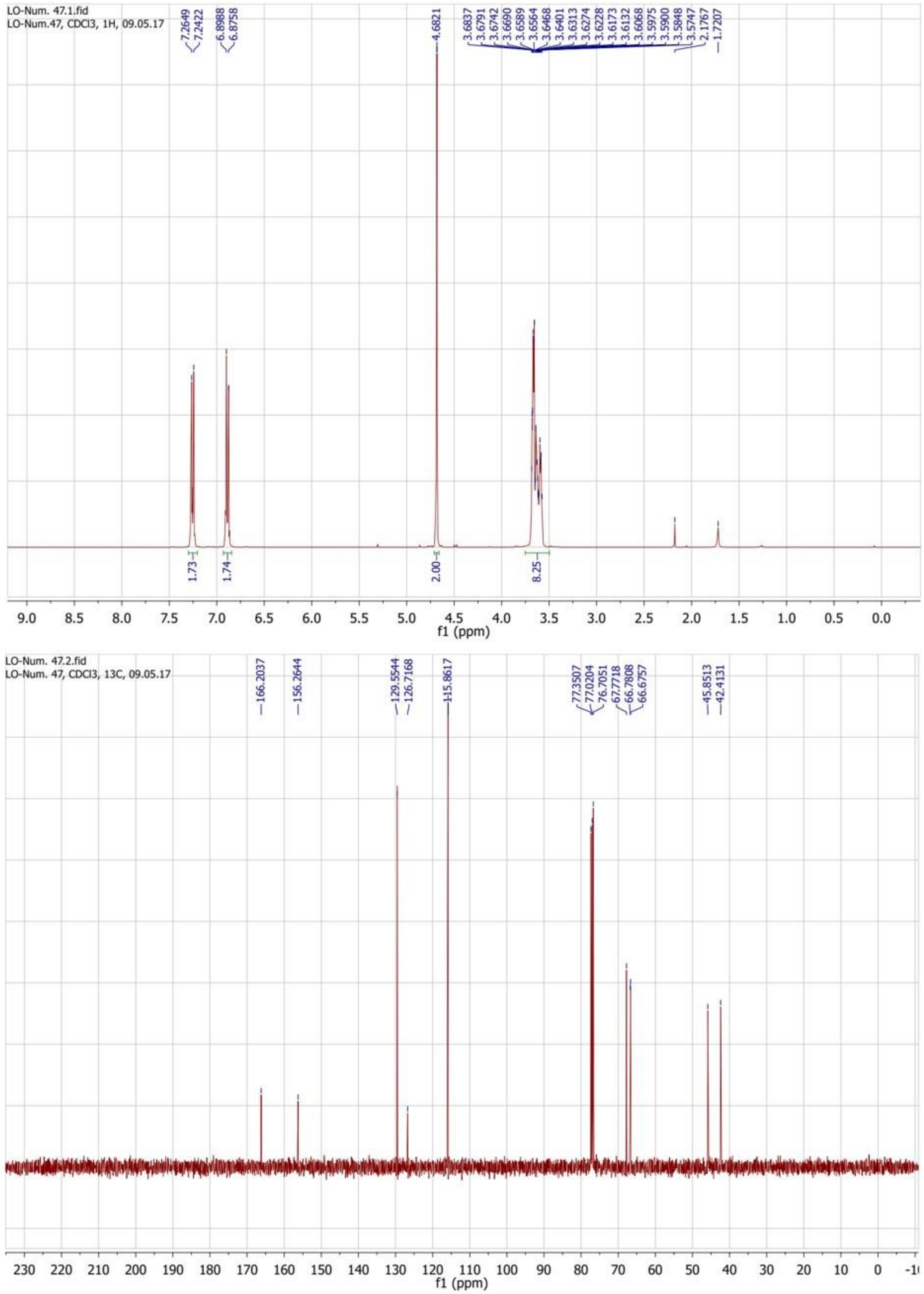


**Figure 61.** $^{1}$H (top) and $^{13}$C (bottom) NMR spectra of compound **1n**.

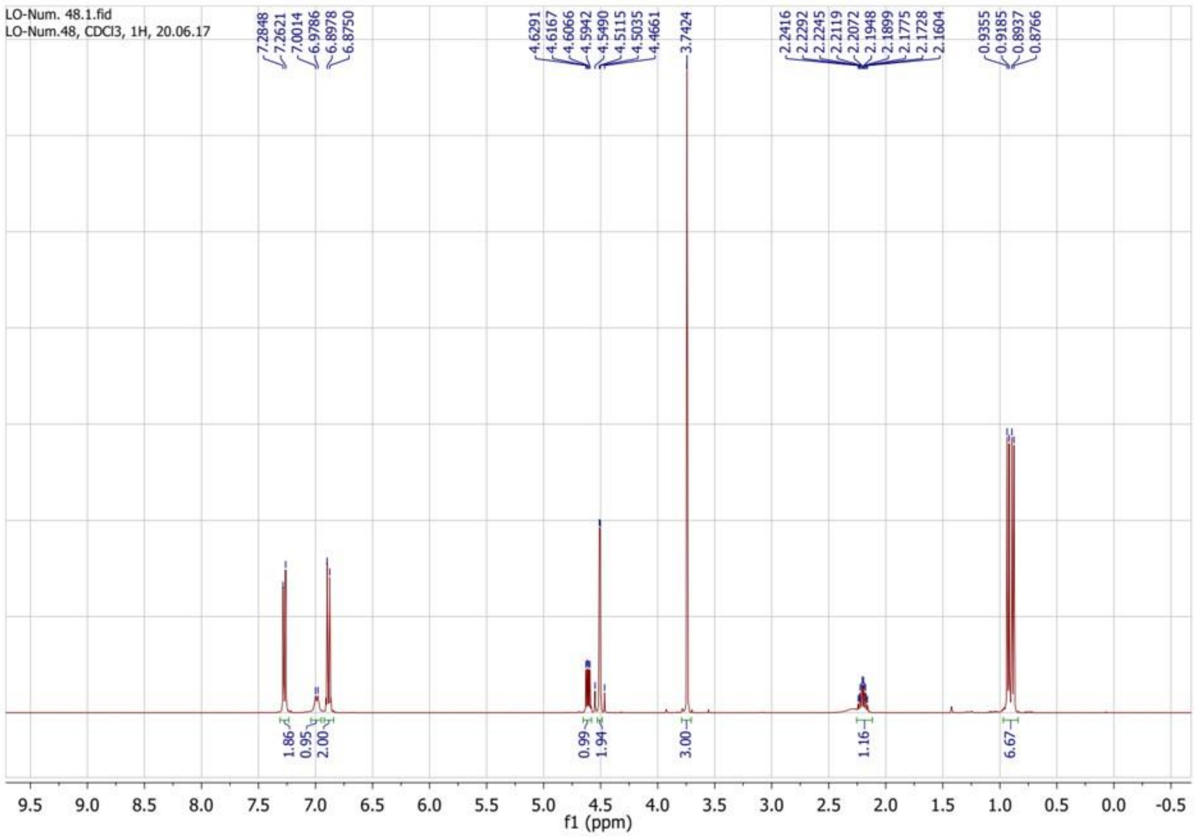


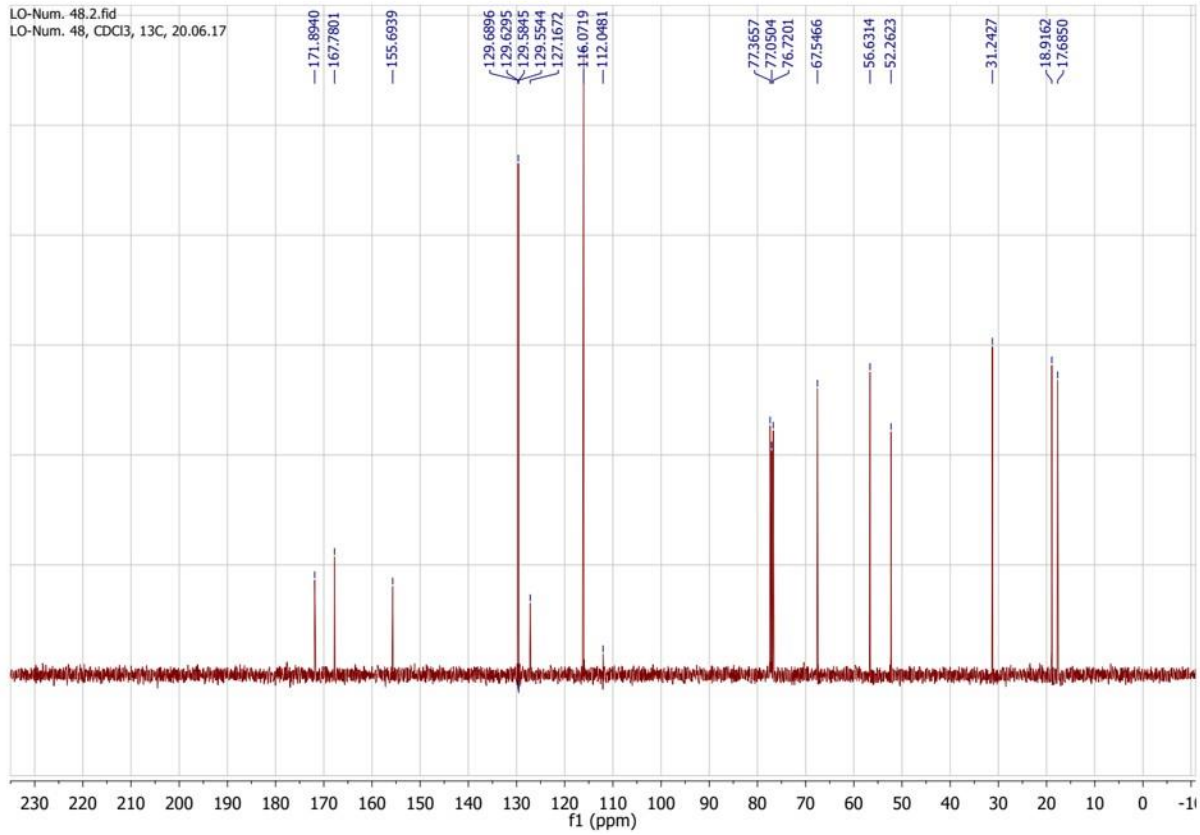


**Figure 62.** $^{1}$H (top) and $^{13}$C (bottom) NMR spectra of compound **1o**.

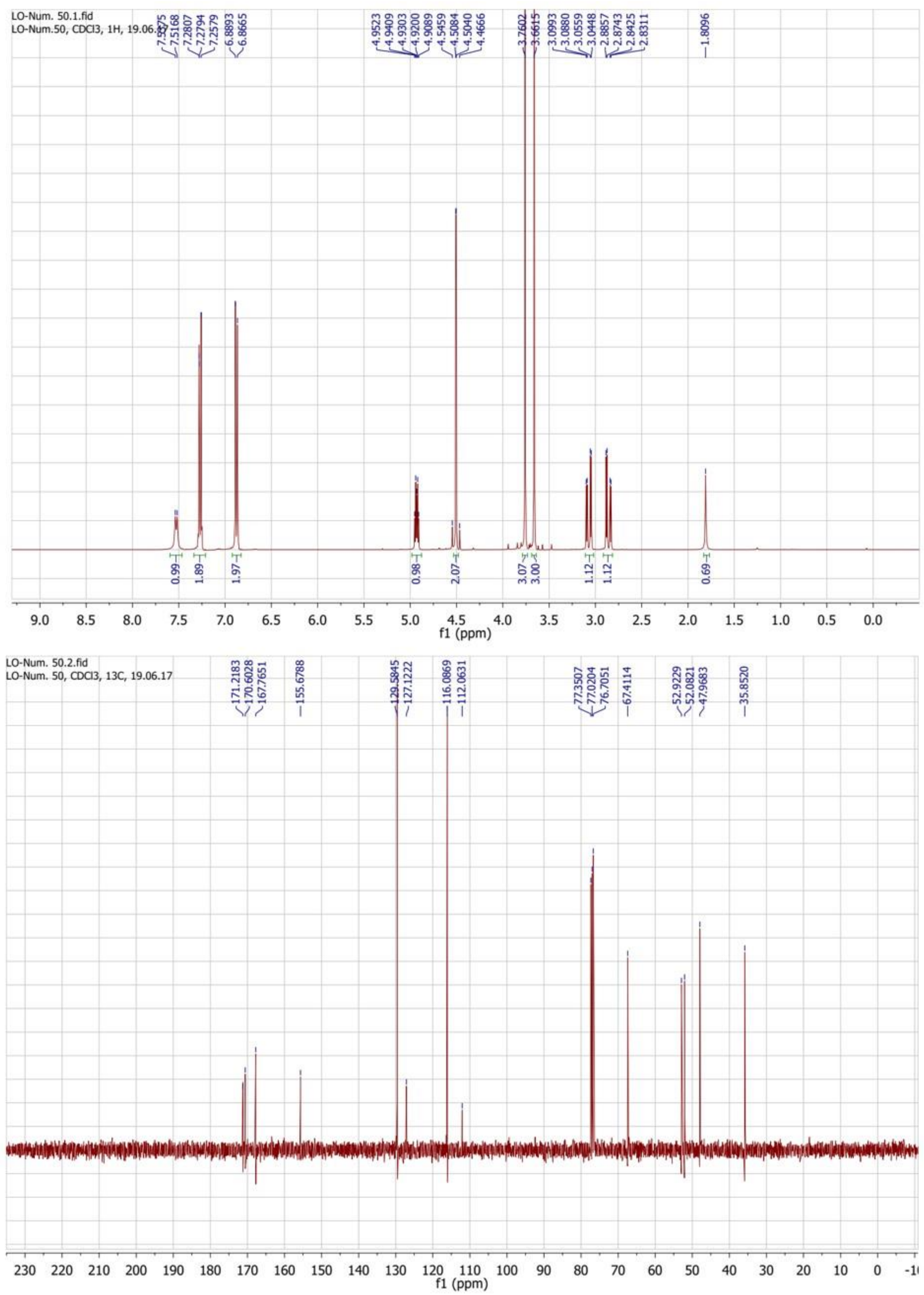


**Figure 63.** $^{1}H$ (top) and $^{13}C$ (bottom) NMR spectra of compound **1p**.

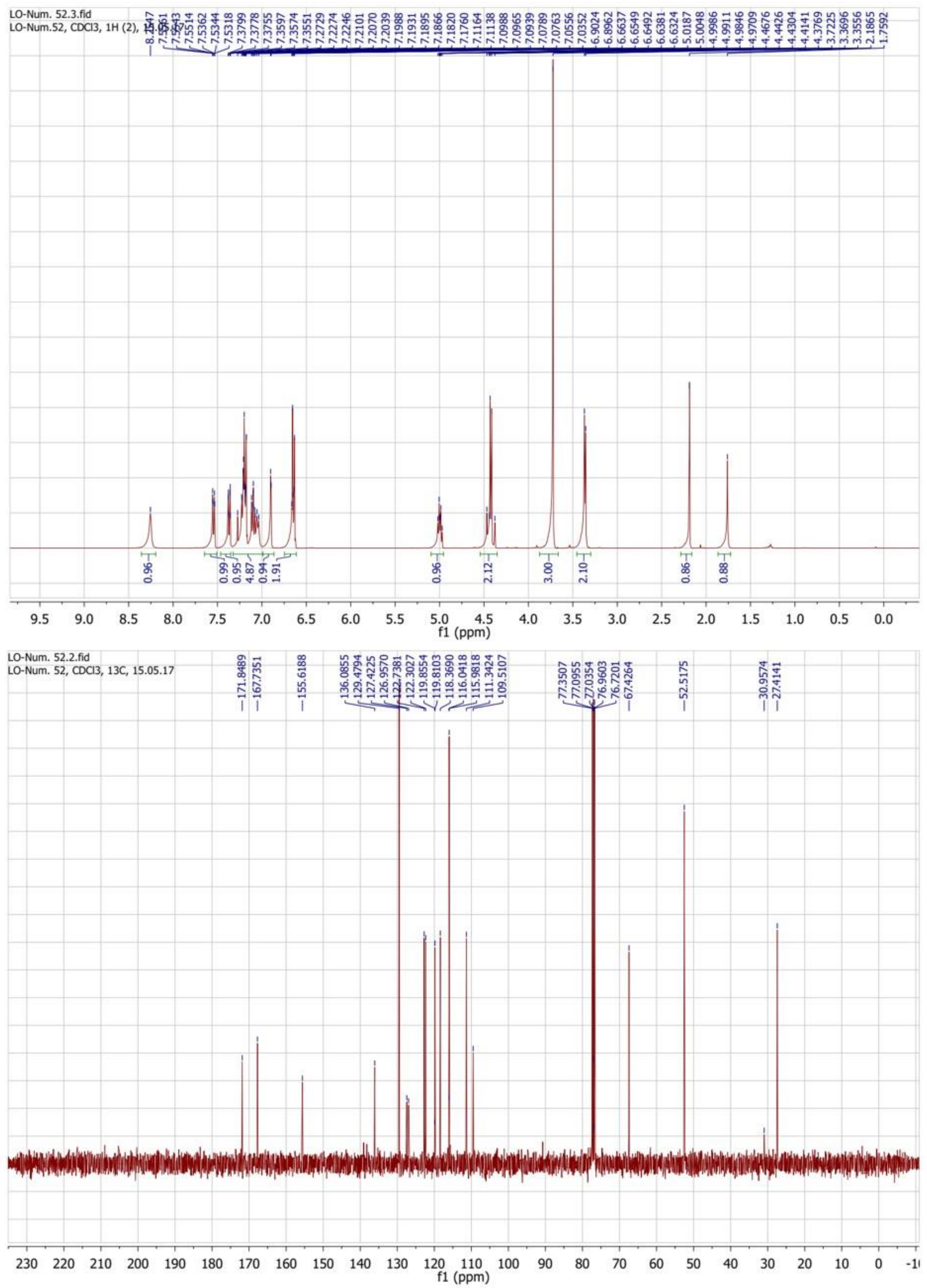


**Figure 64.** $^{1}H$ (top) and $^{13}C$ (bottom) NMR spectra of compound **1q**.

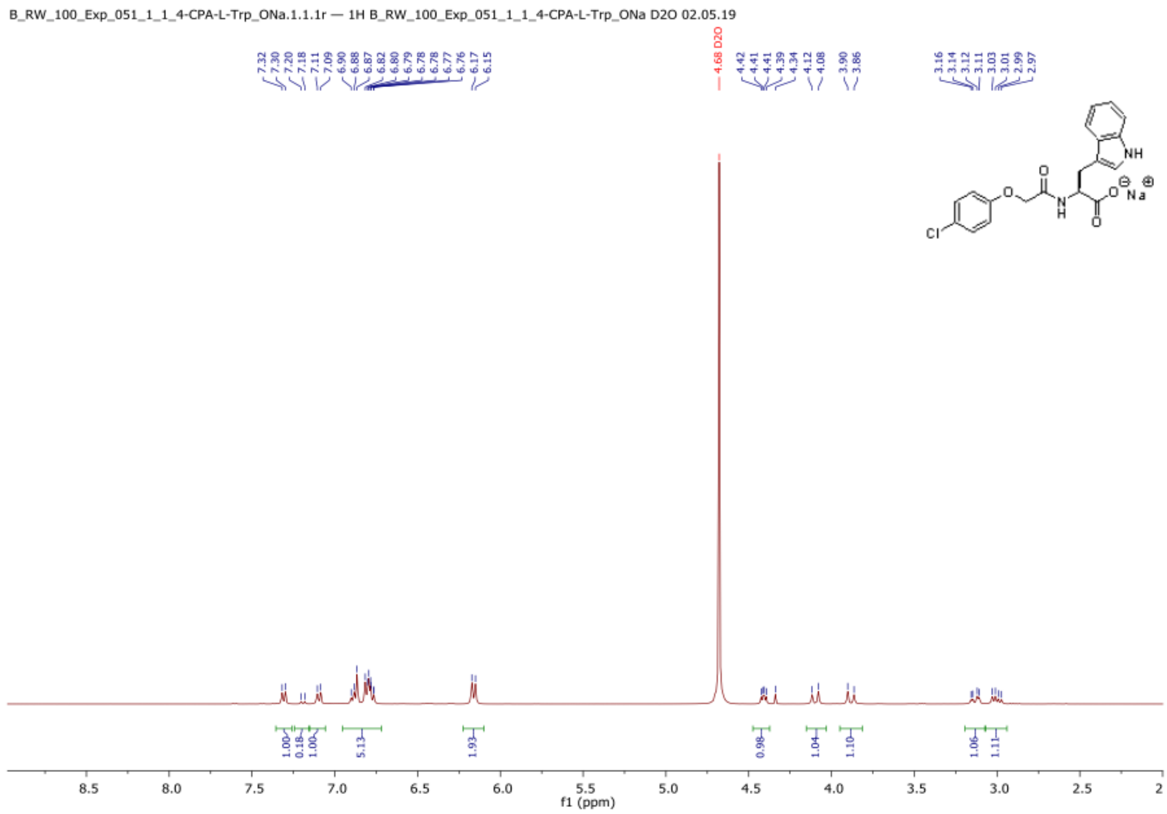

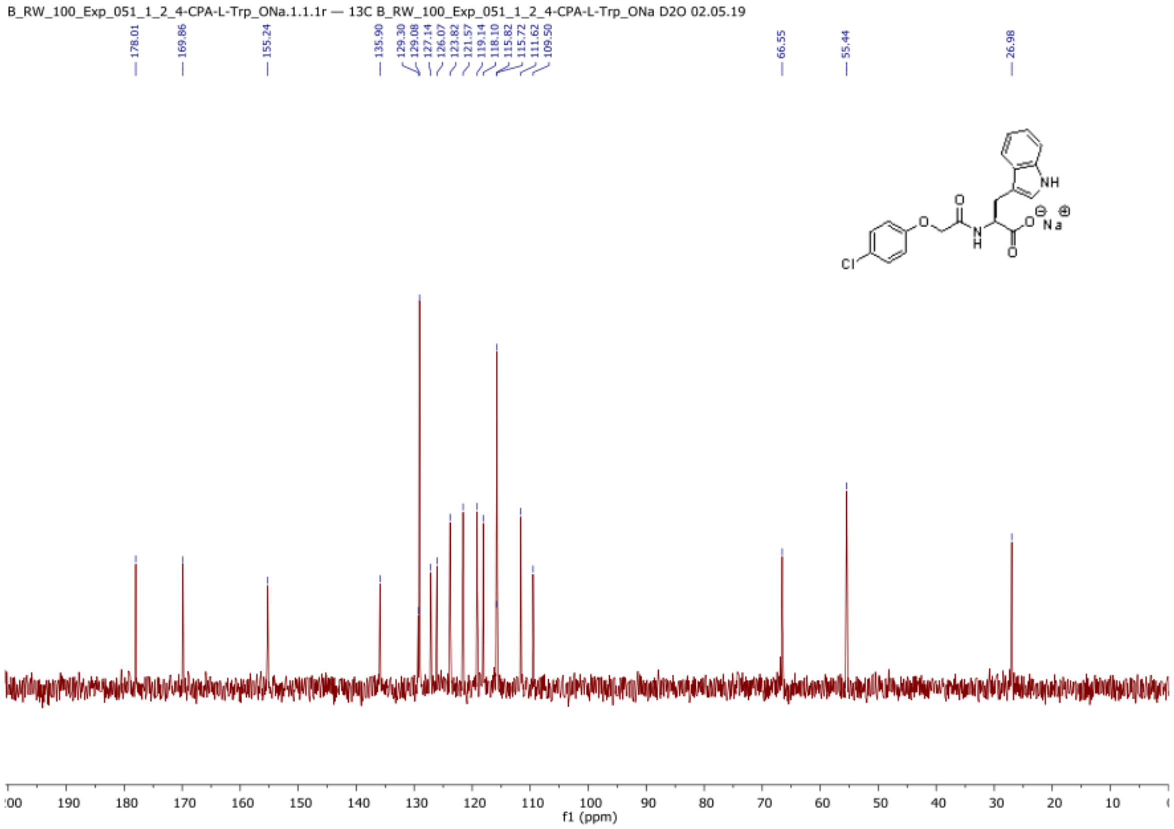

**Figure 65.** $^{1}H$ (top) and $^{13}C$ (bottom) NMR spectra of compound **1r** (in $D_2O$).

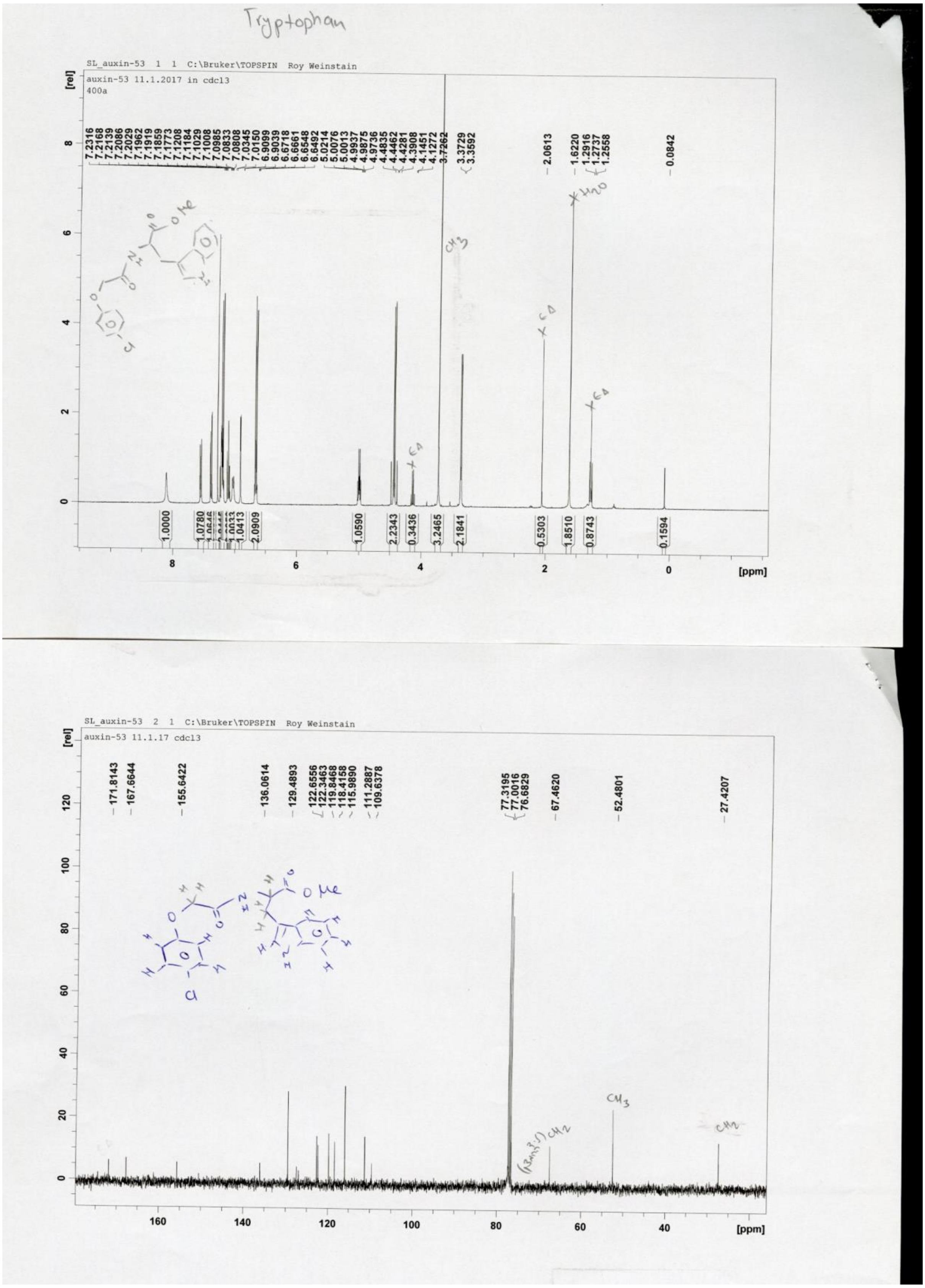


**Figure 66.** $^{1}H$ (top) and $^{13}C$ (bottom) NMR spectra of compound **1s**.

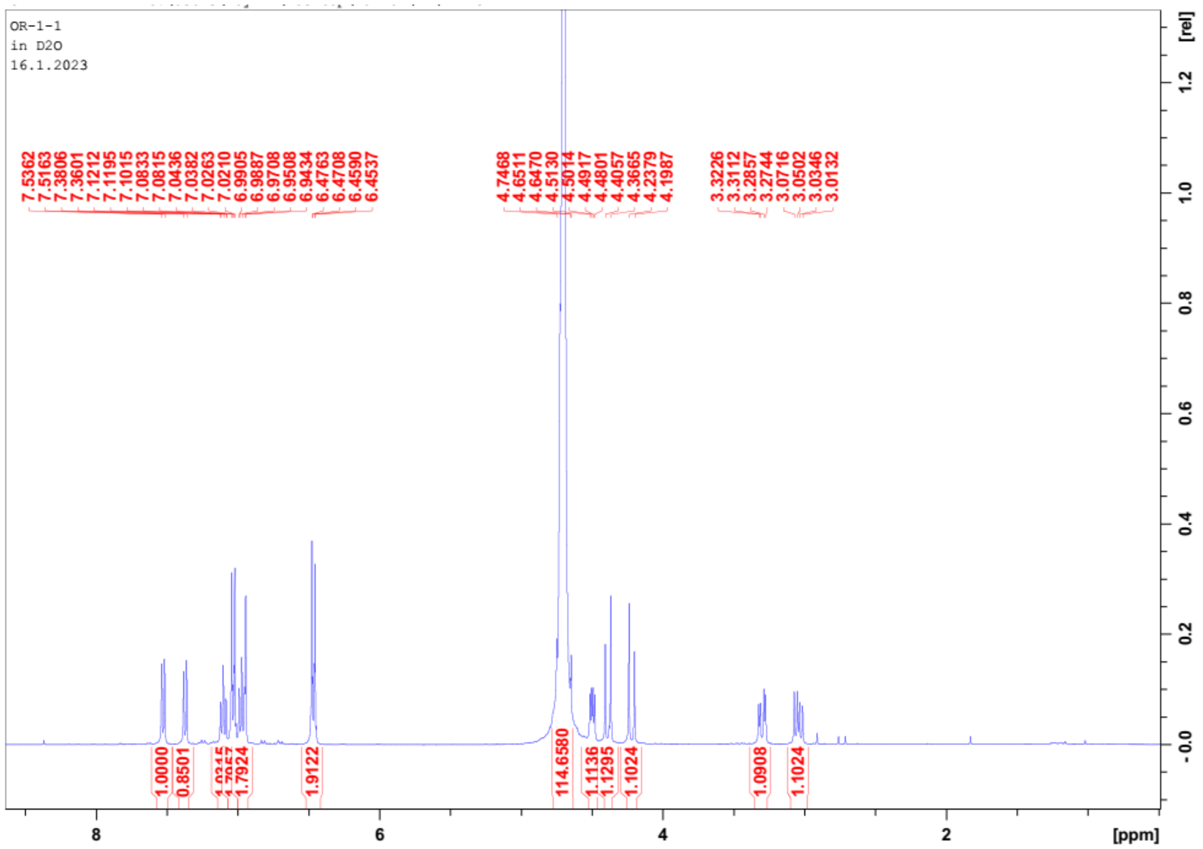


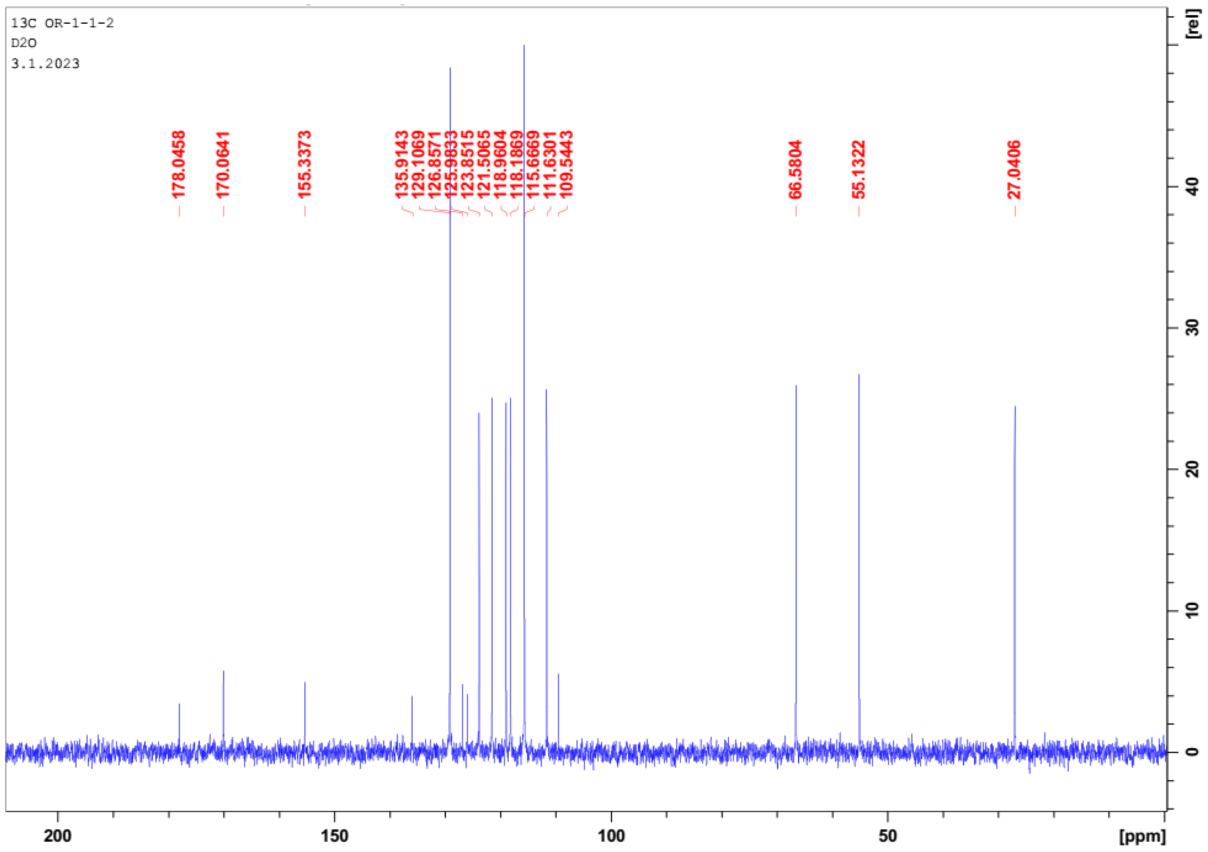


**Figure 67.** $^{1}H$ (top) and $^{13}C$ (bottom) NMR spectra of compound **1t** (in $D_2O$).
